# Oncogenic and tumor-suppressive forces converge on a progenitor niche at the benign-to-malignant transition

**DOI:** 10.1101/2025.06.10.656791

**Authors:** José Reyes, Isabella Del Priore, Andrea C. Chaikovsky, Nikhita Pasnuri, Ahmed M. Elhossiny, Jin Park, Philipp Weiler, Tobias Krause, Andrew Moorman, Catherine Snopkowski, Meril Takizawa, Cassandra Burdziak, Nalin Ratnayeke, Ignas Masilionis, Yu-Jui Ho, Ronan Chaligné, Paul B. Romesser, Aveline Filliol, Tal Nawy, John P. Morris, Zhen Zhao, Marina Pasca Di Magliano, Direna Alonso-Curbelo, Dana Pe’er, Scott W. Lowe

## Abstract

The benign-to-malignant transition is a defining step in cancer progression. To investigate when and how malignancy initiation occurs and tissue reorganization proceeds, we combine single-cell and spatial transcriptomic profiling in mouse models of pancreatic ductal adenocarcinoma (PDAC) that capture spontaneous p53 loss. Among Kras-mutant cells, we find that oncogenic and tumor-suppressive programs, including those controlled by p53, CDKN2A, and SMAD4, are co-activated in a discrete progenitor-like population, engaging senescence-like responses. Using a framework we develop for spatial analysis, we show that a niche centered on these cells undergoes stepwise remodeling during tumor progression, mirroring invasive PDAC. Transient KRAS inhibition depletes progenitor-like cells and dismantles their niche, delaying malignancy initiation. Conversely, p53 suppression enables progenitor cell expansion, epithelial–mesenchymal reprogramming, and immune-privileged niche formation. These findings position the progenitor-like state at the convergence of cancer-driving mutations, plasticity and tissue remodeling, revealing a critical window for intercepting malignancy.

## Introduction

Cancer progression is fueled by the accumulation of genetic alterations, yet clonal expansions bearing oncogenic mutations—despite appearing frequently in histologically normal tissues^1,2^—rarely progress to cancer^3–5^. Only upon acquiring additional genetic or epigenetic changes do they breach regulatory constraints, promoting cellular plasticity, local invasion, and metastasis—features of lethal cancer. How these tissue-level constraints are enforced, and how they ultimately fail, remains poorly understood, owing in part to the difficulty of capturing early events.

Tumor suppressors are physiological mechanisms for cancer detection and interception. In pancreatic ductal adenocarcinoma (PDAC), activating *Kras* mutations drive the formation of transcriptionally diverse premalignant lesions^6,7^, some of which become malignant upon disruption of one or more tumor suppressors, such as *TP53*, *CDKN2A* and *SMAD4*^6,8–12^. Among these, the transcription factor p53 is mutated or lost in 70% of PDAC cases^13^. *TP53* inactivation is rare in low-grade neoplastic lesions^14^, but is more common in high-grade lesions and carcinoma^15^. These findings suggest that p53 directly impairs malignancy—but how and where p53 and other tumor suppressors exert such restraints remains unclear.

Beyond genetics, inflammation is a risk factor for PDAC in humans^16^, and tissue injury accelerates neoplastic progression in mouse models^17^. Furthermore, even in the absence of experimentally induced tissue injury, mutant *Kras* cells can trigger inflammatory programs, implying an inextricable link between oncogenic KRAS activity and tissue remodeling^18–20^.

These events ultimately establish a malignant tumor microenvironment characterized by activated fibroblasts, immunosuppressive myeloid cells, and cytotoxic T-cell exclusion, that shapes tumor progression and therapeutic response^21–25^. Thus, malignant progression not only involves epithelial transformation, but also the coordinated emergence of a tumor-supportive tissue ecosystem. However, we do not understand how genetic lesions, cell state changes, and microenvironmental remodeling act together to trigger the benign-to-malignant switch.

Our prior work revealed that inflammation synergizes with oncogenic KRAS to establish a premalignant subpopulation expressing the pancreatic epithelial progenitor cell marker *Nes*^7,26,27^. *Nes*-positive progenitor-like cells transcriptionally resemble cancer, and exhibit heightened plasticity, permissive chromatin accessibility and cell–cell communication potential^7^. These features suggest that plastic cells have increased propensity to sense and remodel their microenvironment. Notably, the progenitor-like state arises rapidly after injury, whereas the malignant transition takes months^11,28^, suggesting that additional barriers restrain this state and its associated tissue remodeling capability.

Here, we use genetically engineered mouse models to dissect how genetic alterations, cell-state transitions, and microenvironmental remodeling converge to drive early PDAC progression. By reconstructing spatial trajectories of epithelial and stromal remodeling, we show that progenitor-like cells uniquely engage—and are constrained by—tumor-suppressive programs, while assembling a self-reinforcing, cancer-like niche. We show that oncogenic KRAS signaling promotes persistence of this state, whereas p53 enforces its resolution, collapsing the niche and restoring regenerative homeostasis. These findings reveal a plastic cell state and tissue ecosystem that governs the benign-to-malignant transition and highlights a critical window in which malignant progression may be intercepted by targeting the states and niches that enable tumor evolution.

## Results

### Capturing cell states after spontaneous *p53* loss

To investigate the benign-to-malignant transition in pancreatic cancer, we employed the KP^LOH^ model^28^, which marks cells that undergo spontaneous loss of heterozygosity (LOH) of *Trp53* (hereafter, *p53*) during tumor initiation. KP^LOH^ is derived from the KPC mouse (*<u>K</u>ras*^G12D^, *Tr<u>p</u>53*^flox/+^, Ptf1a-<u>C</u>re) and incorporates fluorescent reporters such that *Kras*-mutant pancreatic epithelial cells with wild-type *p53* are mKate2+/GFP+, whereas cells with *p53* LOH become mKate2+/GFP-due to co-deletion of a physically linked GFP reporter (**Figure 1A**). In mice lacking macroscopic PDAC (3–4 months old, termed “pre-tumor” stage), these *p53*-deficient (mKate2+/GFP-) cells appeared as rare, isolated cells, small clusters, or microscopic tumors histologically resembling PDAC^28^ (**Figure 1B,C**). Thus the KP^LOH^ model enables interrogation of the consequences of *p53* loss before detectable PDAC formation.

**Figure 1.**
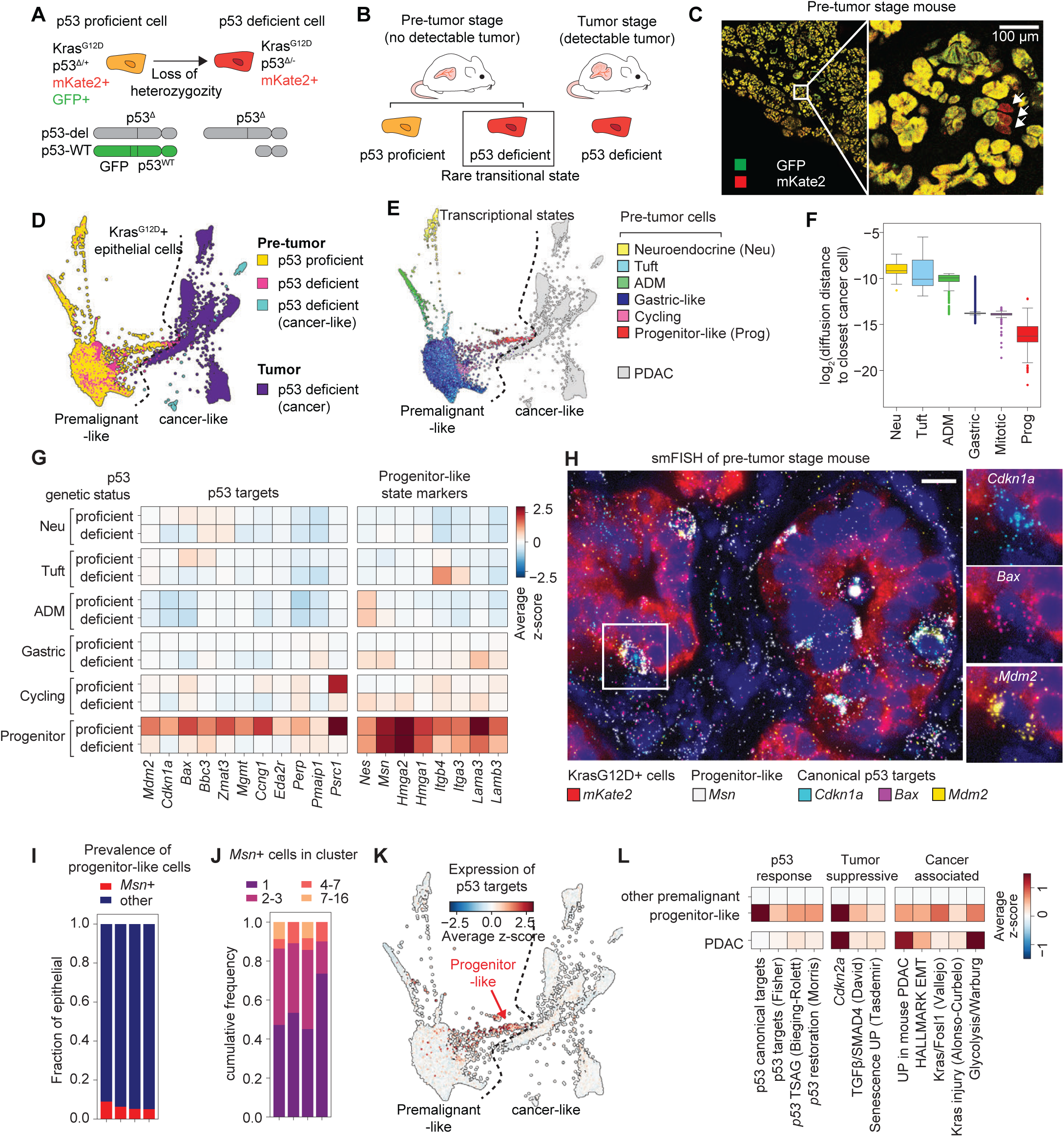
Capturing spontaneous loss of *p53* throughout the premalignant-to-malignant spectrum. **A.** KP^LOH^ mouse model. Loss of GFP linked to the only wild-type *p53* copy in the cell reports *p53* LOH. **B.** Sampling strategy to characterize premalignant-to-malignant progression. **C.** Representative fluorescence image of a pre-tumor stage pancreas section. Arrowheads, rare cells that lost GFP fluorescence upon *p53* LOH. **D.** Force-directed layout (FDL) of single-cell transcriptional data from sorted *Kras*^G12D^ epithelial cells, colored by mouse stage and *p53* status. **E.** FDL of cells in (D), colored by transcriptional signatures of major premalignant subpopulations (Methods). ADM, acinar-to-ductal metaplasia. ‘Progenitor-like’ denotes *Nes*^27^ progenitor marker expression, not the progenitor PDAC subtype from Bailey et al.^101^ **F.** Diffusion distance from pre-tumor *p53*-proficient cells to the closest cancer-like cell. **G.** Expression of known p53 targets and markers of progenitor-like cells in pre-tumor *p53*-proficient and deficient cells, as a function of cell state. **H.** Representative smFISH image of pre-tumor pancreas, showing colocalization of p53 targets and progenitor state marker *Msn*. Scale bar, 10μm. **I.** Fraction of Msn+ cells in premalignant pancreas from pre-tumor stage KP^LOH^ mice. Each bar represents one mouse. **J.** Sizes of progenitor-like cell clusters in four pre-tumor stage KP^LOH^ mice. **K.** FDL of *Kras*^G12D^-positive epithelial cells along PDAC progression, colored by average expression of p53 canonical targets shown in (G). **L.** Expression of tumor-suppressive and oncogenic gene signatures in pre-tumor *p53*-proficient cells (other premalignant and progenitor-like) or tumor *p53*-deficient cells (PDAC). Signatures include: p53 canonical targets in (G); p53 curated targets (Fisher)^102^; p53 tumor suppression–associated genes (TSAG)^103^; p53-restoration^64^; *Cdkn2a* mRNA; TGFβ-dependent SMAD4 targets^32^; HALLMARK EMT, epithelial-to-mesenchymal transition^104^; senescence UP^105^; UP in mouse PDAC (this work; Methods); Kras/Fosl1^34^; Kras injury^26^; glycolysis/Warburg (curated list; Methods).

Using single-cell RNA sequencing (scRNA-seq), we compared rare single-positive cells that underwent *p53* LOH (pre-tumor *p53*-deficient, 1–3% of all mKate2+ cells) to their double-positive counterparts (pre-tumor *p53*-proficient), and to PDAC-derived single-positive cells (tumor *p53*-deficient) (Methods; **Figures 1D** and **S1A,B**). Consistent with prior work, *p53*-proficient premalignant cells occupied heterogeneous transcriptional states that were distinct from both normal acinar and PDAC phenotypes^7,29^ (**Figures 1E** and **S1A–F**). The *p53*-proficient fraction included cells expressing markers of acinar-to-ductal metaplasia (ADM) (*Cpa1, Krt19*), neuroendocrine fate (*Scg5, Chga, Chgb*), tuft fate (*Pou2f3*), proliferation (*Mki67*, *Cdk1*), and gastric fates characteristic of PanIN lesions and the classical PDAC subtype (*Dmbt1, Muc6, Tff1, Tff2, Anxa10*).

Notably, a small subset of premalignant cells expressed the progenitor-like program, an injury-responsive cancer-like cell state transiently induced in Kras^G12D^-expressing epithelium (e.g., *Nes*, *Msn, Hmga2, Hmga1, Vim*)^7^ (**Figures 1E** and **S1D–F**). This subpopulation increased in frequency in *p53-*proficient cells whenever PDAC was also present, implying tissue-level microenvironmental influences (**Figure S1F**) to their emergence or maintenance. Diffusion distance analysis (Methods) revealed that among all pre-tumor *p53-*proficient cells, the progenitor-like population is transcriptionally closest to PDAC (**Figure 1F**), nominating this rare and highly plastic population as a likely intermediate in the benign-to-malignant transition.

We previously showed that *p53* loss is followed by a progressive accumulation of select copy number alterations (CNAs)^28^. Reasoning that this may provide a “timestamp” of a cell’s trajectory toward malignancy, we inferred CNAs from scRNA-seq data. We found that most pre-tumor *p53*-deficient cells had quiet genomes (aside from chr11 loss) and occupied premalignant transcriptional states, indicating that they underwent *p53* LOH but had not yet acquired genomic instability or malignant phenotypes (**Figure S1G,H**). Conversely, some *p53*-deficient cells displayed highly rearranged genomes and transcriptional profiles resembling PDAC. Such “microtumors” are likely clonal expansions of malignant cells that cannot be detected by ultrasound or gross pathology^28^, and also express progenitor state marker HMGA2 (**Figure S1I,J**), suggesting a link between malignant transformation and progenitor-like program engagement.

Some pre-tumor *p53*-deficient cells exhibited intermediate CNA levels, consistent with a transitional state experiencing *p53* LOH but lacking additional genetic alterations required for full malignancy. Many of these cells occupied the progenitor-like state and shared some distinguishing karyotypic changes with highly rearranged, malignant-appearing cells from the same mouse—for example, harboring loss of chr4 (*Cdkn2a*) and chr11 (*p53*) and gain of chr2, while retaining diploid status in other chromosomes altered in microtumor cells from the same sample (e.g., chr5, chr6, chr10, chr13, chr14) (**Figure S1K,L**). Together, these data point to the highly plastic, progenitor-like state, as a likely precursor for malignant tumors.

### Rare progenitor-like cells exhibit peak oncogenic and tumor suppressive activity

Our single-cell data defined a window for interrogating the molecular events that precede or immediately follow *p53* inactivation during early tumorigenesis. To identify the effect of spontaneous *p53* loss on different premalignant subpopulations, we compared the expression of canonical p53 targets between *p53*-proficient and *p53*-deficient cells within each transcriptionally defined premalignant state. Surprisingly, most premalignant epithelial states experienced little impact, with one notable exception—progenitor-like cells (Methods; **Figure 1G**). In *p53*-proficient contexts, this subpopulation expressed the highest levels of p53 targets associated with cell cycle arrest (e.g., *Cdkn1a, Ccng1*), DNA repair (e.g., *Mgmt*) and apoptosis (e.g., *Bbc3, Bax*), which were downregulated following *p53* loss, confirming their p53 dependence. Single molecule fluorescence in situ hybridization (smFISH) revealed that individual *Msn*-positive progenitor-like cells were dispersed throughout glandular structures in the premalignant pancreas, suggesting that they emerge recurrently and independently during tumorigenesis (**Figure 1H–J**). Thus, despite uniform *Kras^G12D^*mutation across the epithelial compartment and loss of acinar identity^7^, p53 activity is selectively confined to progenitor-like cells—the subpopulation most transcriptionally similar to PDAC (**Figure 1K**).

Strikingly, progenitor-like cells also exhibited the highest engagement of the other major tumor suppressive programs in PDAC^13^: CDKN2A^30^ and SMAD4^31^ (**Figures 1L** and **S1M,N**). Progenitor-like cells significantly upregulated *Cdkn2a* relative to other premalignant cells. Furthermore, inspection of splice junctions revealed engagement of both the p19^ARF^ and p16^INK4A^ isoforms encoded by the *Cdkn2a* locus (**Figure S1O**), indicating that both tumor suppressive programs were engaged in premalignant cells^10,30^. In addition, progenitor-like cells selectively elevated the TGFβ pathway (**Figure S1M**), which was confirmed at the level of individual SMAD4-dependent, TGFβ-induced genes^32^ (**Figure S1N**). Thus, all three tumor suppressive programs that are commonly lost during PDAC progression are concurrently engaged in the progenitor-like state.

Regardless of *p53* status, progenitor-like cells selectively upregulated gene expression programs associated with malignant PDAC, including RAS signaling, glycolysis, and epithelial–mesenchymal transition (EMT)^26,33–35^ (**Figures 1L** and **S1M,N**). The simultaneous engagement of tumor-promoting and tumor-suppressive pathways in the same cells is reminiscent of oncogene-induced senescence, a potent p53 and p16^INK4a^-dependent tumor suppressive program triggered by aberrant RAS signaling^36–39^. Indeed, progenitor-like cells were enriched for senescence-associated transcriptional signatures (**Figures 1L** and **S1M,N**). Thus, our results position progenitor-like cells as the premalignant subpopulation in which the critical antagonism between oncogenic and tumor suppressive programs takes place during PDAC initiation.

### Adoption of progenitor-like identity is coupled with tissue reorganization

We previously showed that progenitor-like cells are a highly plastic subpopulation with elevated cell–cell communication potential that expands following caerulein-induced pancreatitis in *Kras*-mutant mice^7,26^. These cells are rare in 12–27-week-old uninjured mice, but accumulate to 60% of the epithelium upon acute pancreatitis, before gradually declining over 3 weeks (**Figure 2A,B**)^7^. Accumulation of progenitor-like cells coincides with broader tissue events, including loss of acinar identity and formation of a fibrotic stromal niche, linking injury-induced progenitor expansion to coordinated tissue remodeling during early premalignant progression.

**Figure 2.**
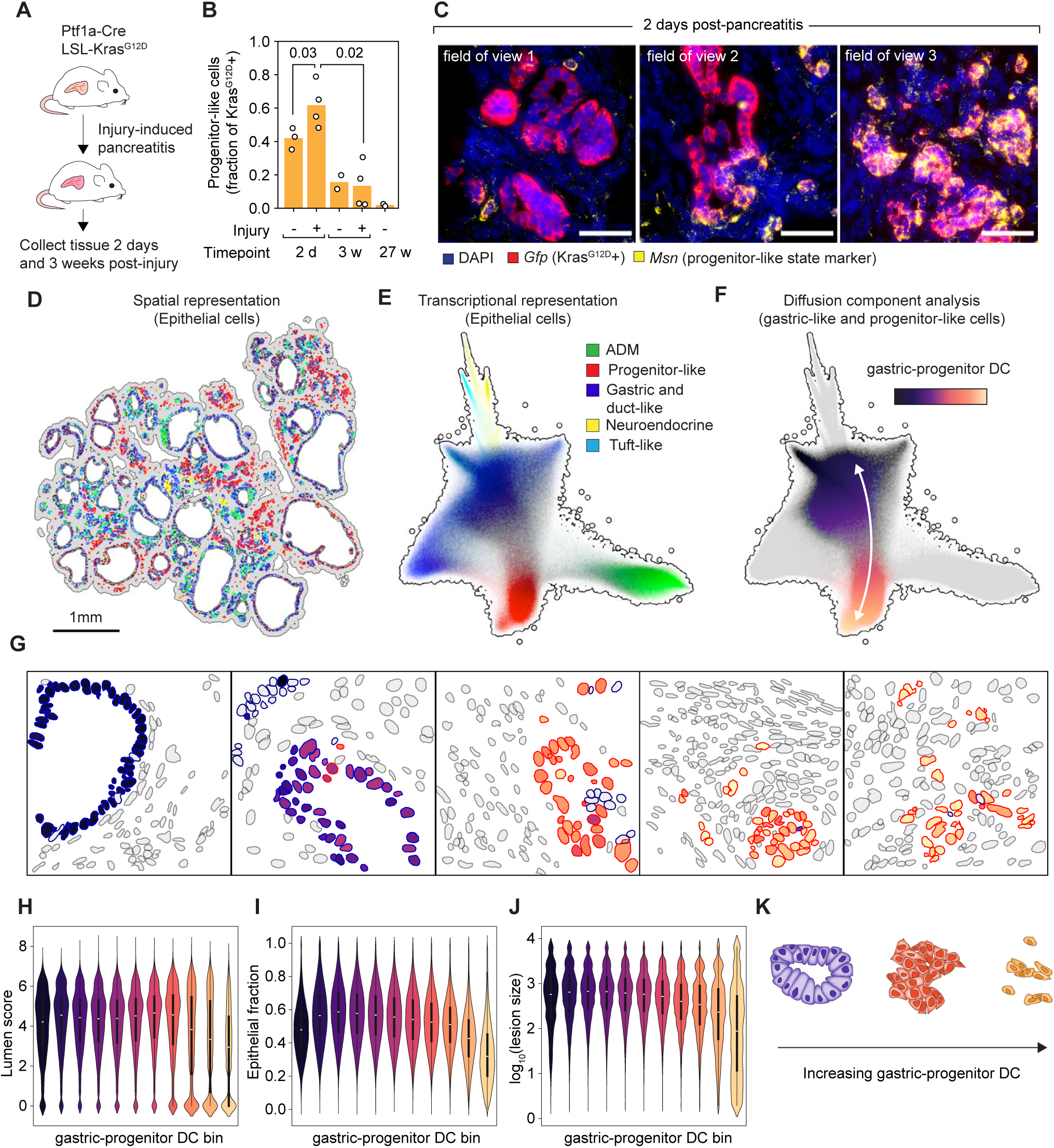
Transcriptional and morphological states undergo coordinated changes in the premalignant epithelium. **A.** Experimental timeline for tissue collection after inducing pancreatitis with caerulein in KC mice. **B.** Fraction of progenitor-like epithelial cells in scRNA-seq data^7^ as a function of injury and time. *p*-values, two-tailed Wilcoxon rank-sums test. **C.** Representative smFISH images from pancreata 2-days post-injury. The three fields of view are from the same tissue. Scale bars, 50 μm. **D,E.** Spatial representation (D) or single-cell transcriptional embedding (E) of Xenium data annotated by signatures of major premalignant subpopulations. **F.** Embedding in (E), colored by gastric–progenitor diffusion component. **G.** Representative fields of view of premalignant epithelial lesions in Xenium data. Segmented nuclei are pseudo-colored by their gastric–progenitor DC value, using the colormap in (F). **H–J.** Lumen score (H), epithelial fraction in local spatial neighborhood (I) and lesion size (J) as a function of gastric–progenitor DC in epithelial cells (see Methods for morphological parameter quantification). **K.** Schematic of lesion morphologies along the gastric–progenitor DC.

To understand how epithelial plasticity unfolds at tissue scale, we leveraged multiallelic mESCs to generate large, synchronous cohorts of KC (LSL-*Kras*^G12D^; *Ptf1a*-Cre) mice^40,41^ in which GFP expression serves as a proxy for mutant *Kras* activation. We used caerulein to induce pancreatitis, and harvested pancreata 2 days or 3 weeks after treament to capture progenitor-like cell dynamics (**Figure 2A,B**). Consistent with dissociated single-cell analyses^7^, progenitor-like cells accumulated within 48 hours of injury-induced pancreatitis, as evidenced by upregulation of MSN, HMGA2, and the tumor suppressors p53 and p19^ARF^ (**Figure S2A–C**).

Lesions enriched with progenitor-like cells (“progenitor lesions”) formed disorganized structures that contrasted with the rosette-like and luminal morphology of premalignant epithelium lacking these cells (**Figures 2C** and **S2A**). Within the same mouse, progenitor-like cells appeared as small, isolated clusters; as nests or cords of cells penetrating stroma; or as mixed glandular and solid lesions—suggesting snapshots of progressive epithelial identity loss during the earliest stages of RAS-driven transformation (**Figure 2C**).

To further characterize progenitor lesion emergence, we performed spatial transcriptomics using the Xenium In Situ platform (10x Genomics). Guided by prior scRNA-seq data^7,29^ in the premalignant pancreas (**Data S1**), we designed a 480-gene panel with markers for resolving premalignant heterogeneity, capturing stromal and immune states, and probing key signaling pathways^7^ in early tumorigenesis (Methods; **Data S2**). We spatially profiled KC mouse tissue harvested 1–2 days (*n* = 10 mice) and 3 weeks (*n* = 5 mice) following caerulein-induced injury, constituting over 3.5 million cells across a continuum of premalignant lesions.

Our computational framework represented each cell’s phenotype as its full transcriptional state, rather than collapsing it to a predefined cell type, and treated spatial and transcriptional data as complementary, integrated dimensions. This dual representation allowed us to analyze continuous phenotypic variation within intact tissue using UMAP embedding and diffusion analysis, while directly relating transcriptional state to spatial location (Methods; **Figure 2D–F**). Our analysis revealed a rich diversity of cell states and spatial organization across multiple length scales—from macroscopic tissue landmarks (e.g. lymphatic channels and lobular architecture) to microscopic structures (e.g, epithelial lesions, ducts and vasculature), down to fine-grained gradients (e.g. fibroblast layering) (**Figure S2D**).

In the epithelial compartment, we recovered major subpopulations identified by scRNA-seq^7,29^ (Methods; **Figures 1E** and **2D,E**) and revealed continuous transcriptional gradients in phenotypic space, consistent with dynamic transitions between premalignant epithelial subpopulations (**Figure 2E**). Diffusion component analysis^42,43^ identified the continuous axis connecting gastric-like and progenitor-like states as the dominant axis of transcriptional variation (hereby, gastric–progenitor DC) (Methods; **Figure 2F**). This axis corresponded to a trajectory in transcriptional state, characterized by the sequential induction of *Msn*, then *Hmga2* and finally mesenchymal marker *Vim* (**Figure S2E**).

The gastric–progenitor DC provided a quantitative framework for assessing how epithelial cells acquire progenitor-like features and remodel their local microenvironments. Ordering epithelial cells along this axis and analyzing the geometric properties of their corresponding lesions (Methods) revealed progressive loss of luminal architecture, reduced epithelial density accompanied by immune and stromal infiltration, and eventual collapse of lesions into isolated progenitor-like cells embedded within stroma (**Figure 2G–K**). These findings demonstrate that epithelial cells undergoing progenitor-like reprogramming progressively lose epithelial organization and gain mesenchymal traits^44^ (**Figure S2F**), defining a continuum of premalignant remodeling in which transcriptional plasticity, epithelial architecture, and local tissue context are tightly coupled.

### Niche pseudotime reconstructs assembly of a PDAC-like progenitor niche

PDAC exists within a wound-repair–like microenvironment dominated by activated fibroblasts and immunosuppressive myeloid populations that restrict cytotoxic T-cell infiltration and limit therapeutic efficacy(^21–25^. To reconstruct the dynamics of microenvironmental remodeling during tumor initiation, we leveraged the asynchronous nature of tissue remodeling captured in our spatial datasets. Tissues contained epithelial cells spanning the gastric–progenitor (G–P) continuum, surrounded by diverse stromal and immune populations (**Figure S2D,E**). Within a given spatial neighborhood, epithelial cells typically occupied similar positions along the G–P axis, forming coherent regional patterns. We formalized these spatial neighborhoods, or niches, as all cells within a 60-μm radius of an epithelial cell ‘anchor’ (**Figure 3A,B**), treating these as the fundamental units of premalignant progression.

**Figure 3.**
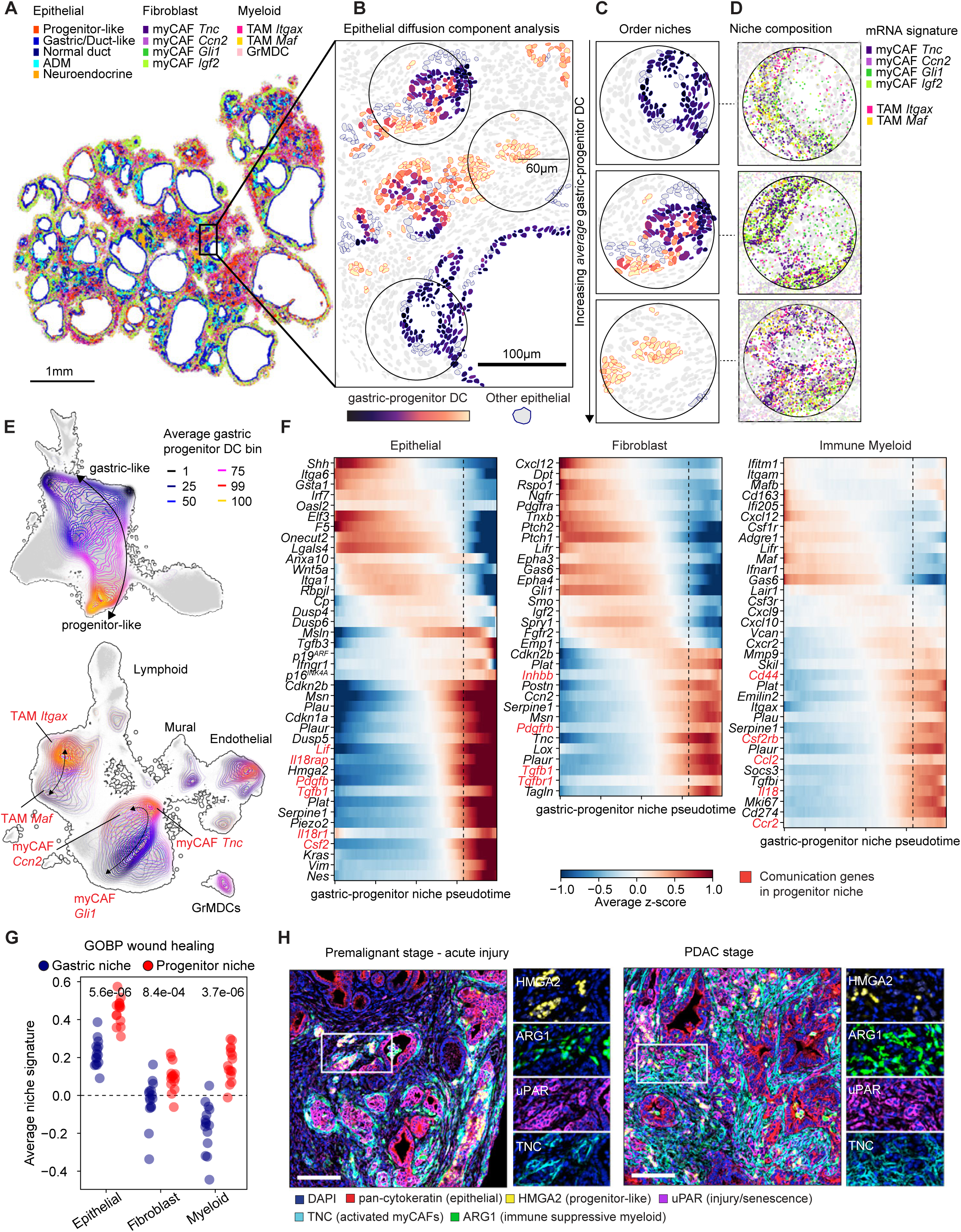
Progenitor niche assembly proceeds via continuous cellular and molecular remodeling events. **A.** Representative Xenium data from premalignant pancreas harvested 2 days post-pancreatitis, colored by cell type. **B.** Projection of gastric–progenitor DC in premalignant cells. Epithelial cells not categorized as gastric-like or progenitor-like are gray with dark blue outline. **C.** Niches, comprising all cells within a 60-μm radius of a central anchor epithelial cell, are ordered by the average gastric–progenitor DC of their constituent epithelial cells. **D.** Location of individual mRNA molecules associated with select myofibroblasts and monocyte/macrophage subpopulations in niches depicted in (C). **E.** Contour plots denoting the binned density of niche epithelial cells along the average gastric–progenitor DC (top) and corresponding shifts in the density of microenvironment cells (bottom). **F.** Average niche expression of select genes in different compartments along niches ordered by average gastric–progenitor DC value (Methods). Dotted lines indicate DC value at which epithelial cells begin expressing progenitor-like markers. Communication genes associated with progenitor niches are highlighted in red. **G.** Average expression of wound-healing response genes (GO Biological Processes) from our Xenium panel. Each dot represents a biological replicate (*n* = 15 mice). Values denote the average z-scored expression of wound-healing genes in gastric or progenitor niches of the specified cellular compartments. p-value, two-tailed Wilcoxon rank sum test. **H.** Immunofluorescence staining for cellular states enriched in the progenitor niche, in premalignant and malignant mouse samples. Scale bars, 100 μm.

To order niches along premalignant progression, we defined a niche pseudotime based on gene expression, driven exclusively by epithelial cells. Analogous to single-cell trajectory inference^42,45,46^, this approach reconstructs a pseudotemporal ordering of niches by averaging the G–P diffusion coordinate of epithelial cells within each niche, yielding a consensus epithelial transcriptional state (**Figure 3C**). We refer to this epithelial-defined ordering as the G–P niche pseudotime. This framework allowed us to connect the gradual adoption of an epithelial progenitor-like transcriptional identity with progressive microenvironmental remodeling, revealing niche dynamics from static tissue snapshots (**Figure 3D**).

To resolve how the microenvironment changes along G–P niche pseudotime, we embedded all non-epithelial cells in transcriptional UMAP space and visualized their densities as a function of G–P niche pseudotime in their respective niche (**Figure 3E**). This continuous, state-based framework revealed gradual and coordinated remodeling of the fibroblast and myeloid compartments that would be missed by analyzing niche composition at the level of coarse cell types (**Figure S2G**). Whereas niches dominated by gastric-like epithelial cells were surrounded by SHH-responsive *Gli1*+ myofibroblasts^47^ and *Maf+*^48^ myeloid cells, we observed progressive enrichment of *Itgax+* monocyte/macrophages and myofibroblasts expressing ECM and TGFβ-related genes (e.g., *Postn*, *Tgfb1*, *Tnc*) as niches became dominated by progenitor-like cells (**Figures 3E,F** and **S2H**). G–P niche pseudotime culminated with the formation of cellular communities characterized by progenitor-like cells with advanced mesenchymal features, surrounded by myofibroblasts expressing *Ccn1* and *Ccn2* (known to be induced by YAP signaling), TGFβ signaling components and hypoxia genes, among other forms of stress^49^. Niche remodeling is thus a progressive, spatially organized process tightly coupled to epithelial dedifferentiation.

Shifts along the G–P niche continuum involved gradual, compartment-specific gene expression changes (**Figure 3F**) that were robust to the choice of niche radius (30 μm to 200 μm; Methods; **Figure S2I**), reinforcing the progressive rather than binary nature of niche remodeling. Analysis of dissociated single-cell datasets from injured KC mice revealed that the dominant axes of variation in fibroblast and myeloid cell transcriptomes mirrored the niche trajectory (**Figures 3F** and **S2J**), despite not contributing to the definition of G–P niche pseudotime, indicating that spatial context is a major source of transcriptional heterogeneity in the premalignant pancreas. This analysis also revealed that myeloid cells in progenitor-associated niches progressively expressed transcriptional programs characteristic of immune suppressive subpopulations (*Spp1*, *Arg1* and *Il1b*)^50–52^, while fibroblasts upregulated activation markers (*Acta2, Timp1, Tgfb1, Tnc*) and acquired features of senescent myofibroblasts (*Cdkn2a, Cdkn2b, Plaur*) previously linked to PDAC progression^53^.

Given that progenitor-like cells display features of senescence (**Figure S1M,N**), a state implicated in tissue repair and fibrosis^54,55^, we hypothesized that their emergence reflects an aberrant wound-healing response co-opted by oncogenic KRAS. Consistent with this, multiple compartments in the progenitor niche induced a wound-healing response signature (**Figure 3G**), including mediators of inflammation and fibrosis^56,57^ (e.g. *Pgdfb* and *Tnfrsf12a*) and the plasminogen pathway (e.g. *Plaur*, *Plat*, *Plau*, *Serpine1*) (**Data S1**), as well as staining for the senescence and injury-related protein uPAR (**Figure S2K,L**). These findings suggest that progenitor-like cells initiate a conserved, multi-lineage wound-healing program that, when sustained by oncogenic signaling, promotes the assembly of tumor-permissive niches.

Together, these spatially resolved trajectories reconstruct an ordered tissue remodeling program that culminates in the formation of the progenitor niche—a multicellular community defined by the presence of ARG1+ macrophages and TNC+ myofibroblasts. This niche recapitulates desmoplastic, immunosuppressive microenvironment of malignant PDAC in patients^21^ and mouse models (**Figure 3H**), yet arises in the premalignant setting, consistent with progenitor-like cells acting as early architects of a cancer-like niche.

### Rare communities in human pancreatitis resemble the progenitor niche

To determine whether a state analogous to progenitor-like cells exists in non-transformed human pancreas, we reanalyzed scRNA-seq data from pancreatic epithelial cells obtained at warm autopsy from cancer-free individuals^2^. Projection of murine progenitor-like signatures onto these data revealed a rare subset of ADM- and duct-like cells across multiple donors expressing progenitor-state markers *MSN* and *HMGA1*, as well as markers of KRAS activation and tumor suppressive responses, including senescence-related programs (**Figures 4A,B** and **S3A,B**).

**Figure 4.**
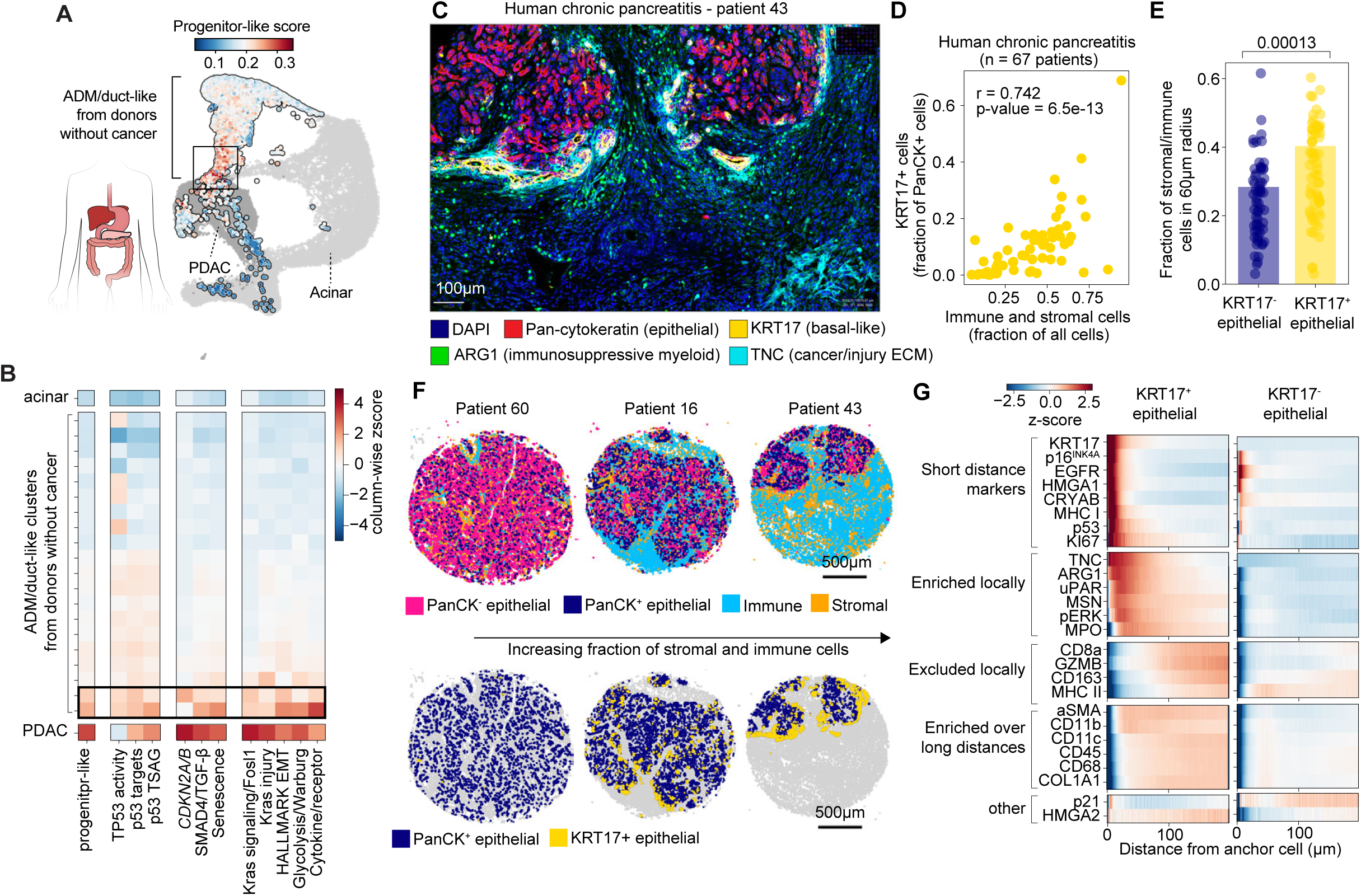
Features of the progenitor niche are present in human chronic pancreatitis samples. **A.** UMAP of cells from cancer-free human pancreas and PDAC tissue in ref.^2^, colored by progenitor-like signature. PDAC cells and acinar cells are grayed out. Box highlights cells that exhibit highest progenitor-like signatures. **B.** Expression of tumor-suppressive and oncogenic signatures in pancreatic epithelial cells from donors with and without cancer. Each row of ADM/duct-like cells corresponds to a PhenoGraph cluster. Colors represent z-score of average signature scores in each column. Box highlights the two PhenoGraph clusters with highest progenitor-like signatures in cells from donors without cancer. **C.** Representative multiplex immunofluorescence image of human pancreatitis sample, visualizing KRT17+ epithelial cells and their associated niches. **D.** Fraction of epithelial cells that are KRT17+, as a function of the fraction of stromal and immune cells in individual patient samples. p-value, Spearman’s rank correlation. **E.** Niche composition of KRT17+ or KRT17-epithelial cells. Each dot is a single patient. Samples with fewer than 15 KRT17+ cells (*n* = 6 patients) were excluded from comparison. *p*-value, two-tailed Wilcoxon rank-sums test. **F.** Representative spatial plots of individual patients spanning a range of stromal and immune abundances. Each dot is a single cell, colored by cell type. **G.** Z-scored average expression of each protein marker in a niche cell (located up to 200 μm from anchor centroid), as a function of distance to its PanCK^high^/KRT17+ or PanCK^high^/KRT17-anchor cell. Expression is averaged across all cells for each anchor cell type, followed by standardization over all distances and anchor types.

Notably, whereas both *Hmga1* and *Hmga2* mark progenitor-like cells in mice (**Figure S1E**), only *HMGA1* was upregulated in human cells with progenitor-like signatures (**Figure S3A**). While *KRAS* mutational status could not be inferred, the coordinated activation of KRAS signaling with tumor-suppressive and senescence-related programs indicates that this rare epithelial population mirrors the core transcriptional logic of the mouse progenitor-like state.

Next, we sought to identify a robust marker for progenitor-like states that would enable spatial characterization in human samples. A pancreatic epithelial cell cluster from cancer-free donors expressing progenitor-like signatures consistently upregulated the basal marker KRT17, and recent work identified KRT17 positivity in intra-papillary pancreatic neoplasms, underscoring the capacity of pancreatic epithelial cells to adopt basal-like states in pathological conditions^58^ (**Figure S3A,C**). We thus used KRT17 as a robust epithelial anchor for identifying progenitor-like lesions in human pancreas.

Because progenitor-like cells accumulate upon tissue injury, we reasoned that local communities analogous to the progenitor niche may emerge during pancreatitis. Using highly multiplexed immunofluorescence, we analyzed 67 tissue cores from patients with mild, acute and chronic pancreatitis, including 27 non-neoplastic and 40 PDAC tumor-adjacent samples (**Data S3**), staining for epithelial identity, oncogenic and tumor-suppressive signaling, fibrosis and immune suppression markers (Methods; **Data S3**). Strikingly, we identified KRT17+/pan-cytokeratin^high^ cells (termed KRT17+ cells) in both pancreatitis and tumor-adjacent non-neoplastic samples from multiple patients, which correlated in abundance with immune cell infiltration and stromal expansion (**Figures 4C–F** and **S3D**), and tended to localize to the edge of parenchymal lobules (**Figure 4F**). Rare basal-like cells, therefore, emerge reproducibly in non-transformed human pancreas in the context of tissue injury (**Figure S3E,F**).

To assess microenvironmental changes as a function of distance from basal cells, we quantified marker expression in concentric radial bins extending from KRT17+ or KRT17-cell centroids, controlling for patient representation and tissue localization (Methods). This analysis revealed a layered architecture around KRT17+ cells—their immediate vicinity was enriched for senescence-related markers including p53, p16^INK4A^, and CRYAB; murine progenitor markers such as HMGA1, TNC and ARG1; and mitogenic signaling factors pERK and KI67; and it was depleted of CD8a and GZMB, suggesting T-cell exclusion (**Figures 4G** and **S3G**). Collectively, these data define a spatially organized niche in human pancreas that closely resembles the mouse progenitor niche in its architecture and microenvironmental composition.

### Rewiring of communication creates a progenitor niche tissue circuit

The tight spatial and transcriptional coupling of distinct cell states within the progenitor niche suggested a coordinated assembly process driven by intercellular signaling. Strikingly, as niches transition along G–P niche pseudotime, genes encoding ligands, receptors, and downstream signaling emerge among the most strongly induced programs across cellular compartments (**Figure 3F**).

To infer signaling interactions, we leveraged spatial information by enforcing that cells expressing cognate ligands and receptors must colocalize in the same niche, a critical biological constraint for communication potential (Methods). Inspection of cognate ligand–receptor pairs expressed by distinct compartments within progenitor niches revealed spatially coordinated signaling axes spanning juxtacrine signaling (e.g., epithelial *Jag1* – fibroblast *Notch3*); ECM production coupled to receptor upregulation (e.g., fibroblast *Postn* and *Tnc* – epithelial *Itgb3* and *Sdc1*); and paracrine interactions (e.g., epithelial *Pdgfb* – fibroblast *Pdgfrb*; myeloid *Nrg1* – epithelial *Itgb3*), potentially mediating MAPK activation in receiving cells^59^ (**Figure S4A**). In contrast, ligand–receptor pairs expressed in spatially segregated compartments (e.g., epithelial *Lif* – fibroblast or myeloid *Lifr*) lacked communication potential and may reflect emergent spatial patterning by antagonistic interactions^60^. Notably, cell–cell communication potential has been shown to evolve during pancreatic tumor progression^61^.

To identify signaling interactions most relevant to progenitor niche formation, we further required spatially permissible ligand–receptor pairs to change as niches transition along G–P niche pseudotime. In order to overcome Xenium signal sparsity and enable robust quantification of changes in communication potential along G–P niche pseudotime, we aggregated ligand and receptor expression across all cells from a specified cellular compartment within a niche, constructing niche-level communication matrices (Methods).

Using this framework, we observed progressive and coordinated engagement of multiple cognate ligand–receptor pairs as niches enter the progenitor state, including myeloid *Il18*–epithelial *Il18rap* and epithelial *Csf2*–myeloid *Csf2rb* (**Figure 5A,B**). These coexpression patterns indicate that bi-directional signaling becomes spatially and transcriptionally enabled between epithelial and myeloid compartments as niches transition from gastric to progenitor architecture. Importantly, ligands and receptors do not strictly need to covary; upregulation of just one component—paired with broad expression of its cognate partner—may be sufficient to confer spatially constrained communication potential. Selective upregulation of *Tgfb1* across epithelial, fibroblast, and myeloid cells of the progenitor niche (**Figure 5C**), for example, can activate TGFβ signaling in adjacent epithelial populations that ubiquitously express *Tgfbr1* and *Tgfbr2* (**Figure S4B**). This pattern is consistent with the enrichment of TGFβ transcriptional signatures in progenitor-like cells (**Figure S1M**) and, accordingly, treatment of organoid cultures derived from KP^LOH^ mice pancreata with TGFβ upregulated *Hmga2* and downregulated gastric-like cell marker *Lgals4* (Methods; **Figure 5D**).

**Figure 5.**
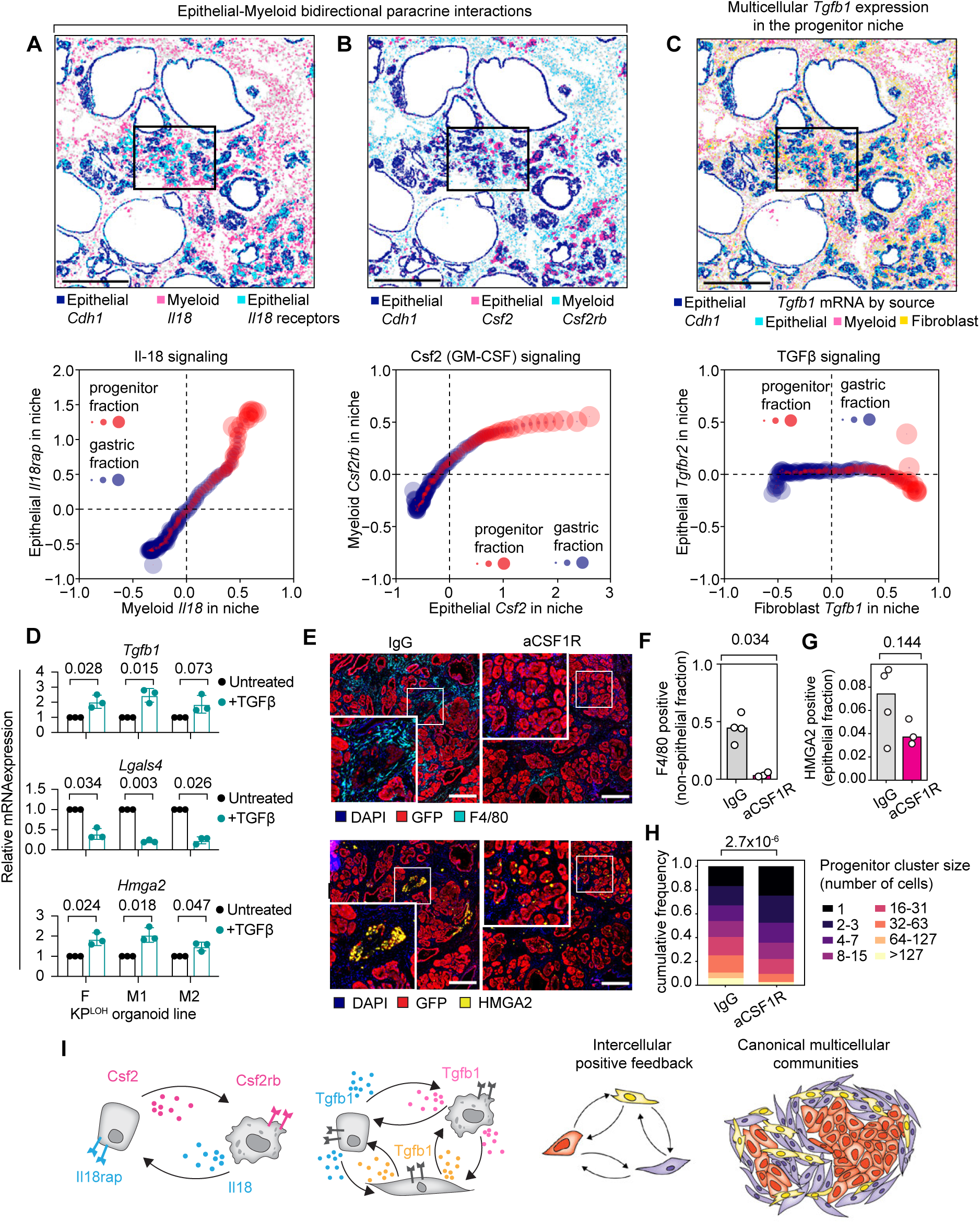
Intercellular communication modules define the progenitor niche. **A–C.** Top, Xenium data from premalignant tissue derived from KC^shCtrl^ mice, 2 days post-pancreatitis. mRNA molecules (dots) expressed by cells of a specific compartment reveal colocalization of cognate ligand–receptor pairs for IL-18 (A), GM-CSF (B), or TGFβ (C) signaling. *Cdh1* stains all epithelial cells. Bottom, corresponding plots of average niche expression of the receptor against its cognate ligand, binned along G–P niche pseudotime. **D.** Expression of *Tgfb1, Hmga2* (progenitor marker) and *Lgals4* (gastric marker) in premalignant organoids treated with recombinant TGFβ *in vitro*. Each pair of bars represents an independent organoid line. Each dot is a biological replicate. F, organoids derived from female mice; M, from male mice. p-values, paired t-test. **E.** Representative immunofluorescence images of premalignant pancreas treated with anti-CSF1R or IgG control for four weeks, harvested three weeks post-pancreatitis, and stained for F4/80 (macrophage marker) or HMGA2 (progenitor-like marker). Scale bars, 250μm. **F,G.** Fraction of non-epithelial F4/80+ (F) or epithelial HMGA2+ (G) cells in mice treated with anti-CSF1R (*n* = 3) or IgG control (*n* = 4) antibody. p-values, two-tailed Wilcoxon rank sums test. **H.** Progenitor-like cell cluster size as a function of IgG (*n* = 4) or aCSF1R (*n* = 3) treatment. p-value, Kolmogorov-Smirnov test. **I**. Schematic of intercompartmental signaling circuits enabled by the engagement of communication modules in the progenitor niche.

Given the enrichment of macrophages within progenitor niches (**Figure S2G**) and their prominent involvement in multiple epithelial–myeloid communication axes (**Figure 3F**), we next tested whether macrophages functionally contribute to progenitor niche stability *in vivo*. We treated KP^LOH^ mice with anti-CSF1R antibody to deplete macrophages or anti-IgG control, followed by caerulein treatment and continued antibody dosing for three weeks. Macrophage depletion, confirmed by IF (**Figure 5E,F**), led to a reduction in progenitor-like epithelial cells and a pronounced loss of larger progenitor lesions and cell clusters (**Figure 5G,H**), indicating that macrophages contribute to the stabilization and expansion of progenitor niches once formed.

Our findings support a model in which progenitor niches arise through a self-organizing circuit of spatially constrained, bidirectional intercellular signaling. Oncogenic KRAS activation together with inflammation triggers the emergence of progenitor-like epithelial cells, which express a distinctive repertoire of signaling ligands and receptors, enhancing their communication potential and shaping their surrounding microenvironment. In turn, these stromal populations provide feedback—through ligand production and ECM remodeling—that stabilizes the progenitor-like state and reinforces niche architecture (**Figure 5I**). The result is a self-reinforcing, cancer-like ecosystem that may facilitate malignant conversion.

### Oncogenic KRAS inhibition dismantles the progenitor niche

A key question is whether the progenitor-like state actively orchestrates niche assembly, and by extension, whether disrupting this state could destabilize the progenitor niche. Because oncogenic KRAS induces the signaling programs engaged in progenitor-like cells (**Figures S4C**), and progenitor-like cells display heightened KRAS signaling (**Figures 1L**)^7^, we hypothesized that this population—and the niche it supports—depends on sustained KRAS signaling. We subjected KP^LOH^ mice to caerulein-induced pancreatitis followed by a 48-hour pulse with the KRAS^G12D^-specific inhibitor MRTX1133^62^ (Methods; **Figure 6A**), enabling of the immediate consequences of KRAS blockade without directly perturbing the microenvironment. KRAS^G12D^ inhibition, confirmed by reduced epithelial phospho-ERK staining (**Figure S5A**), triggered rapid depletion of HMGA2+ progenitor-like cells, but did not completely ablate the epithelium (**Figure 6B**). scRNA-seq data revealed a dramatic, 24-fold depletion of progenitor-like cells after MRTX1133 treatment—significantly more than other subpopulations, such as cycling and gastric pit-like cells, which were reduced 10-fold and 8-fold, respectively (**Figure S5B,C**). Accordingly, within 4 hours of MRTX1133, apoptotic marker CC3 was induced in 44% of progenitor-like cells, but less than 6% of gastric and other cell types, across premalignant lesions (**Figure S5D,E**). Progenitor-like cells thus depend on persistent oncogenic KRAS signaling for survival.

**Figure 6.**
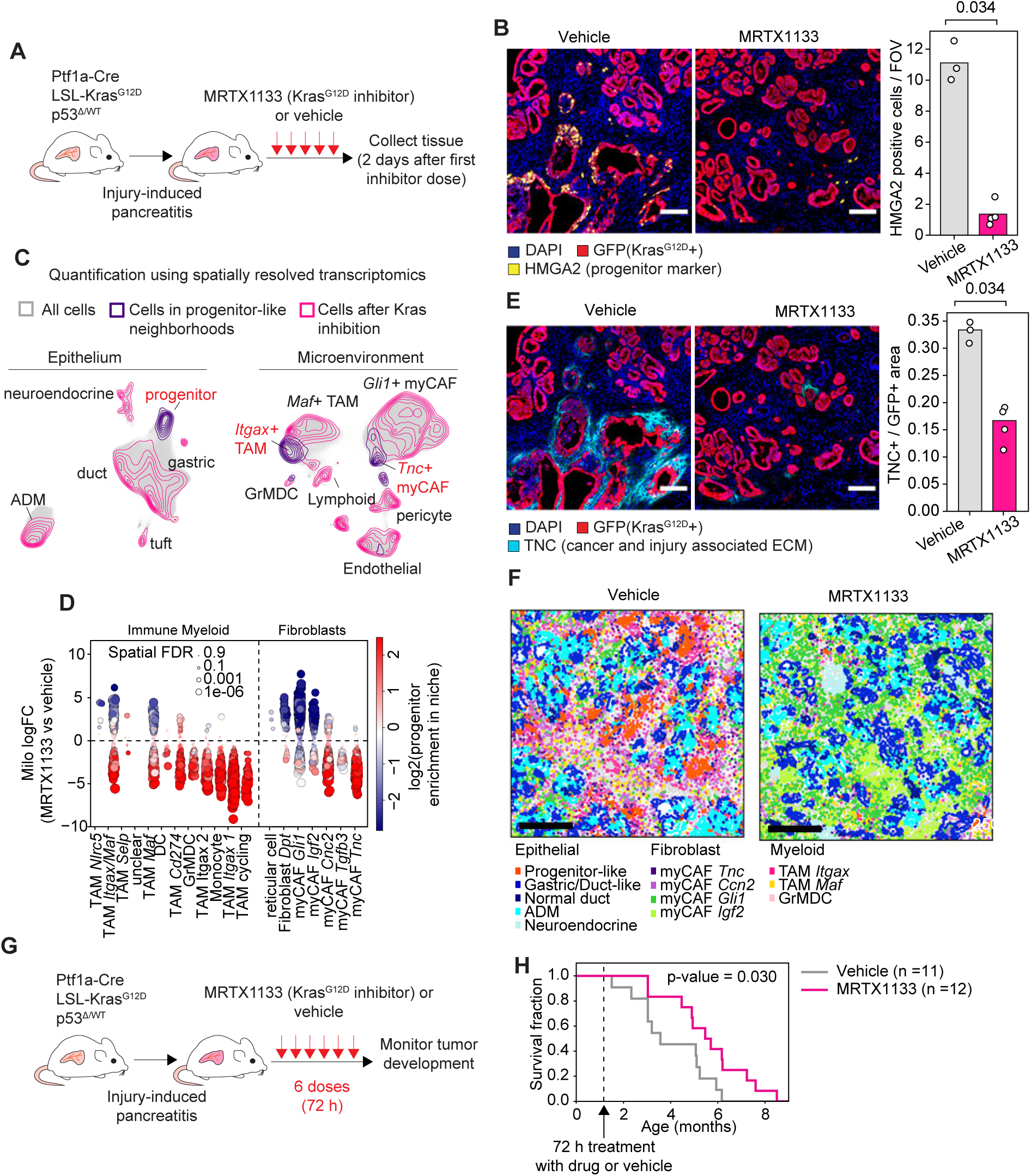
Consequences of acute oncogenic KRAS inhibition in the premalignant pancreas. **A.** Timeline of acute oncogenic KRAS inhibition in the premalignant pancreas. **B.** Representative images and quantification of HMGA2 staining in KP^LOH^ mice treated with vehicle (*n* = 3) or MRTX1133 (*n* = 4). Tissue was collected 2 days after the first dose. Scale bars, 50 μm. **C.** Xenium single-cell gene expression data from mice treated with vehicle (*n* = 2) or MRTX1133 (*n* = 4). Purple contours, density of transcriptional states in the vicinity of progenitor-like epithelial cells; pink contours, density after MRTX1133 treatment. **D.** Differential abundance of fibroblast or myeloid cell transcriptional neighborhoods in MRTX1133-treated compared to vehicle-treated samples. Each dot represents a transcriptional neighborhood defined by MiloR^63^ (Methods), colored by the enrichment of progenitor-like cells in the spatial vicinity of cells in that transcriptional neighborhood. Significantly enriched or depleted transcriptional neighborhoods are outlined in black. **E.** Representative images and quantification of TNC staining in KP^LOH^ mice treated with vehicle (*n* = 3) or MRTX1133 (*n* = 4). Tissue was collected 2 days after the first dose. **F.** Representative Xenium images from vehicle or MRTX1133 treated mice. Each dot is a cell centroid, colored by cell state. Scale bars, 250 μm. **G.** Procedure for testing the long-term consequences of acute oncogenic KRAS inhibition in the premalignant stage. **H.** Survival of mice as a function of treatment with oncogenic KRAS inhibitor or vehicle control. p-value, logrank test. **B,E.** Scale bars, 50 μm. Bars correspond to the average of individual mouse values. p-value, two-tailed Wilcoxon rank sum test.

To determine the consequences of progenitor-like cell depletion on local microenvironments, we applied Xenium profiling to premalignant pancreas tissue 48 hours after MRTX1133 (*n* = 4) or vehicle (*n* = 2) treatment. Analysis of 2,686,667 cells revealed that loss of progenitor-like cells was closely linked to widespread depletion of *Tnc+* myofibroblasts and *Itgax+* macrophages/monocytes, hallmark components of the progenitor niche (**Figures 6C–F** and **S5F**). Using our dual spatial–phenotypic representation (**Figure 2D,E**), we adapted Milo^63^ to spatial transcriptomics data, enabling us to interpret the differential abundance of transcriptional states with respect to the niches they occupy (Methods; **Figure S5G**). This analysis confirmed that the dominant effect of MRTX1133 treatment was the depletion of progenitor niche-associated populations (**Figures 6D–F** and **S5H**), along with the enrichment of populations excluded from progenitor niches, such as *Gli1+* myofibroblasts (**Figures 6D,F** and **S5H**).

Although other premalignant epithelial cell states were also affected (**Fig. S5B,C**), MRTX1133 treatment predominantly dismantled the progenitor niche itself (**Fig. 6D–F**), suggesting that progenitor-like cells actively shape their microenvironment and that targeting this state is sufficient to collapse the niche.

To assess whether early, acute depletion of the progenitor niche influences malignant progression, we treated 6-week-old KP^LOH^ mice with caerulein to induce pancreatitis, followed by a 72-hour pulse of MRTX1133, and monitored cohorts for PDAC development (**Figure 6G**). Remarkably, even this brief intervention at the earliest stages of neoplasia was sufficient to significantly delay PDAC onset (**Figure 6H**). Tumors that eventually arose in treated mice tended to have a more classical identity, marked by LGALS4 expression (**Figure S5I,J**). These data imply that depletion of the progenitor-like state and its niche produces a durable protective effect on tumor initiation.

### p53 enforces resolution of progenitor-like epithelial states

Although oncogenic KRAS inhibition rapidly dismantles the progenitor niche, this injury-induced tissue state resolves naturally over time (**Figure 2B**)^7^. What enforces the resolution of regenerative plasticity at the critical boundary between benign repair and malignant persistence? Because progenitor-like cells engage the p53 program, we hypothesized that p53 functions as an endogenous regulator of progenitor clearance during tissue injury.

To assess the consequences of *p53* loss during injury-induced tumorigenesis, we used a mouse model permitting temporally controlled *p53* suppression in the premalignant pancreatic epithelium via a doxycycline-inducible short hairpin RNA targeting *p53* (KC^shp53^), or a negative control hairpin (KC^shCtrl^)^40^ (**Figure 7A**). We transiently induced either shp53 or shCtrl for one week, triggered pancreatitis, and profiled tissue using immunofluorescence and scRNA-sequencing three weeks later.

**Figure 7.**
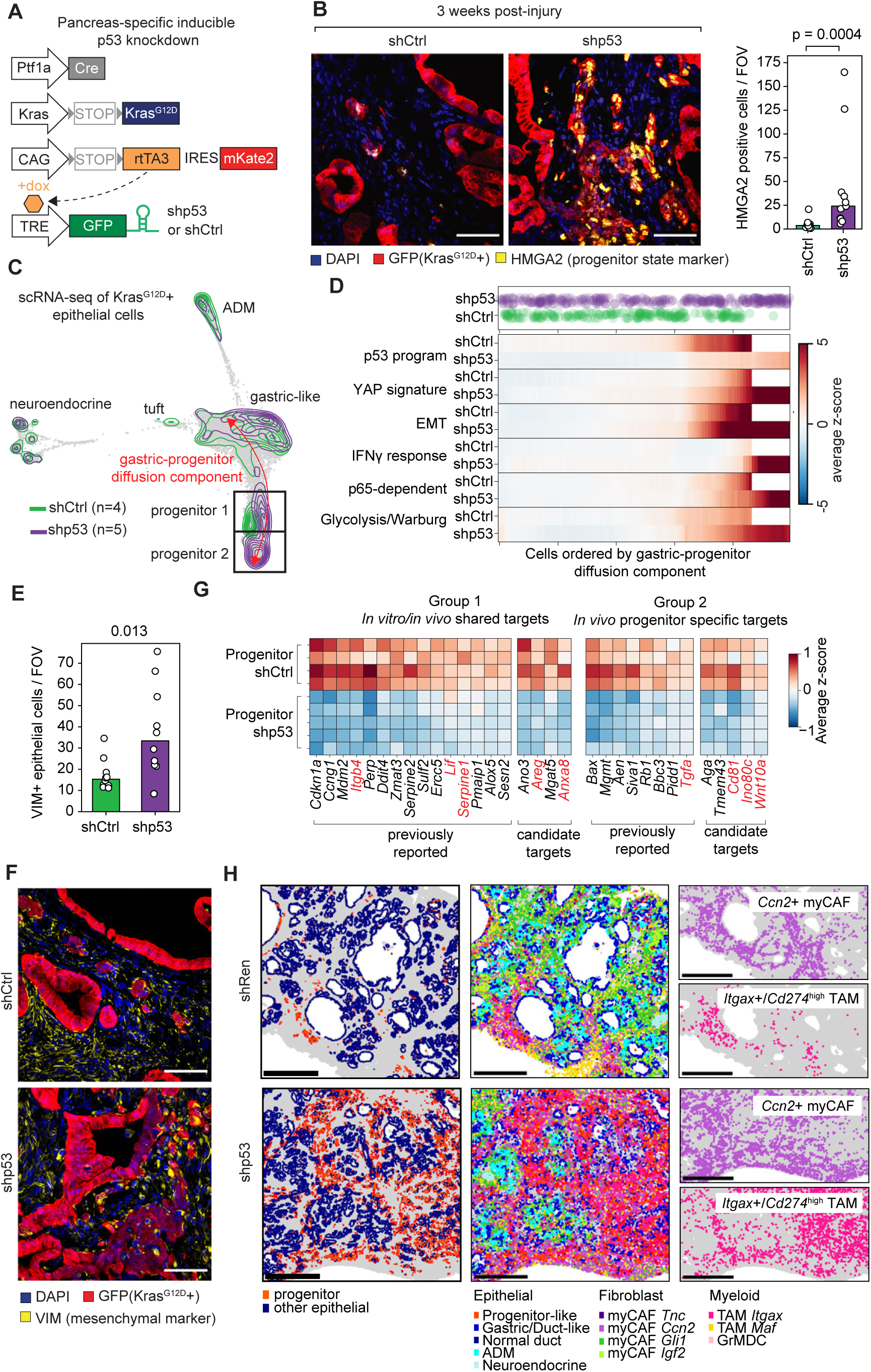
Consequences of *p53* knockdown in the premalignant pancreas. **A.** Mouse model enabling doxycycline-inducible knockdown of *p53* in the premalignant pancreatic epithelium. **B.** Representative images and quantification of HMGA2 staining 3 weeks post-pancreatitis in shp53 (*n* = 11) or shCtrl (*n* = 10) mice. **C.** FDL of scRNA-seq data from shRen (*n* = 4) or shp53 (*n* = 5) *Kras*^G12D^+ pancreatic epithelial cells. Colored contours indicate cell densities. **D.** Randomly sampled shp53 or shRen cells (top) and average score of expression signatures in shRen or shp53 cells (bottom) binned along the gastric–progenitor DC (bins <10 cells are not plotted). **E,F.** Quantification (E) and representative images (F) of VIM staining 3 weeks post-pancreatitis in shp53 (*n* = 10) or shRen (*n* = 10) mice. **G.** Representative genes downregulated upon *p53* knockdown in progenitor-like cells, highlighting both known and candidate targets. Genes in red are non-canonical p53 targets that have evidence for p63-dependent regulation in public ChIP-seq data. **H.** Representative Xenium images from KC^shRen^ or KC^shp53^ mice 3 weeks post-pancreatitis. Each dot is a cell centroid, colored by cell state for epithelial (left), epithelial and fibroblast (center) or *Ccn2*+ myofibroblast or *Itgax+/Cd274*^high^ macrophage/monocyte (right) cells. Scale bars, 500 μm. **B,F**. Scale bars, 50 μm. Bars in bar graphs correspond to the average of individual mice. p-values, two-tailed Wilcoxon rank sum test.

Whereas the injury-induced accumulation in HMGA2+ progenitor-like populations largely dissipated by this time point in control tissue, these cells persisted following *p53* loss (**Figures 7B** and **S6A**). Diffusion component analysis of scRNA-seq data (**Note S1**) revealed that *p53* loss drives epithelial cells further along the G–P continuum, giving rise to a distinct state (termed ‘progenitor 2’) with heightened mesenchymal features such as *Vim* mRNA and protein levels (**Figure 7C–F**), as well as inflammatory and oncogenic transcriptional programs, including interferon signaling, YAP signaling, and glycolysis (**Figures 7D** and **S6B,C**). These results indicate that *p53* loss facilitates progression of a RAS-driven epithelial plasticity program and imply that the ability of p53 to restrain epithelial–mesenchymal plasticity is central to its tumor suppressive role.

Next, we aimed to characterize the transcriptional program controlled by p53 during progenitor clearance. Differential gene expression analysis considering all progenitor-like cells revealed downregulation of canonical p53 targets alongside epithelial identity genes (e.g., *Cdh1*, *Epcam*) (**Figure S6D**); however, when we restricted our analysis to cells at comparable positions along the G–P axis, only repression of canonical targets remained (**Figure S6E**). Thus, loss of epithelial identity largely reflects indirect consequences of progression along the G–P axis, rather than direct p53 suppression of mesenchymal programs (**Figure S6F**). They also point to the targets and effector programs actively engaged by p53 at the time and cell state where it is actively suppressing cancer.

To refine these putative p53 targets, we isolated *p53*-proficient *Kras*^G12D^-expressing epithelial cells one day after injury and generated single-cell RNA and ATAC multiome profiles (*n* = 9,081 cells), enabling the identification of transcripts with accessible p53 response elements in progenitor-like cells (Methods; **Data S4**). We then integrated four complementary datasets: (1) transcriptional differences between gastric-like and progenitor-like states; (2) transcriptional responses to *p53* restoration in PDAC cells in vitro^64^; (3) public p53 binding annotations^65^; and (4) differential expression following *p53* knockdown *in vivo*. This integrated analysis resolved eight gene classes (**Figure S6G** and **Data S4**), including 138 genes displaying both p53-motif accessibility and p53-dependent expression in progenitor-like cells, out of which 84 were also upregulated upon p53 restoration *in vitro*, and 54 were specific to progenitor-like premalignant cells *in vivo*, highlighting context-specificity in p53-dependent gene regulation.

Pathway analysis of the inferred direct p53 target gene classes revealed a transcriptional program within the progenitor state, encompassing well-established p53 effector functions (**Figures 7G** and **S6H**; **Data S4**), including cell-cycle arrest (*Cdkn1a, Ccng1*), apoptosis (*Bbc3, Bax, Perp*), senescence (*Serpine1, Serpine2*), DNA repair (*Mgmt, Ercc5*), and metabolic stress adaptation (*Sesn2, Ddit4*). This p53-activated program also included genes implicated in less anticipated functions, such as epithelial adhesion, migration, and extracellular matrix interactions, including *Itgb4* and *Sulf2*^66^, consistent with engagement of epithelial homeostasis and repair-associated programs during progenitor-state resolution. Notably, inferred p53 targets were enriched for genes bound by p63 in other epithelial contexts (e.g., *Itgb4*^67,68^*, Areg*^69^) (**Figure S6I**), consistent with their shared response elements in p53-family members and underscoring prior links to epithelial repair and regenerative programs. These data support a model in which p53 coordinates elimination of progenitor-like cells through canonical stress response and repair associated transcriptional programs.

### *p53* naturally collapses the progenitor niche

To evaluate how *p53*-dependent epithelial states reshape tissue organization, we profiled tissues from KC^shp53^ and KC^shCtrl^ mice (9 and 5 samples, respectively; 4,611,972 cells total) using the Xenium platform. Spatial mapping uncovered a striking shift in the progenitor-like niche architecture, which expanded from small focal regions in control tissue into large, continuous domains across *p53*-deficient pancreata (**Figure S7A–C**). These changes were heterogeneous, reflecting incomplete penetrance and suggesting that progression at this premalignant stage remains probabilistic rather than deterministic (**Figure S7C**).

Expansion of progenitor niche domains in *p53*-deficient tissues was accompanied by coordinated remodeling of the surrounding tissue microenvironment. While *Ccn1*+ and *Ccn2*+ myofibroblastic cancer-associated fibroblasts (CAFs) were enriched within these niches (**Figure 7H**), the most prominent change was the accumulation of *Itgax+/Cd274^high^* (PD-L1^high^) macrophages, defining a distinct immunosuppressive state confirmed by both scRNA-seq and spatial analyses (**Figures 7H** and **S7D,E**). Macrophage abundance increased sharply once progenitor-like cells exceeded a threshold (**Figure S7F**), and these more advanced niche-associated macrophages exhibited elevated immunosuppressive and alternative-activation programs while downregulating MHC-II components (**Figure S7E**). Failure of p53 to resolve progenitor-like niches thus enables coordinated tissue remodeling and the emergence of an immune-privileged microenvironment.

Given the tight coupling between progenitor-like cells and PD-L1^high^ macrophages, we examined whether loss of *p53* rewires epithelial–immune communication in the progenitor niche. Applying the Calligraphy framework^7^ to systematically identify communication modules (**Data S5**) and statistically significant ligand–receptor interactions, we detected a restricted set of interactions that emerged specifically upon *p53* loss—most prominently, bidirectional communication between progenitor 2 epithelial cells and PD-L1^high^ macrophages, consistent with a feedback loop (**Figure S7G,H**). The inferred feedback spanned multiple signaling axes, including TGFβ-and CSF2-mediated interactions that were already active in the progenitor niche (**Figure 5**) and persisted in progenitor 2, WNT signaling from progenitor cells to macrophages, and SPP1 signaling from macrophages to progenitor cells (**Figure S7H**). These results indicate that *p53* loss strengthens multi-axis, bidirectional epithelial–immune communication, providing a plausible systems-level basis for stabilization of the immune-privileged progenitor niche.

Collectively, these findings demonstrate that either KRAS inhibition or *p53* engagement is sufficient to dismantle the progenitor-like population and its mutually reinforcing niche. While oncogenic KRAS activity drives epithelial–mesenchymal plasticity and microenvironmental remodeling—hallmarks of the regenerative phase of wound healing—p53 restrains these same processes, mirroring the resolution phase of tissue repair. Beyond its canonical role in maintaining genomic stability^28,70,71^, p53 acts as a critical barrier to both cell-intrinsic and microenvironmental plasticity, enforcing regenerative resolution at a key inflection point between benign and malignant states.

## Discussion

By visualizing and reconstructing the cellular and molecular events of the benign-to-malignant transition, we identify progenitor-like cells as the focal point of oncogenic and tumor-suppressive forces during early pancreatic tumorigenesis. This transient population, induced by oncogenic KRAS in the context of tissue injury, engages p53, CDKN2A, and SMAD4 tumor suppressive pathways, triggering senescence and intercellular communication programs that remodel the surrounding environment. Progenitor-like cells assemble a multicellular niche with hallmarks of invasive cancer, likely via reciprocal signaling between epithelial, fibroblast and immune compartments. Malignant progression ensues when this progenitor-like state escapes tumor-suppressive surveillance, enabling immune evasion, persistent epithelial plasticity, and stromal co-option. These findings position the progenitor-like state as a gatekeeper of malignant transformation, defining a discrete window during which targeting the signaling circuits and cell states that support the assembly of a cancer-like niche may enable cancer interception.

Our study was enabled by genetically engineered mouse models that capture sporadic tumor initiation by marking spontaneous *p53* loss^28,72,73^ and high-resolution spatial transcriptomics technologies^74^ that can be used to reconstruct the ecosystems surrounding progenitor-like cells. By anchoring spatial analysis on epithelial states that exist along a defined continuum, we could map niche states and gene expression trajectories, revealing coordinated shifts in epithelial plasticity, tissue architecture and remodeling events that progressively assemble the progenitor niche—a cancer-like environment characterized by immunosuppression and activated wound-healing programs—along the path to malignancy. These approaches establish a generalizable framework for investigating how cell-state transitions orchestrate tissue-scale organization in regeneration, fibrosis, and early tumor evolution.

We used our framework to pinpoint when, where, and how key oncogenic and tumor-suppressive forces act during malignant progression. Surprisingly, the major tumor suppressor pathways implicated in PDAC—p53, CDKN2A, and SMAD4—are not broadly engaged across the premalignant epithelium, but converge on a discrete progenitor-like population with high KRAS signaling—one that is transcriptionally poised for transformation. Single-cell analyses of other premalignant tissues^29,36,75–77^ have identified senescent-like or transitory cell states, suggesting that restricting progenitor-like programs represents a conserved tumor suppressive mechanism. The selective engagement of tumor suppressors in the highly plastic progenitor-like state echoes the long-standing finding that both p53 and p16^INK4A^ (a *CDKN2A* isoform) suppress induced pluripotency^78–82^, the ultimate example of cellular plasticity. Viewed through this lens, our data provide direct evidence that p53 functions as a guardian of plasticity, facilitating resolution of regenerative states that, if unchecked, promote maladaptive remodeling and tumorigenesis. Therefore, the decision between benign persistence and malignant progression occurs within the narrow window defined by the emergence, activation, and resolution of the progenitor-like state.

Aligning *p53*-proficient and *p53*-deficient epithelial cells at matched positions along the gastric–progenitor diffusion axis allowed us to distinguish direct *p53*-dependent transcriptional outputs from indirect effects due to epithelial plasticity. This analysis revealed that mesenchymal identity genes that might appear to be directly repressed by p53 are instead indirectly suppressed by p53 via its primary role in constraining plasticity. In contrast, direct p53 targets encompassed canonical tumor suppressor programs governing cell-cycle arrest, apoptosis, senescence, metabolism, and oxidative stress responses, together with a distinct set of genes linked to epithelial homeostasis, wound repair, and tissue-level regulation^83,84^. While the relative contribution of each output to progenitor-like cell depletion remains to be determined, these findings suggest that p53 coordinates multiple, context-dependent processes that together limit oncogenic plasticity and prevent the persistence of aberrant progenitor-like states that fuel malignant progression.

We further revealed the potential for bidirectional communication between progenitor-like and microenvironmental cells via coordinated activation of cell–cell signaling circuits across compartments, including TGFβ signaling, ECM/ECM–receptor communication, and immune cytokine-mediated heterotypic crosstalk. The modular and reciprocal nature of these interactions suggest that positive feedback loops stabilize the progenitor niche, consistent with principles governing regenerative and developmental transitions^85^, and a role for intercellular crosstalk in fuelling malignant progression^86^. Consistent with this model, acute KRAS inhibition in the epithelial compartment depleted progenitor-like cells while simultaneously collapsing their microenvironment, producing rapid loss of immunosuppressive macrophages and activated myofibroblasts. Conversely, macrophage depletion reduced the size and extent of progenitor lesions, indicating cooperative interactions between epithelial and innate immune compartments. Thus, malignant progression is not driven by epithelial transformation alone, but by the emergent properties of a multicellular ecosystem.

These insights converge on a broader principle: the malignant potential of progenitor-like cells is shaped by the interplay between genetic lesions, epithelial plasticity, and microenvironmental remodeling. Prior studies indicate that oncogenic KRAS derails normal regenerative programs, driving chromatin remodeling that induces a highly plastic progenitor state with heightened cell–cell communication potential^7^—features that mimic physiologic wound healing programs but become pathologically sustained in cancer^87^. In the context of regeneration, the progenitor-like state functions as both a target of tumor suppressive engagement and a hub where persistent KRAS signaling impedes wound resolution; targeting oncogenic KRAS activity or engaging p53 transcriptional programs allows resolution to proceed. These results align with emerging evidence that p53 restrains exaggerated injury responses in other epithelial tissues^75,88,89^ and support a model in which p53 and KRAS co-modulate both cell-intrinsic programs and tissue-scale dynamics to govern cancer risk. The benign-to-malignant transition can thus be viewed as a failure to terminate a normally transient regenerative state.

Our data indicate that the progenitor-like niche is preserved across disease stages and species. In early human pancreatic preneoplasia, basal-like/KRT17+, MSN+ epithelial cells reside within fibrotic, immunosuppressive microenvironments that mirror the murine progenitor niche. Similar ecosystems are present in PDAC and other malignancies marked by KRAS pathway activation and *TP53* mutation^90^, indicating that this niche becomes progressively entrenched as tumors evolve. Furthermore, a high-plasticity cell state analogous to the progenitor-like cells was recently shown to be required for lung cancer maintenance^91^. These insights inform interception strategies: transient inhibition of oncogenic KRAS during premalignancy collapsed the niche and produced a durable delay in tumor initiation, illustrating that brief perturbation of this ecosystem can redirect potentially malignant tissue trajectories. Consistent with this, KRAS inhibitors preferentially eliminate basal/mesenchymal-like populations in advanced PDAC, and remodel the tumor microenvironment in ways that resemble niche collapse, in some instances enhancing responsiveness to immunotherapy^92–95^.

*TP53* mutations are strongly associated with aggressive, treatment-refractory cancers, and our work implies this is due, in part, because p53 loss enables unchecked expansion of highly plastic epithelial cells that build a pro-fibrotic, immune-suppressive niche. Thus, rather than attempting to biochemically restore p53 function, it may be more tractable to target the pathological progenitor-like state and multicellular ecosystem that p53 naturally eliminates.

Indeed, emerging work in advanced disease contexts illustrates that niche-associated vulnerabilities can, in principle, be therapeutically exploited^90^. Our data argue that interception efforts should not only target initiating mutations but also the specific cell states and multicellular ecosystems in which p53 operates. By defining progenitor-like states and their associated niches as actionable units of tumor evolution naturally targeted by p53, our findings highlight a therapeutic paradigm that phenocopies p53 function without requiring its restoration.

## LIMITATIONS OF STUDY AND FUTURE DIRECTIONS

Here we define how wild-type p53 constrains the benign-to-malignant transition using spontaneous knockout and inducible knockdown models. While most *TP53* mutations in human PDAC are missense, some of which display allele-specific gain-of-function activities, loss of wild-type p53 activity is a universal consequence of *TP53* mutation, and a substantial fraction of cases harbor truncating alleles^96^. Whether and how mutant p53 proteins further modulate the early progenitor states and niche dynamics described here remains an important question for future studies.

How progenitor-like epithelial states are resolved is another open question. While our data establish that p53 eliminates this state, they do not distinguish whether resolution occurs through apoptosis^97,98^, senescence followed by immune clearance^99^, or differentiation^64,75,100^. The marked plasticity of this state suggests that multiple mechanisms may coexist, rather than a single deterministic outcome. Addressing these possibilities will require fate-mapping strategies and longitudinal *in vivo* approaches capable of tracking cell-state transitions at single-cell resolution across distinct *p53* genetic contexts.

Finally, our computational and perturbation analyses reveal strong interdependence between progenitor-like cells and their microenvironment. Although progenitor-like cells are particularly sensitive to acute KRAS inhibition and their niche is enriched in macrophages targeted by anti-CSF1R blockade, these perturbations also impact epithelial populations outside the progenitor niche, underscoring the coupled nature of this system. Resolving how distinct cell–cell circuits contribute to progenitor niche maintenance and resolution will require next-generation strategies for perturbing defined epithelial states or niche-associated signaling pathways *in vivo*.

## AUTHOR CONTRIBUTIONS

Conceptualization, J.R., D.P., S.W.L.; Investigation, J.R., I.D.P., A.C.C., N.P., C.S., M.T., N.R., I.M., A.F., J.P.M.IV., Z.Z., D.A.-C.; Formal analysis, J.R., A.M.E., J.P., P.W., T.K., A.M., C.B., Y.-J.H.; Data curation, N.P., A.M.E., I.M.; Software, T.K., A.M.; Project administration, P.B.R., A.F., R.C.; Supervision, M.P.d.M., D.P., S.W.L.; Funding acquisition, P.B.R., M.P.d.M., D.P., S.W.L.; Writing – original draft, J.R., D.P., S.W.L.; Writing – review and editing, J.R., T.N., D.A.-C., D.P., S.W.L.

## RESOURCE AVAILABILITY

### Lead contact

Further information and requests for resources and reagents should be directed to and will be fulfilled by the Lead Contact, Scott W. Lowe.

### Materials availability

Materials used and generated in this study, including multiallelic mESCs to generate pancreatic tumorigenesis cohorts and premalignant organoid systems, will be made available from the Lead Contact upon request.

### Data and code availability

● Single-cell RNA-seq, multiome and spatial transcriptomics data generated as part of this study have been deposited at GEO: GSE315243 and are publicly available as of the date of publication. Human pancreatic epithelial single-cell data from ref.^2^ is available at GEO: GSE226829. Injury-induced tumorigenesis data from ref.^7^ is available at GEO: GSE207920. Raw and processed imaging data will be made available from the Lead Contact upon request.
● All original code has been deposited at the Pe’er lab github repository at (https://github.com/dpeerlab/p53-niche-dynamics) and Zenodo (doi: 10.5281/zenodo.18892592), and is publicly available at of the date of publication.
● Any additional information required to reanalyze the data reported in this paper is available from the Lead Contact upon request.

## Supporting information

Reyes_2026_SupplementalData

## ACKNOWLEDGEMENTS

We thank Jiaying Tan and the anonymous reviewers for their helpful guidance, critical feedback and suggestions throughout review of our manuscript. We thank Francisco M. Barriga, Kaloyan M. Tsanov, Brinda Alagesan, Elif Ozcelik, Domhnall McHugh, Luis Soto, Tom Dougherty, Doron Haviv, Roshan Sharma, Jo Adams, Emma Chen, and members of the Lowe and Pe’er labs for insightful discussions throughout this project. We thank Mara Sherman and Tuomas Tammela for insights on single-cell heterogeneity in the premalignant pancreas; Carol Prives and James Manfredi for insights into p53 transcriptional regulation; Anupriya Singhal and Hannah Styers for protocols and advice on oncogenic KRAS inhibitor treatments; Jeff Moffitt, Brianna Watson, Jenna Hurley and Sam Aviles (Boston Children’s Hospital) for guidance and Siting Gan and Dig Vijay Kumar Yarlagadda for contributions in setting up our multiplex smFISH imaging infrastructure. We thank Robert Weinberg and Arthur Lambert for sharing their signature of p63-regulated targets in breast cancer; the Parada lab for cryostat and histology infrastructure; and Sha Tian, Wei Luan, Exequiel Sisso and Janelle Simon for technical assistance. We thank the Flow Cytometry (Fang Fang, Mark Kweens, Barbara Oliveira, Magdalena Parys), Mouse Genetics (Yasuhide Furuta, Jenny Liu and Yijie Wang), Molecular Cytology (Ning Fan, Eric Chan, Eric Rosiek), and Integrated Genomics Operation core facilities at MSK; and the Rodent Genetic Engineering Core at New York University.

## SUPPLEMENTAL INFORMATION

**Data S1.** Supporting analyses for single-cell and spatial data, related to Figures 1, 2, 3, 5, and 7 **A.** Representative FACS plot showing typical gates used for sorting mKate2+/GFP+ (*p53*-proficient) or mKate2+/GFP- (*p53*-deficient) cells from 4.5-month-old KP^LOH^ mouse, and their frequencies. a.f.u., arbitrary fluorescence units. **B,C.** FDL of *Kras*^G12D^+ epithelial cells from KP^LOH^ mice, colored by GFP mRNA expression (B) or transcriptional signatures from major premalignant cell states in pre-tumor stage mice derived from Burdziak, Alonso-Curbelo et al.^7^ (C). *p53*-deficient cells from PDAC samples and microtumor clusters are grayed out in (C). **D.** Expression of transcriptional signatures from (C) in PhenoGraph-defined clusters (*k* = 30) representing major premalignant cell states in pre-tumor stage mice (Methods). **E.** FDL from (B), colored by diffusion component 2 (DC2), capturing the continuity between gastric-like and progenitor-like premalignant cells. **F.** Binned cell density along DC2. Dashed line, DC2 threshold value used to identify progenitor-like cells (dark bars). **G.** Discretization of the gastric–progenitor continuum using DC2 threshold identified in (F). **H.** Expression of canonical p53 targets and progenitor state markers as a function of *p53* status. Each row summarizes expression in an individual sample; number of cells per sample are in parentheses. **I**. Distribution of oncogenic and tumor suppressive signature scores from Figure 1L in *p53*-proficient cells, as a function of cell state (progenitor-like or other). **J.** Expression of marker genes in single-cell reference data. Each dot represents a metacell computed using SEACells^108^ (Methods) and plotted in linear scale. Genes are ordered by their highest expression in any metacell. Dashed lines represent expression thresholds informing which markers we selected for our Xenium panel (Methods). **K.** Representative smFISH staining of marker genes in the premalignant pancreas. For each marker, an inset highlighting positive cells (top) is also displayed at high resolution (bottom). Scale bars, 20 μm. **L–P.** Left panels, representative Xenium images from a KC^shRen^ mouse, 3 weeks post-pancreatitis, showing spatial organization of transcriptional clusters in the major cellular compartments (Methods)—fibroblast (L), immune myeloid (M), immune lymphoid (N), mural cell (O) and endothelial (P). Each dot represents a cell centroid, colored by transcriptional cell state. Center panels, UMAP visualization of transcriptional clusters depicted in left panels, with cell types in corresponding colors. Right panels, Xenium-based quantification of representative marker genes for transcriptional clusters identified in each compartment. DC, dendritic cell; GrMDC, granulocytic-myeloid derived cell; ILC, innate lymphoid cell; Treg, regulatory T cell; gdT, gamma-delta T cell. **Q**. Epithelial, immune and fibroblast markers expressed in the progenitor niche (co-visualization of individual compartments in **Figure 5A–C**). Each dot is an individual mRNA, colored by the cell type that expresses it. **R.** Average compartment-wise expression of wound-healing response genes (GO Biological Processes) with highest upregulation in progenitor relative to gastric niches. Each dot represents a biological replicate (*n* = 15 mice). Values denote the average z-scored expression of wound-healing genes in either gastric or progenitor niches. p-values, two-tailed Wilcoxon rank sum test. **S–U.** Gene–gene correlation matrices of communication genes robustly expressed in epithelial (T), myeloid (U) and fibroblast (V) cells, identified in dissociated single-cell data from KC^ctrl^ and KC^shp^^53^ mice 3 weeks post-pancreatitis.

**Data S2.** Custom 10x Xenium library design with annotations of cellular compartment or biological processes probed by each gene, related to Figures 2, 3, 5, 6 and 7.

**Data S3.** Human pancreatitis tissue array metadata and multiplex immunofluorescence panel, related to Figure 4.

**Data S4.** p53 motif and program metadata, related to Figure 7.

**Data S5.** Calligraphy cell-cell communication analysis, related to Figure 7.

**Data S6**. Source data for immunofluorescence, related to Figures 1, 2, 3, 4, 5, 6, 7.

**Data S7.** Custom smFISH and 10x probe sequences and metadata, related to Figures 1 and 2.

**Data S8.** Gene signatures used in this study, related to Figures 1, 3, 4 and 7.

**Data S9.** Differential gene expression results, related to Figures 1 and 7.

**Data S10**: Quality control metrics of single-cell, multiome and spatial embeddings, related to Figures 3, 5, 6 and 7.

**Data S11.** Communication genes correlated with gastric-progenitor niche pseudotime in each cellular compartment, related to Figure 3.

**Supplementary Figure S1.**
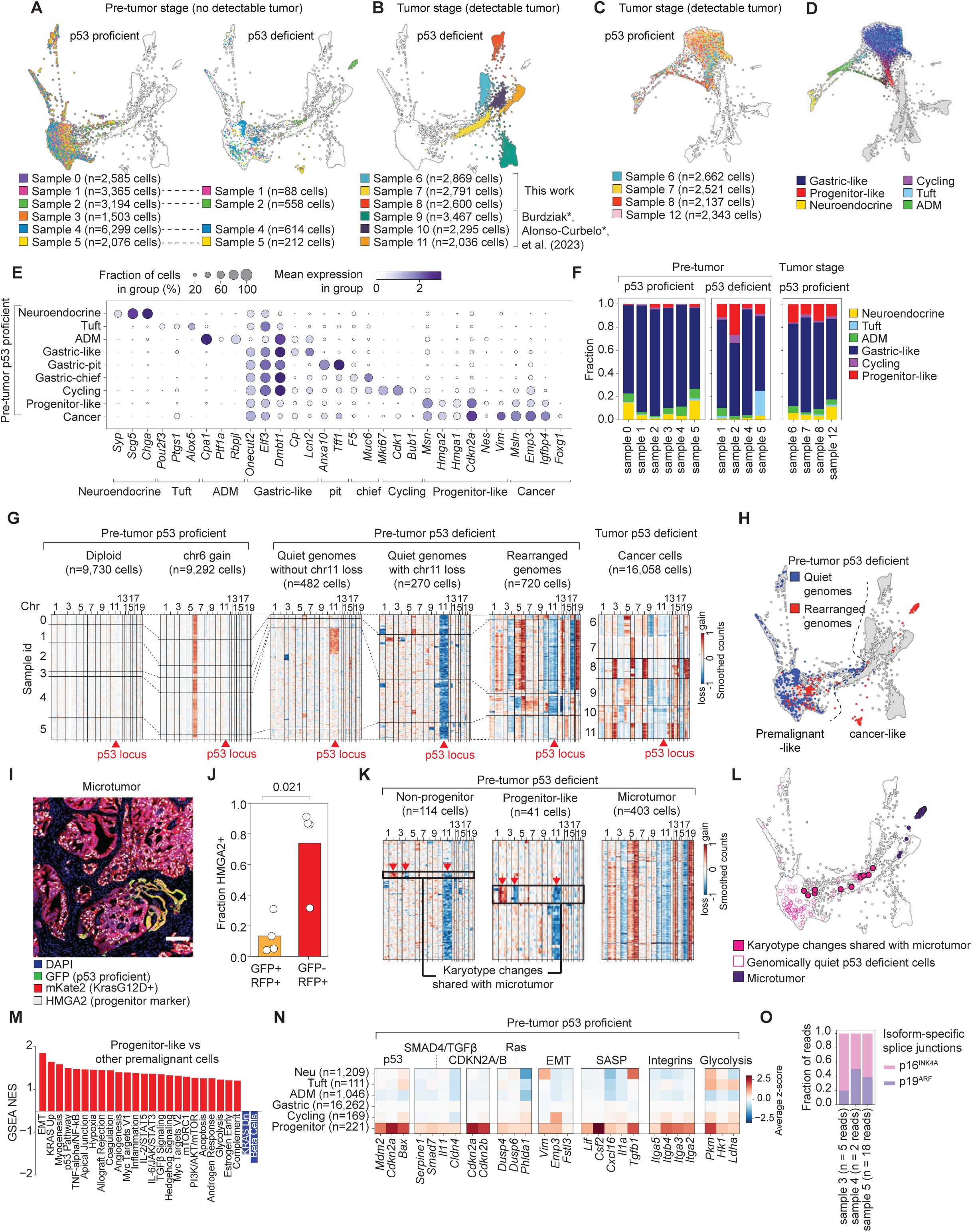
Annotation of spontaneous tumorigenesis single-cell data, related to Figure 1. **A,B.** FDL of *Kras*^G12D^+ epithelial cells from KP^LOH^ mice, colored by pre-tumor (A) or tumor (B) stage sample of origin. **C.** FDL of *p53*-proficient cells from tumor stage mice, colored by sample of origin. **D.** FDL from (C), colored by transcriptional signatures of major premalignant cell states derived from Burdziak, Alonso-Curbelo et al.^7^. *p53*-deficient cells from PDAC samples and microtumor clusters are grayed out. **E.** Expression of markers for PDAC (Cancer) or various premalignant states in distinct *Kras*^G12D^+ epithelial cell subpopulations. **F.** Fractions of premalignant states as a function of tumor stage and p53 genetic status. Each bar represents a biological replicate. **G.** InferCNV estimates of copy number change, based on log_2_-fold difference in binned gene expression from a diploid reference (Methods). Rows correspond to cells, columns to ordered genomic regions. Chr, chromosome number. Cells are partitioned by PDAC development stage, *p53* status or extent of karyotype changes, then grouped by originating sample and clustered by karyotype profile within each partition; dashed lines connect cells from the same sample. **H.** FDL of *Kras*^G12D^+ epithelial cells from KP^LOH^ mice, with pre-tumor stage (*p53*-deficient) cells colored by genomic state. **I.** Representative immunofluorescence image of microscopic PDAC detected in pre-tumor stage KP^LOH^ mice. Scale bar, 100μm. **J.** Fraction of HMGA2+ cells in *p53*-deficient microtumors (GFP-mKate2+) or *p53*-proficient regions (GFP+ mKate2+). Each dot is one mouse; p-value, two-tailed Wilcoxon rank sum test. **K.** Karyotypes of *p53*-deficient cells in a pre-tumor stage mouse, partitioned by karyotype and transcriptional profiles consistent with PDAC (Microtumor), progenitor-like or other premalignant states. Boxes highlight cells with premalignant transcriptional profiles that share a subset of karyotype changes with microtumor cells. **L.** FDL of *p53*-deficient cells from a single sample, colored by karyotype status. **M.** Gene set enrichment analysis computed using log_2_-fold changes in gene expression between *p53*-proficient progenitor-like cells and other *p53*-proficient premalignant cells, as estimated by differential gene expression (Methods). **N.** Expression of representative genes upregulated in progenitor-like cells, corresponding to tumor suppressive and oncogenic signatures in Figure 1L. **O.** Fraction of isoform-specific *Cdkn2a* reads in pre-tumor *p53*-proficient samples.

**Supplementary Figure S2.**
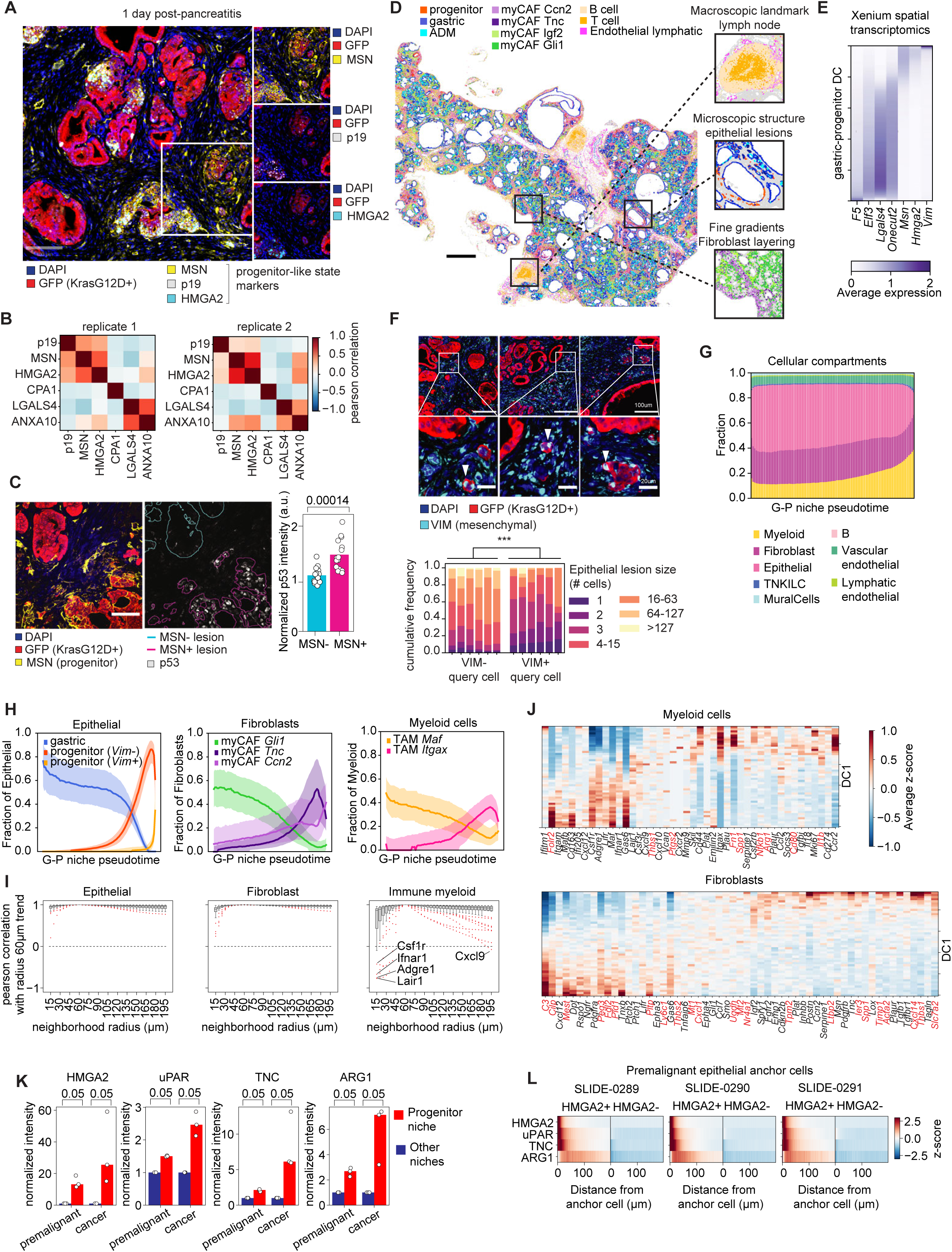
Identification of progenitor-like lesions upon pancreatic injury, related to Figures 2 and 3. **A.** Multiplexed immunofluorescence staining of progenitor-like markers in tissue from a 6-week-old pre-tumor stage KP^LOH^ mouse, 1 day post-pancreatitis, on the COMET (Lunaphore) platform. Inset highlights a progenitor lesion. Right, inset region from left, showing each progenitor marker individually with premalignant epithelium (GFP) and nuclei (DAPI). Scale bar, 100μm. **B.** Pearson correlation between the expression of premalignant epithelium markers, quantified with the COMET platform. **C.** Immunofluorescence staining (left) and quantification of p53 protein (right) in 6–7-week-old KC^shRen^ mice, 2 days post-pancreatitis. Left and right images are adjacent tissue sections; epithelial lesions were manually identified for MSN status in the MSN-labeled tissue sections (blinded for p53 staining). p-value, two-tailed Wilcoxon rank sum test. Scale bar, 50μm. **D.** Representative Xenium images from a KC^shRen^ mouse, 3 weeks post-pancreatitis (Methods). Each dot represents a cell centroid, colored by transcriptional cell state. **E.** Average expression of select gastric and progenitor markers as a function of gastric–progenitor DC bin in premalignant epithelial cells, quantified using the Xenium platform. **F.** Immunofluorescence staining and quantification of the mesenchymal marker VIM in KC^shRen^ mice harvested 3 weeks post-pancreatitis. Each bar is a biological replicate; ***, p < 0.001, Kolmogorov-Smirnov test on pooled replicates. Scale bars (top) 100μm, (bottom) 20μm. **G**. Coarse cell type distribution in niches as a function of the average gastric–progenitor DC in the niche (gastric niches at left) in Xenium data from *Kras*-mutant, *p53*-proficient pancreatic epithelium (pre-tumor KC^shRen^ or KP^LOH^), pooling tissue samples 2 days and 3 weeks post-pancreatitis (Methods). **H.** Frequency of select cell populations as a function of average gastric–progenitor DC in niche epithelial cells. Bold lines, median fraction; shaded areas, interquartile range. **I.** Robustness analysis of compartment-specific gene expression trends as a function of the gastric–progenitor niche continuum. Boxplots show Pearson correlation between gene trends of specific genes computed on neighborhoods of a specified radius (x-axis) relative to our reference 60-μm radius. **J.** Gene expression in myeloid cells and fibroblasts from dissociated single-cell data from 2 mice with *Kras*-mutant, *p53*-proficient pancreatic epithelium (KC^shRen^ or uninduced KC^shp53^), 3 weeks post-pancreatitis. Cells are in 50 bins along the major axis of variation (diffusion component 1, DC1) within each cellular compartment. Genes highlighted in red were not measured in Xenium data. **K.** Protein expression in 60-μm cellular niches surrounding HMGA2+ (red bars) or HMGA2-cells (navy bars), in premalignant stage (*n* = 3) or cancer-bearing (*n* = 3) KP^LOH^ model mice. For premalignant mice, both HMGA2+ and HMGA2-cells are also GFP+ (*p53*-proficient). For cancer-bearing mice, HMGA2+ cells are GFP- (*p53*-deficient) and HMGA2-cells are also GFP+ (*p53*-proficient). Expression is normalized to the average of HMGA2-niches within each biological replicate. p-values, two-tailed Wilcoxon rank sum test. **L.** Z-scored average expression of each protein marker in a niche cell, as a function of distance to its GFP+/HMGA2+ or GFP+/HMGA2-anchor cell. Expression is averaged across all cells for each anchor cell type, followed by standardization over all distances and anchor types.

**Supplementary Figure S3.**
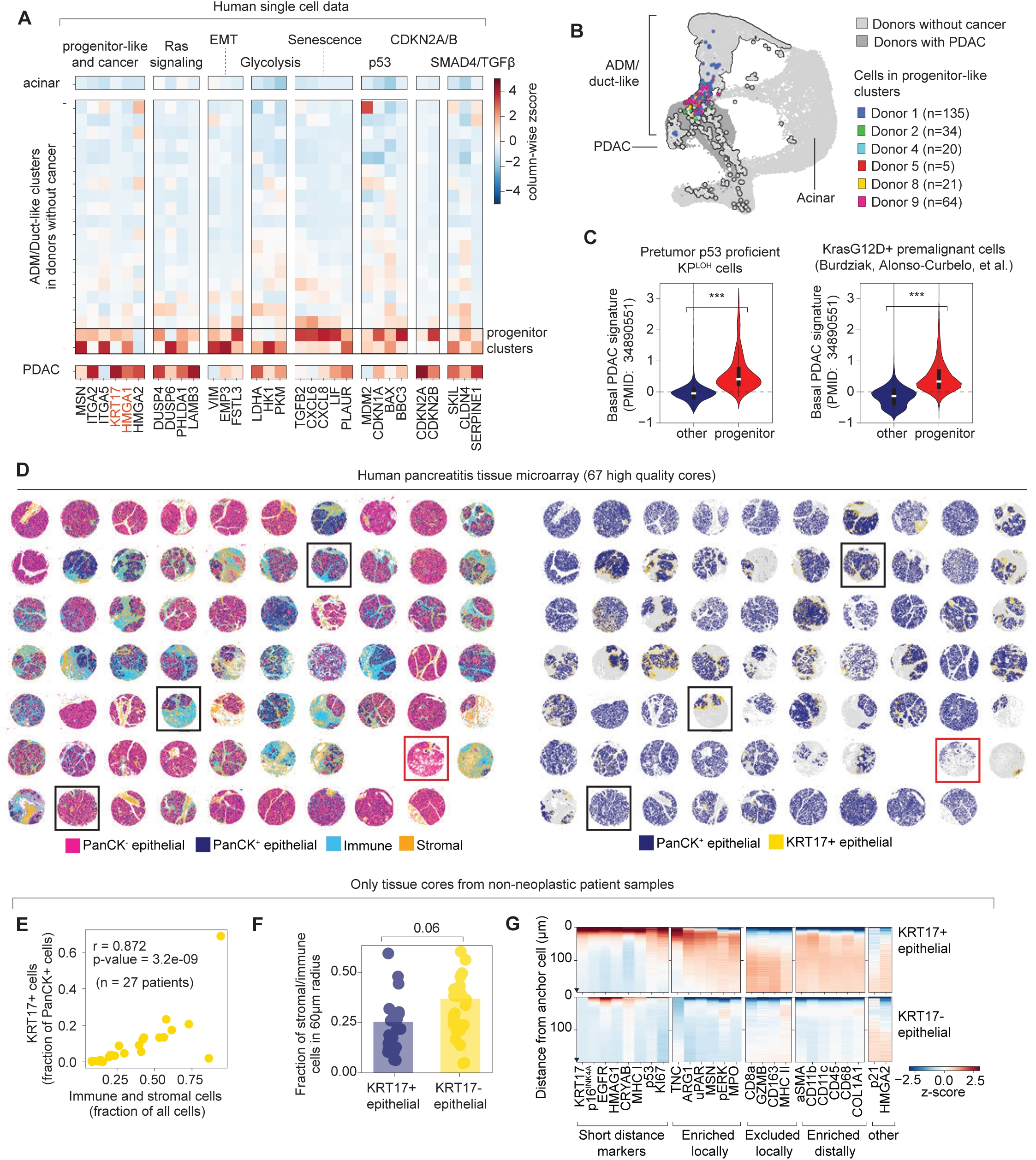
Detection of basal-like cells in non-neoplastic human pancreatic tissue, related to Figure 4. **A.** Average z-scored expression of representative genes from oncogenic and tumor suppressive signatures associated with the progenitor-like state in human-derived single-cell data from Carpenter, Elhossiny, Kadiyala, et al.^2^. Each row of ADM/duct-like cells corresponds to a PhenoGraph cluster (Methods); two PhenoGraph clusters with highest expression of progenitor-like signatures from Figure 2F (progenitor clusters) are boxed. **B.** UMAP of cells from human pancreatic samples. ADM/duct-like cells are outlined are progenitor-like cells from (A) are colored by donor. **C.** Basal PDAC signature scores (average z-scored expression of signature genes) in progenitor-like cells or other cells from pre-tumor *p53*-proficient cells derived from the KP^LOH^ model, or *Kras*^G12D^+ premalignant epithelial cells from our single-cell atlas of pancreatic tumorigenesis^7^. p-values, two-tailed Wilcoxon rank sum test. **D.** Spatial plots of the tissue array. Each dot is a cell, colored by cell type. Tissue cores boxed in black are shown in Figure 4F. The core boxed in red was removed from analysis due to poor staining. **E.** Fraction of epithelial cells that are KRT17+, as a function of the fraction of stromal and immune cells in individual patient samples that are not neoplasia associated. p-value, Spearman’s rank correlation. **F.** Niche composition of KRT17+ or KRT17-epithelial cells in patient samples that are not associated with neoplasia. Each dot is a single patient. Patients samples with fewer than 15 KRT17+ cells (*n* = 3 patients) were excluded from comparison to limit effects stemming from a limited number of cells. p value, two-tailed Wilcoxon rank-sums test.**G.** Z-scored average expression of each protein marker in a niche cell, as a function of distance to its PanCK^high^/KRT17+ or PanCK^high^/KRT17-anchor cell. Expression is averaged across all cells for each anchor cell type, followed by standardization over all distances and anchor types.

**Supplementary Figure S4.**
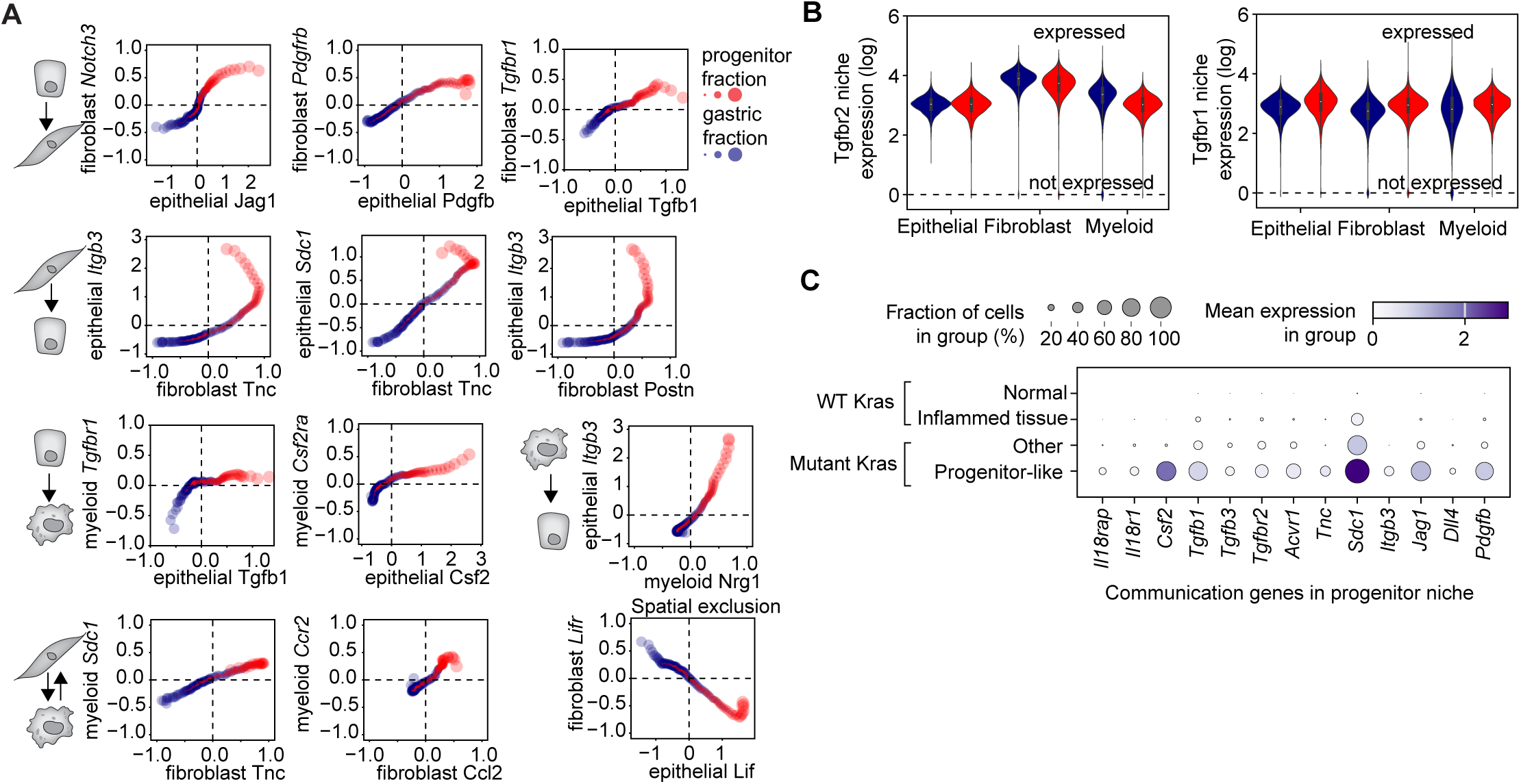
Concerted upregulation of cognate ligand–receptor pairs in the progenitor niche, related to Figure 5. **A.** Expression of representative cognate ligand–receptor pairs as a function of the average gastric–progenitor DC in the niche. Each dot represents the averaged z-scored niche expression of the specified genes in their corresponding cellular compartment (Methods). Dot color and size correspond to the relative frequency of gastric or progenitor-like cells in the niche. **B.** Expression of markers associated with the progenitor niche in epithelial, immune and fibroblasts. Each dot is an individual mRNA, colored depending on the producer cell type. **C.** Dissociated single-cell expression in pancreatic epithelial cells. Top row: acinar cells from uninjured mice^7^; second row: ADM cells 2 days after pancreatic injury in the context of WT *Kras*^7^; third row: *Kras*^G12D^+ cells outside of the progenitor-like state, harvested 3 weeks post-pancreatitis (this work); fourth row: *Kras*^G12D^+ progenitor-like state harvested 3 weeks post-pancreatitis (this work). Dot size represents the fraction of cells that express a given mRNA in the group. Dot color represents the level of expression in positive cells within the group.

**Supplementary Figure S5.**
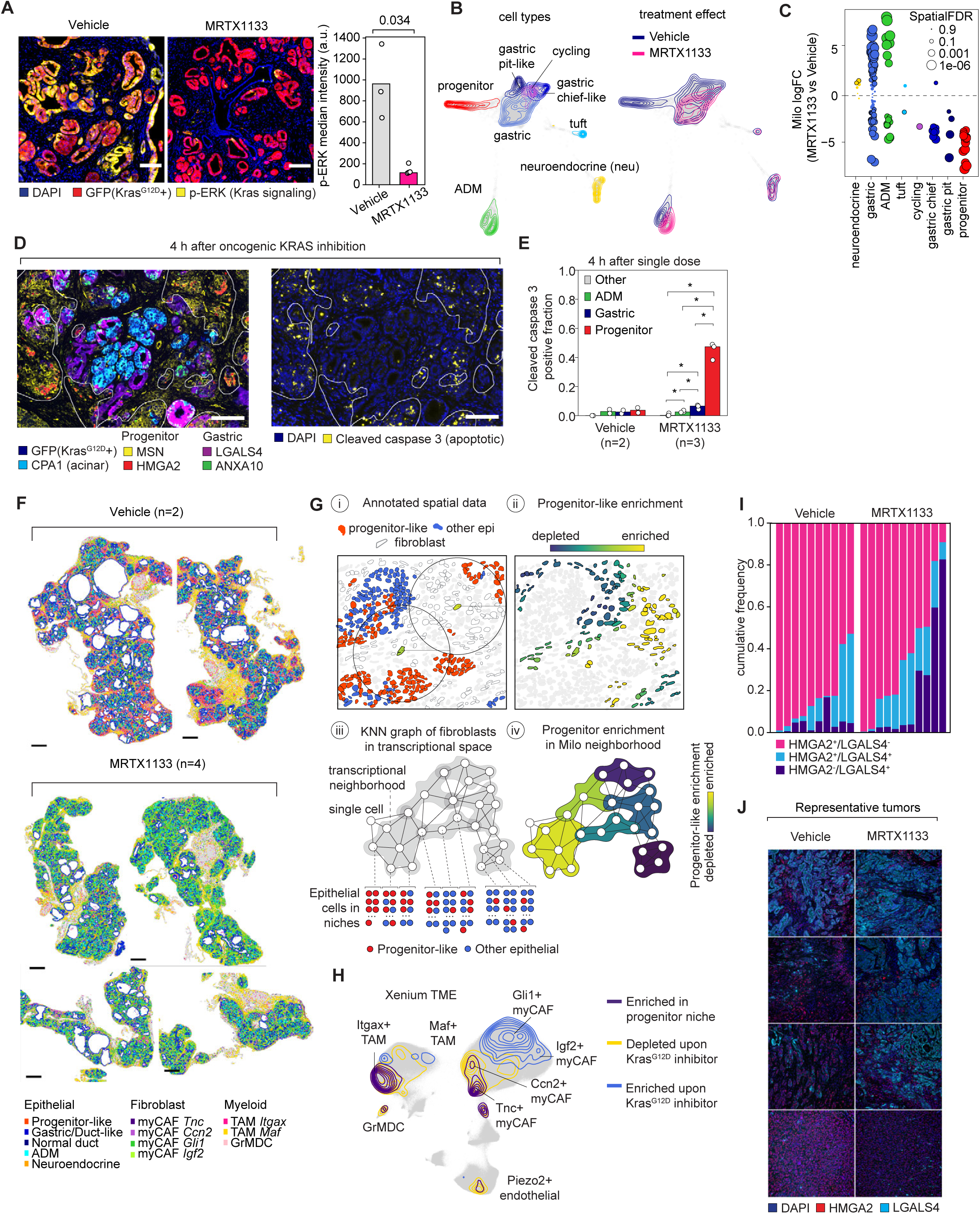
Consequences of acute oncogenic KRAS inhibition in premalignant cells, related to Figure 6. **A.** Representative images and quantification of p-ERK staining in KP^LOH^ mice 2 days after treatment with vehicle (*n* = 3) or MRTX1133 (*n* = 4). Bars represent averages of median p-ERK signal in individual mice. *p*-value, two-tailed Wilcoxon rank-sums test. **B.** FDL of scRNA-seq data from *Kras*^G12D^+/*p53*-proficient pancreatic epithelial cells derived from MRTX1133 (*n* = 3) or vehicle (*n* = 3) treated mice during injury-induced pancreatitis, overlaid with density maps of premalignant state (left) or treatment condition (right). **C.** Differential abundance testing for differences between MRTX1133 or vehicle-treated mice using MiloR^63^. Each dot is a transcriptional neighborhood of single cells, grouped and colored by the most abundant cell state in the neighborhood. Outlined dots show transcriptional neighborhoods that are differentially abundant between conditions (SpatialFDR < 0.1). **D.** Representative multiplex immunofluorescence images of premalignant state markers (left) and the apoptotic marker cleaved caspase 3 (CC3) in premalignant pancreas harvested 4 hours after a single dose of MRTX1133. Outlines denote progenitor lesions. Scale bar, 100μm. **E.** Fraction of CC3-positive cells in each premalignant state, as a function of treatment (vehicle *n* = 2, MRTX1133 *n* = 3). *, p-value = 0.05, the smallest value possible from a one-sided Wilcoxon rank sum test with two groups of *n* = 3. **F.** Representative Xenium images of whole pancreas tissue sections from vehicle (left) or MRTX1133 (right) treated pre-tumor stage KP^LOH^ mice. Each dot is a cell centroid, colored by cell states that demarcate differences between gastric and progenitor niches. **G.** Integration of niche-level analysis with differential abundance testing. (i) For each cell, we calculate the fraction of epithelial cells annotated as either progenitor-like or gastric-like in their niche (the plot highlights two fibroblast cells and their niches). The progenitor enrichment score is defined as the log ratio of the fraction of progenitor-like cells in the niche and the fraction of progenitor-like cells in the tissue. (ii) We quantify enrichment scores systematically to generate a tissue map reflecting continuous spatial variation in progenitor-like cell abundance. Fibroblasts in the image are colored by their progenitor enrichment score. (iii) We construct transcriptional neighborhoods using MiloR. For each cell in a Milo neighborhood, we extract epithelial cells in their niche (bottom). (iv) We use the fraction of progenitor-like cells relative to other epithelial cells in the niche of cells identified in (iii) to compute a progenitor enrichment score for every Milo neighborhood (Methods). **H.** Xenium single-cell gene expression data from mice treated with vehicle (*n* = 2) or MRTX1133 (*n* = 4). Purple contours, density of transcriptional states in the vicinity of progenitor-like epithelial cells; gold contours, density of states depleted after MRTX1133 treatment; royal blue contours, density of states enriched after MRTX1133 treatment. **I.** Distribution of HMGA2+ (basal PDAC marker), LGALS4+ (classical PDAC marker) or HMGA2+/LGALS4+ (mixed) cells in PDAC harvested at end point following acute treatment with MRTX1133 or vehicle at the premalignant stage. **J.** Representative tumors from data quantified in (I).

**Supplementary Figure S6.**
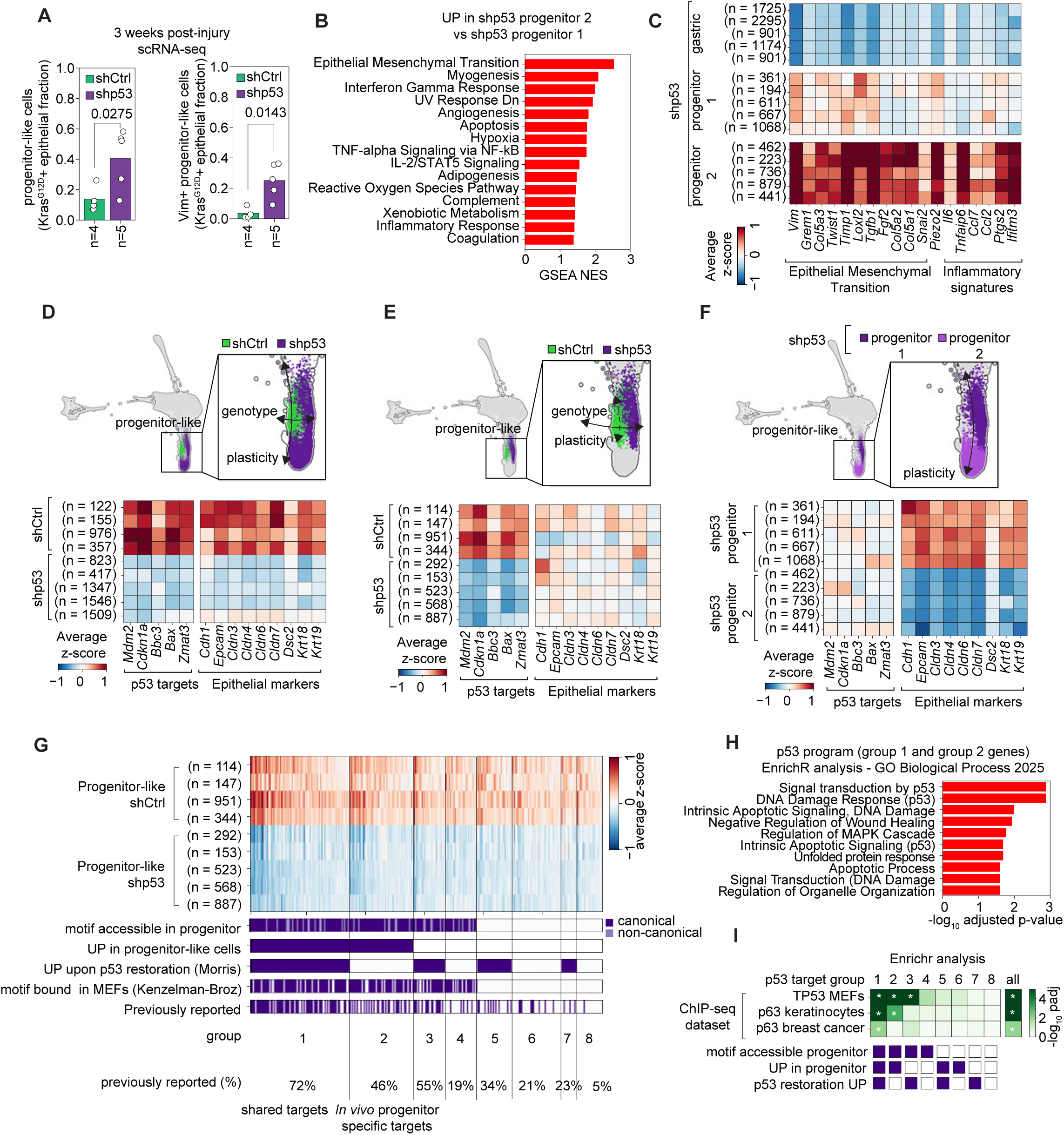
Transcriptional consequences of *p53* loss in the context of oncogenic KRAS activation and pancreatic injury, related to Figure 7. **A.** scRNA-seq-based quantification of Hmga2+ (left) or Vim+ (right) epithelial cells in KC^shp53^ or KC^shCtrl^ mice 3 weeks post-pancreatitis. Bars represent the average of individual mice. p-value, two-tailed Wilcoxon rank sum test. **B.** Gene set enrichment analysis (GSEA) for GO Biological Process 2025 gene sets of transcripts upregulated in progenitor 2 shp53 cells. Log_2_-fold changes in gene expression between PhenoGraph-clustered progenitor 1 and progenitor 2 shp53 cells, calculated using diffxpy (https://github.com/theislab/diffxpy), were used as the ranking variable for gene set enrichment analysis. All signatures shown are enriched or depleted with FDR < 0.1. NES, normalized enrichment score. **C.** Expression of representative genes of select signatures enriched in progenitor 2 over progenitor 1 cells, as identified in (B). Columns represent individual biological replicates, grouped by premalignant cell state. Parentheses denote number of cells per group. **D–F.** Top, FDL of *Kras*^G12D^+ pancreatic epithelial cells derived from KC^shCtrl^ (*n* = 4 mice) or KC^shp53^ (*n* = 5) samples, 3 weeks post-pancreatitis. Box highlights progenitor-like clusters, which are colored by *p53* knockdown status (D,E) or cell state (F), and are partitioned in three ways (Methods): aggregated across biological replicates to summarize gene expression regardless of progression along an epithelial–mesenchymal plasticity axis (D); aggregated the same way, but excluding a cluster that accumulates in KC^shp53^ and is rare in KC^shCtrl^ (E); and only aggregating KC^shp53^ cells as a function of cluster assignment (F). Bottom, expression of canonical p53 targets or epithelial identity markers in *p53*-proficient (shCtrl) or *p53*-deficient (shp53) progenitor-like cells. **G.** Top, expression of candidate p53 targets in *p53*-proficient (shCtrl) or *p53*-deficient (shp53) progenitor-like cells, standardized only within this cell subpopulation. Bottom, annotation of genes according to in-house and public datasets. **H.** Pathway analysis of candidate p53 targets (group 1 and group 2) using Enrichr and the GO Biological Process 2025 database. **I.** Significance of enrichment of p53 or p63-bound targets in public ChIP-seq data (TP53 MEFs^65^, p63 keratinocytes^106^, p63 breast cancer^67^ using Enrichr^107^. Analysis was conducted independently for each group of candidate p53 targets defined in (G), or for all predicted p53 targets. Asterisk, adjusted p-value < 0.05.

**Supplementary Figure S7.**
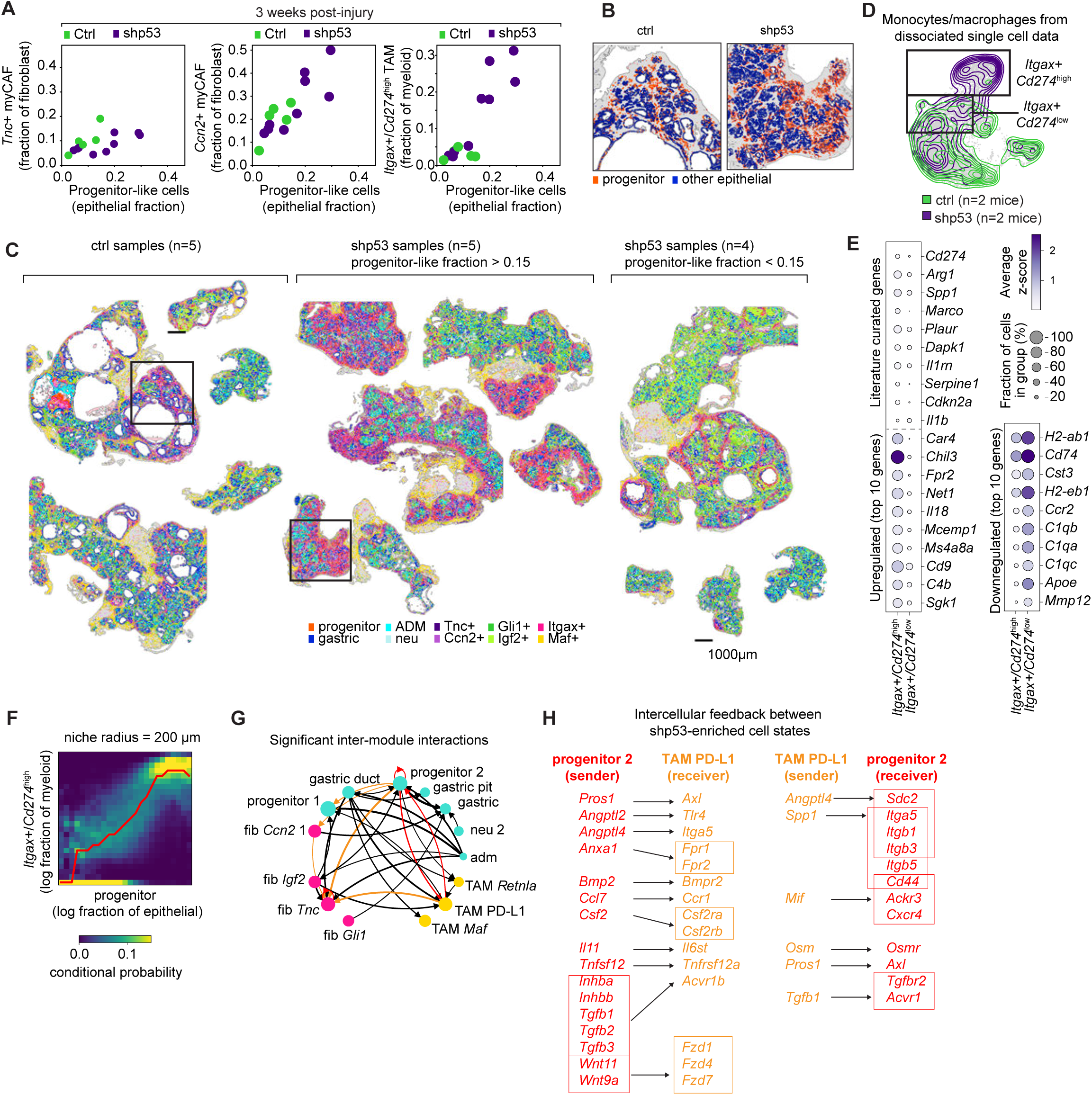
Tissue remodeling upon p53 knockdown in the context of oncogenic KRAS and pancreatic injury, related to Figure 7. **A.** Xenium-based quantification of microenvironment subpopulations associated with the progenitor niche, as a function of the fraction of progenitor-like cells in the tissue. Each dot is a single biological replicate. **B.** Representative Xenium data in control or *p53* knockdown tissue, colored by epithelial cell state. Each dot is a single cell centroid. **C.** Visualization of progenitor niches in our entire collection of KC^shRen^ and KC^shp53^ tissues collected 3 weeks post-pancreatitis. Each dot is a single cell colored by cell state. Boxes denote regions highlighted in (B). **D.** UMAP of macrophage/monocyte cells from dissociated single-cell data from the premalignant pancreata of shp53 (*n* = 2) and control (*n* = 2) mice, 3 weeks post-pancreatitis, colored by condition of origin. **E.** Subset of genes differentially expressed in *Itgax*+/*Cd274*^high^ vs *Itgax*+/*Cd274*^low^ cells. The top group is selected for known roles in immune regulation and tissue injury, and lower left and right groups include the top 10 upregulated and downregulated genes in *Itgax*+/*Cd274*^high^ cells, respectively. **F.** Distribution of the abundance of *Itgax*+/*Cd274*^high^ cells (fraction of myeloid), conditioned on the abundance of progenitor-like cells (fraction of epithelial) in the niche. Red line denotes trend. **G**. Significant inter-module interactions (p < 0.05) identified using Calligraphy^7^. Arrow weight is proportional to -log_10_(p-value). Orange arrows highlight interactions between cell states coexisting in the progenitor niche, as identified in our spatial data. Red arrows highlight feedback between progenitor 2 and PD-L1^HIGH^ macrophages/monocytes. **H.** Ligand–receptor pairs that support inferred feedback between progenitor 2 and PD-L1^high^ macrophages/monocytes. Boxes denote groups of ligands or receptors that interact with the same cognate communication gene.

**Data S1.1.**
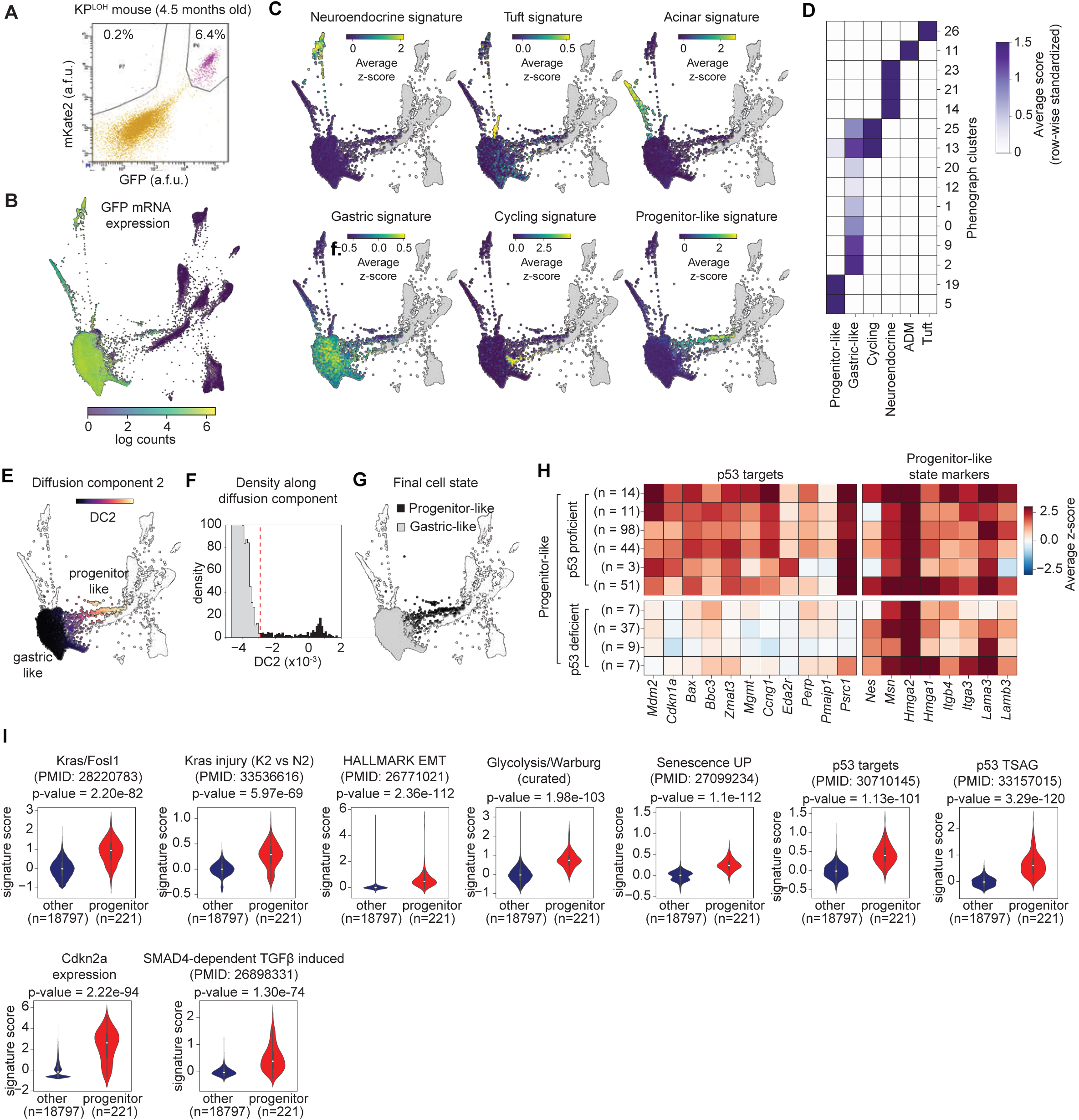
Supporting analysis of single cell data, related to Figure 1.

**Data S1.2.**
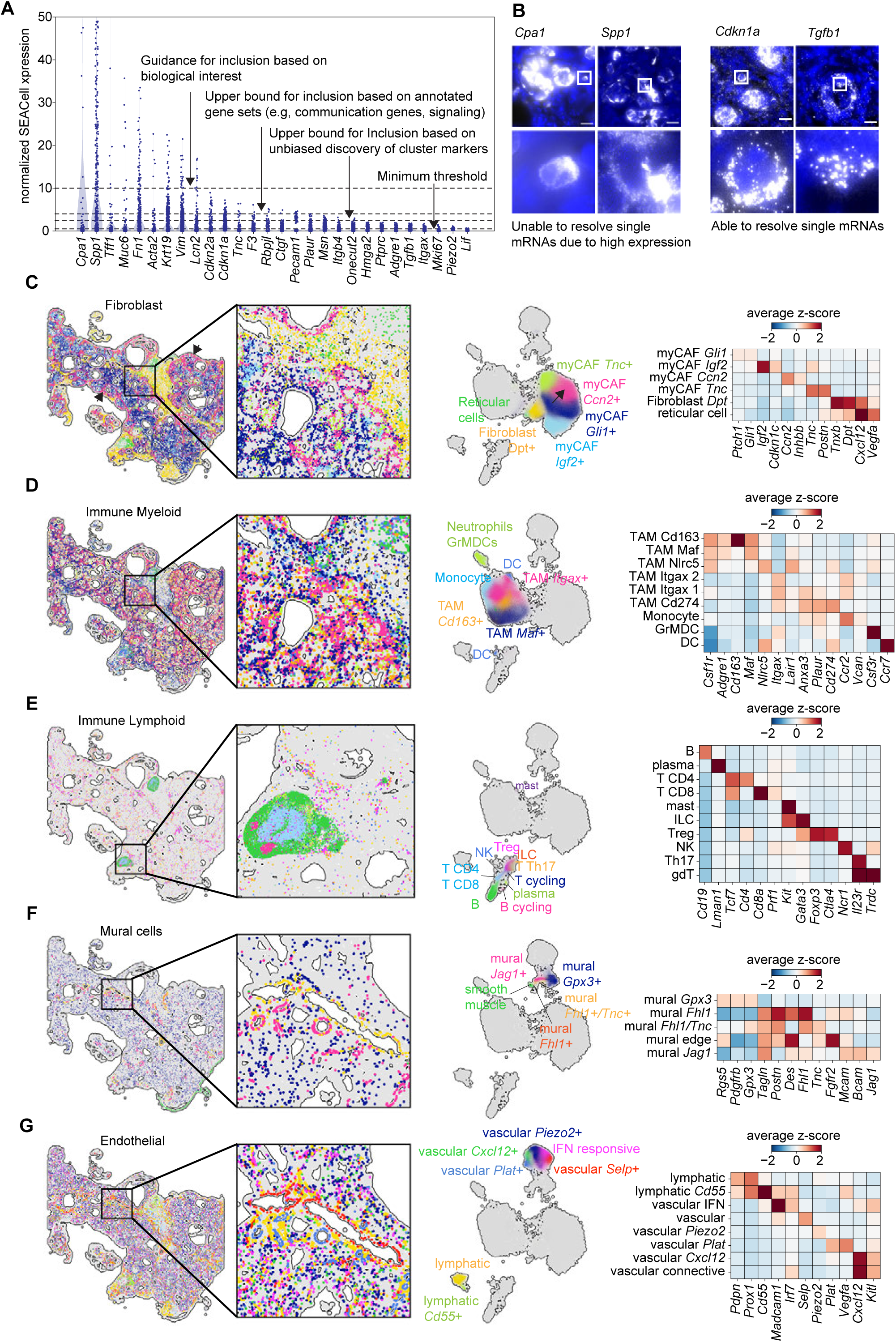
Supporting analysis of spatial data related to Figures 2 and 3.

**Data S1.3.**
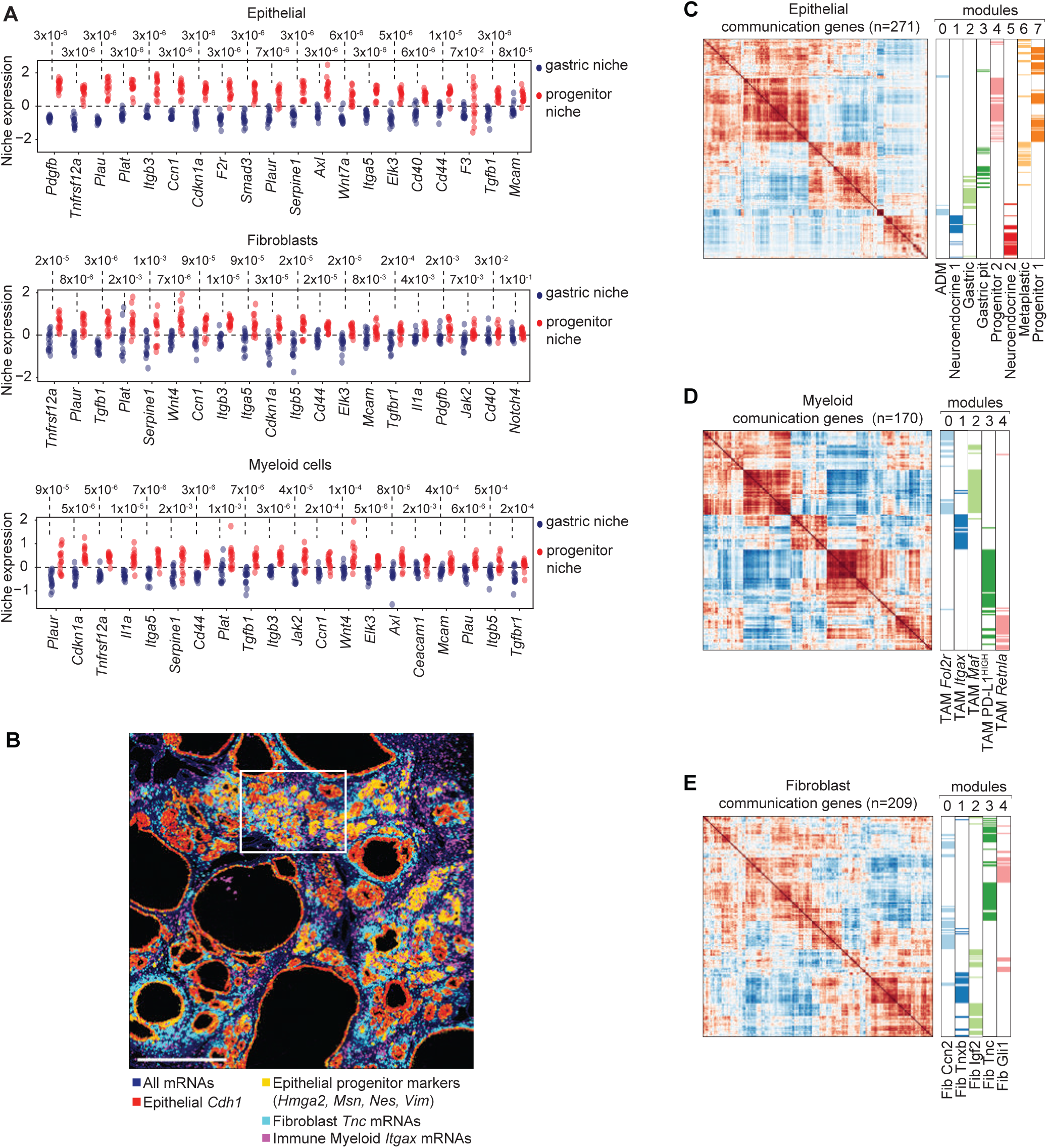
Supporting analysis of single-cell and spatial data, related to Figures 3, 5 and 7.

**Table S1.**
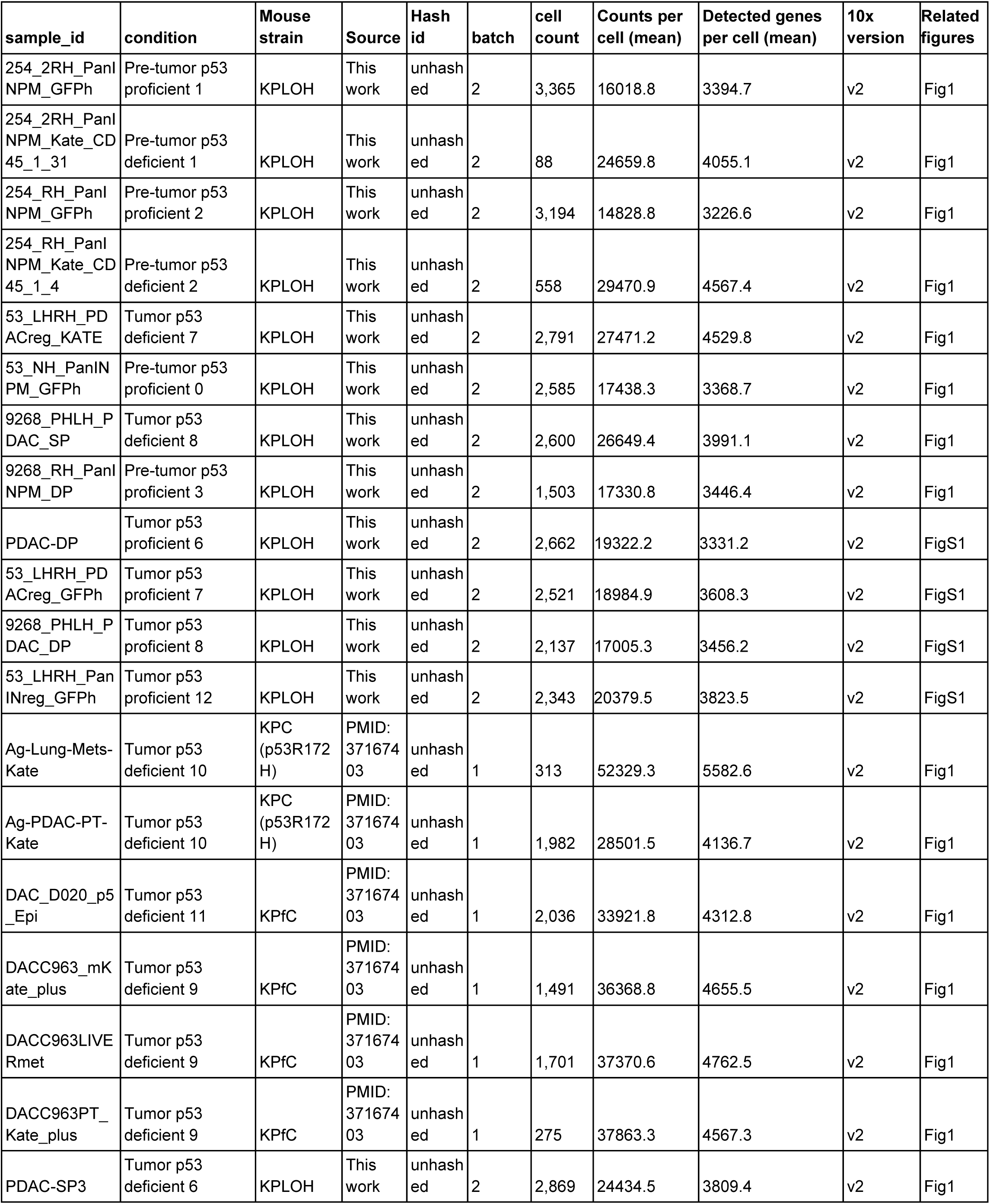
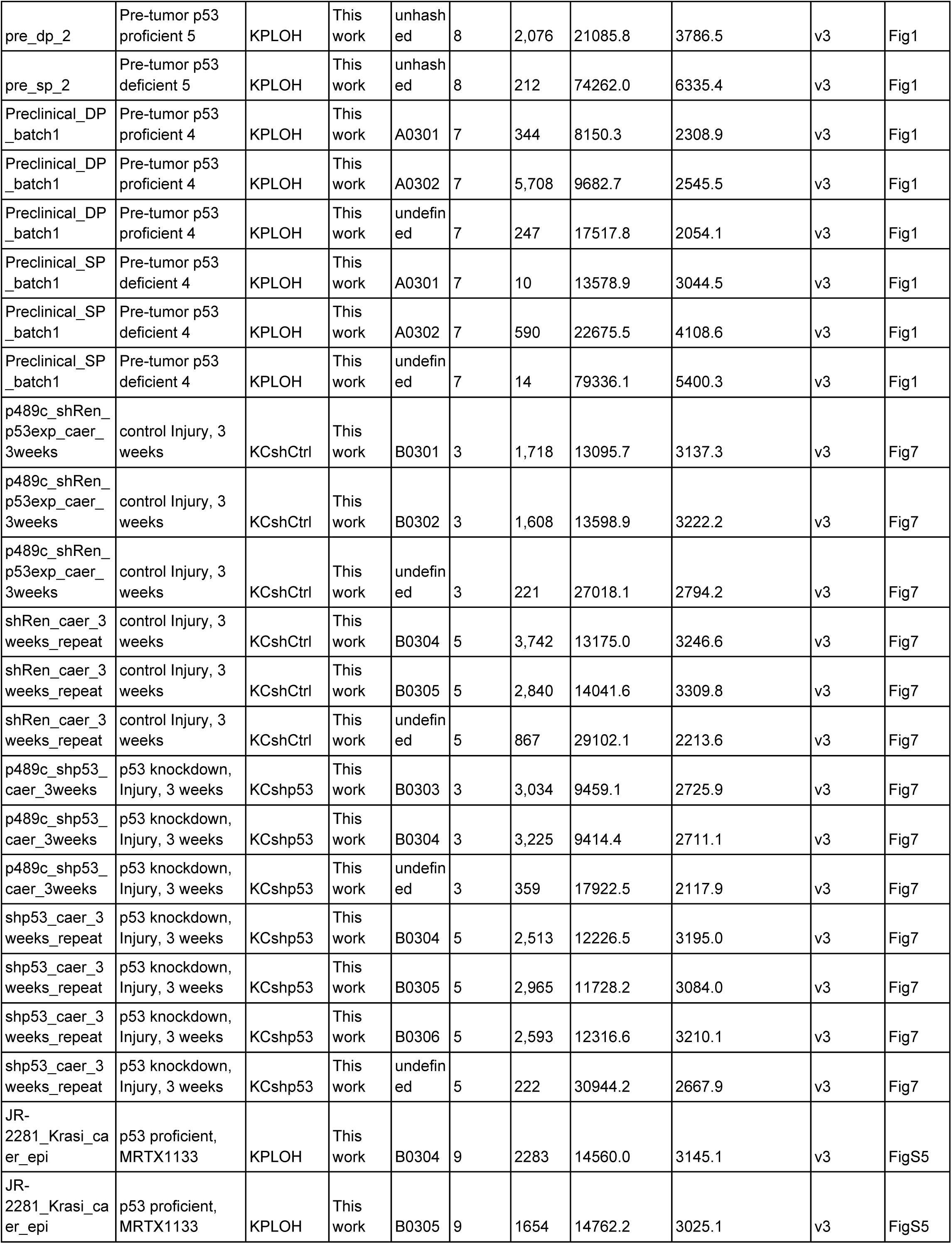
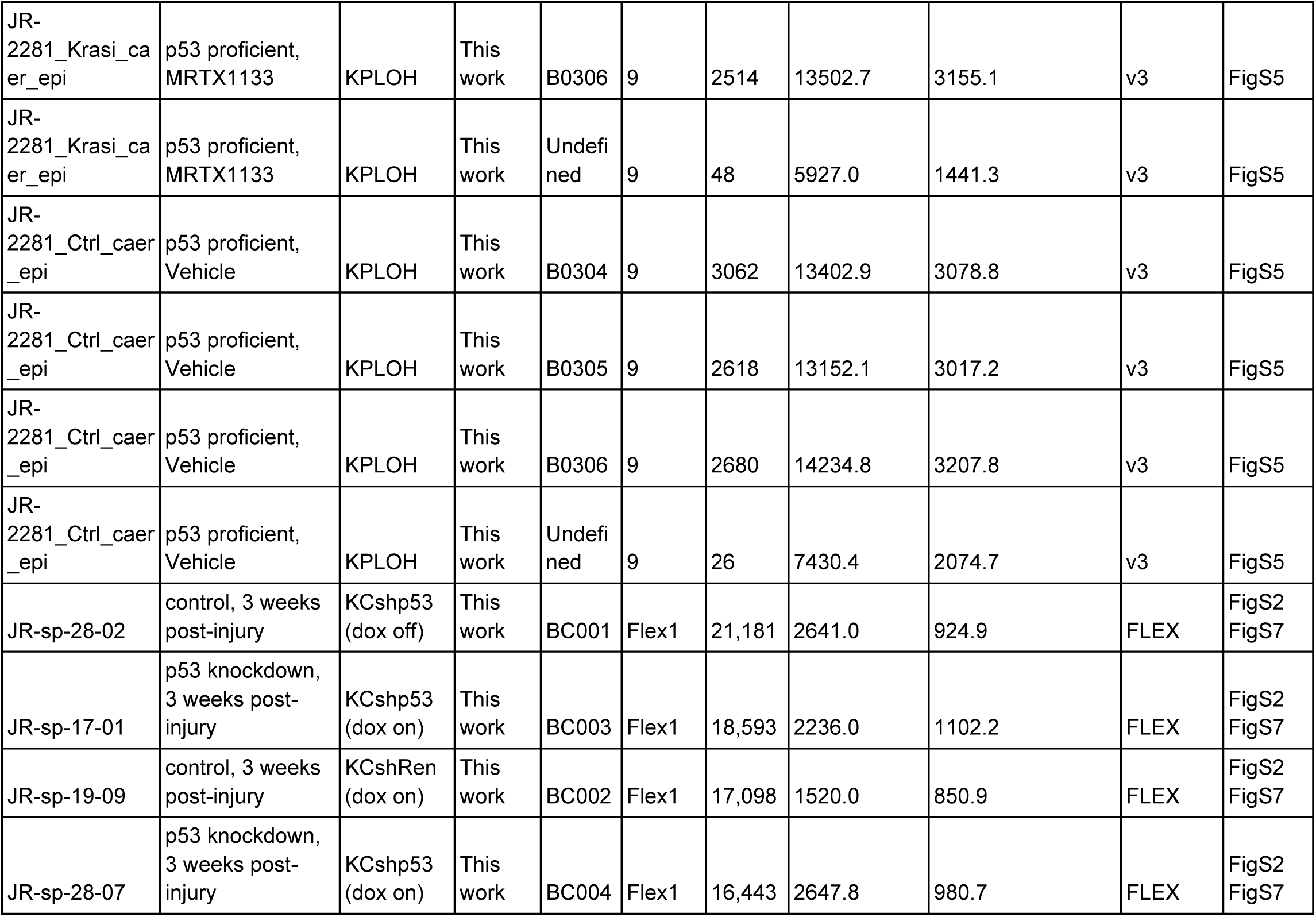
Metadata of scRNA-seq produced in this study, related to Figures 1, 3 and 7.

**Table S2.** Xenium sample information and embeddings, related to Figures 2, 3, 5, 6 and 7.

| xenium batch | slide id | slide name | mouse id | mouse strain | treatment | timepoint | Counts per cell | Num genes detected per cell |
| --- | --- | --- | --- | --- | --- | --- | --- | --- |
| JR-2885 | 0004279_Region_1 | JR0025 | JR-sp-04-03 | KCshCtrl | injury | 2d post-injury | 81.6 | 56.4 |
| JR-2885 | 0004279_Region_2 | JR0104_section 1 | JR-sp-12-23 | KPLOH | injury | 1d post-injury | 118.3 | 68.5 |
| JR-2885 | 0004279_Region_3 | JR0104_section 2 | JR-sp-12-24 | KPLOH | injury | 1d post-injury | 112.1 | 66.4 |
| JR-2885 | 0004329_Region_1 | JR0077 | JR-sp-09-05 | KCshCtrl | injury | 3w post-injury | 103 | 62.5 |
| JR-2885 | 0004329_Region_2 | JR0033 | JR-sp-04-02 | KCshCtrl | injury | 3w post-injury | 100.4 | 60.6 |
| JR-2885 | 0004329_Region_3 | JR0081 | JR-sp-09-03 | KCshp53 | injury | 3w post-injury | 112.4 | 65.1 |
| JR-3090 | 0011178_Region_1 | JR-sp-15-04 | JR-sp-15-04 | KPLOH | injury + MRTX1133 | 2d post inhibitor | 95.3 | 58.3 |
| JR-3090 | 0011178_Region_2 | JR-sp-15-09 | JR-sp-15-09 | KPLOH | injury + MRTX1133 | 2d post inhibitor | 97.4 | 58.8 |
| JR-3090 | 0011181_Region_1 | JR-sp-15-06 | JR-sp-15-06 | KPLOH | injury + MRTX1133 | 2d post inhibitor | 82.8 | 52.7 |
| JR-3090 | 0011181_Region_2 | JR-sp-15-02 | JR-sp-15-02 | KPLOH | injury + MRTX1133 | 2d post inhibitor | 96.2 | 58.6 |
| JR-2918 | 0015409_Region_1 | JR0079 | JR-sp-09-01 | KCshp53 | injury | 3w post-injury | 96.9 | 59.9 |
| JR-2918 | 0015409_Region_2 | JR0026 | JR-sp-04-07 | KCshp53 | injury | 2d post-injury | 93.7 | 59.4 |
| JR-2918 | 0015409_Region_3 | JR0080 | JR-sp-09-02 | KCshp53 | injury | 3w post-injury | 91.9 | 57.4 |
| JR-2918 | 0015409_Region_4 | JR0034 | JR-sp-04-10 | KCshp53 | injury | 3w post-injury | 97.1 | 59.2 |
| JR-2918 | 0015409_Region_5 | JR0027 | JR-sp-04-08 | KCshp53 | injury | 2d post-injury | 93.2 | 60.4 |
| JR-2918 | 0015410_Region_1 | DACE616 | DACE616 | KCshCtrl | injury | 2d post-injury | 92.5 | 59.5 |
| JR-2918 | 0015410_Region_2 | JR0078 | JR-sp-09-06 | KCshCtrl | injury | 3w post-injury | 84.4 | 53.5 |
| JR-2918 | 0015410_Region_3 | JR0024 | JR-sp-04-01 | KCshCtrl | injury | 2d post-injury | 81.3 | 55.7 |
| JR-2918 | 0015410_Region_4 | DACE617 | DACE617 | KCshCtrl | injury | 2d post-injury | 95 | 60.4 |
| JR-3083 | 0027845_Region_1 | JR0039 | JR-sp-06-09 | KCshCtrl | injury | 1d post-injury | 106.4 | 62.5 |
| JR-3083 | 0027845_Region_2 | JR0040 | JR-sp-06-10 | KCshCtrl | injury | 1d post-injury | 115.5 | 66.6 |
| JR-3083 | 0027845_Region_3 | JR-sp-15-05 | JR-sp-15-05 | KPLOH | injury + Vehicle | 2d post vehicle | 113.3 | 65 |
| JR-3083 | 0027846_Region_1 | JR0042 | JR-sp-06-04 | KCshp53 | injury | 1d post-injury | 84.5 | 54.1 |
| JR-3083 | 0027846_Region_2 | JR0041 | JR-sp-06-03 | KCshp53 | injury | 1d post-injury | 113.5 | 65.7 |
| JR-3083 | 0027846_Region_3 | JR-sp-15-07 | JR-sp-15-07 | KPLOH | injury + Vehicle | 2d post vehicle | 112 | 65.4 |
| JR-3177 | 0028094_Region_1 | JR-sp-19-09 | JR-sp-19-09 | KCshCtrl | injury | 3w post-injury | 88.5 | 57.5 |
| JR-3177 | 0028094_Region_2 | JR-sp-19-10 | JR-sp-19-10 | KCshCtrl | injury | 3w post-injury | 95.6 | 59.5 |
| JR-3433 | 0042536_Region_1 | JR-sp-28-25 | JR-sp-28-25 | KCshp53 | injury | 3w post-injury | 78 | 52.7 |
| JR-3433 | 0042536_Region_2 | JR-sp-28-08 | JR-sp-28-08 | KCshp53 | injury | 3w post-injury | 84.8 | 54.6 |
| JR-3311 | 0042569_Region_3 | JR0019 | JR-sp-02-17 | KPLOH | Spontaneous | pre-tumor | 60.2 | 43.4 |
| JR-3433 | 0042724_Region_1 | JR-sp-17-01 | JR-sp-17-01 | KCshp53 | injury | 3w post-injury | 80.6 | 52.5 |
| JR-3433 | 0042724_Region_2 | JR-sp-28-07 | JR-sp-28-07 | KCshp53 | injury | 3w post-injury | 85.4 | 55.4 |
| JR-3433 | 0042724_Region_3 | JR-sp-28-06 | JR-sp-28-06 | KCshp53 | injury | 3w post-injury | 89.2 | 57.1 |

## METHODS

## MOUSE MODELS

All animal experiments were performed in accordance with protocols approved by the Memorial Sloan Kettering Institutional Animal Care and Use Committee (approval number: 11-06-018). Mice were maintained under specific pathogen-free conditions, and provided with food and water *ad libitum*. In all experiments with PDAC models, tumors did not exceed a volume corresponding to 10% of body weight (typically 12–15-mm diameter). Mice were evaluated daily for signs of distress or end-point criteria, and immediately euthanized if they presented signs of cachexia, weight loss beyond 20% of initial weight or breathing difficulties, or if they developed tumors of 15 mm in diameter. Animals were housed on a 12 h light–12 h dark cycle under standard temperature (18–24 °C) and humidity (40–60%).

### Mouse model genetics

The KP^LOH^ model^11,28^ allows the identification and isolation of cells that undergo spontaneous *p53* loss of heterozygosity (LOH) during pancreatic cancer initiation. This model is derived from multi-allelic embryonic stem cells and harbors *Ptf1a-Cre*; *LSL-Kras^G12D^*; *p53^flox/WT^* alleles that predispose mice for spontaneous cancer development. Embryonic expression of *Ptf1a*-Cre in pancreatic epithelial progenitor cells leads to Cre-dependent excision of the LSL (lox-STOP-lox) cassette upstream of mutant *Kras^G12D^*, and deletion of one copy of *p53*. Oncogenic KRAS activity in the pancreatic epithelium leads to the formation of premalignant lesions^109^ and eventual PDAC development^11,28^ upon loss of the remaining wild-type copy of *p53* in premalignant cells (*p53*-LOH).

The KP^LOH^ model harbors fluorescent proteins that trace the lineage of cells that experienced Cre activity, as well as cells that have undergone *p53*-LOH. This model includes the *Rosa26-CAGGS-LSL-rtta-IRES-mKate2 (RIK)* allele^110^, in which Cre-dependent excision of the LSL cassette leads to constitutive polycistronic production of the reverse tetracycline transcriptional activator (rtTA3) and mKate2. Both proteins serve as proxies for oncogenic *Kras^G12D^* activation in epithelial cells (detected through mKate2 immunofluorescence), single molecule fluorescence in situ hybridization (smFISH) or transcriptomics. Furthermore, upon doxycycline administration, rtTA3 expression allows selective induction of transgenes downstream of a promoter harboring the tetracycline-regulated element. This genetic configuration thus allows both detection and perturbation of premalignant cells *in vivo*.

Lastly, the KP^LOH^ model harbors the doxycycline-inducible *TRE-GFP-shRen.713* allele, which produces GFP and a short-hairpin RNA (shRNA) targeting *Renilla* luciferase (shRen) upon doxycycline administration. We have used this allele extensively as a non-targeting negative control^40^ to account for non-specific effects of shRNA-expression and interaction with the RNAi machinery in a cell. In the context of the KP^LOH^ model, this allele serves the unique purpose of reporting for the genetic status of *p53 in vivo*. The TRE-GFP-shRen allele is located in the *Col1a1* safe-harbor locus in mouse chromosome 11, in cis with the single *p53*^WT^ allele (due to how we designed our breeding scheme). Given the selective pressure for homozygous *p53* loss during PDAC development, and the fact that *p53*-LOH events most frequently occur through whole chromosome or large segmental deletion events^28^, loss of GFP serves as a proxy for *p53*-LOH in this model.

All cohorts in this study were derived from multiallelic mouse embryonic stem cells (mESCs) harboring the genetic configuration described above (KPfCRC shRen clone YMZ). While this model uniquely allows tracing and isolation of cells that underwent spontaneous *p53*-LOH events *in vivo*, we have also used these mESCs to study *p53*-proficient cells in the premalignant pancreas due to their high efficiency in generating experimental mice.

KC^shRNA^ models harbor the same genetic configuration as the KP^LOH^ model, except that they have *p53*^WT/WT^ alleles. We use two variants of this model: KC^shCtrl^ harbors the *TRE-GFP-shRen.713* allele, serving as a non-targeting negative control (shRen), and KC^shp53^ harbors the *TRE-GFP-shp53.1224* allele^40^ targeting shp53. We used these models to investigate the consequences of inducible *p53* knockdown *in vivo*. All KC^shRNA^ mice in this study were generated using multiallelic mESCs (KC^shCtrl^: clone p48-9c shRen.713 c2; KC^shp53^: clone p48-9c shp53.1224 #6)^41^.

The identities of ESCs and ESC-derived mice were authenticated by genomic PCR using a common Col1a1 primer paired with an shRNA-specific primer:

- Col1a1: 5’-CACCCTGAAAACTTTGCCCC-3’

- shRen.713: 5’-GTATAGATAAGCATTATAATTCCTA-3’ (∼250 bp band)

- shp53.1224: 5’-TGTATTACACATGTACTTGTAGTGG-3’ (∼210 bp band)

The presence of the RIK allele in mESCs was confirmed using PCR with the following primers:

- 5’-GGTGAGCGAGCTGATTAAGG-3’

- 5’-TTTTGCTGCCGTACATGAAG-3’ (∼200 bp band)

In addition, we confirmed shp53 and shRen expression at the single-cell level by aligning reads to the unique sequences that distinguish TRE-GFP-shp53 and TRE-GFP-shRen alleles:

>TGM_shRen_unique

TGCTGTTGACAGTGAGCGCAGGAATTATAATGCTTATCTATAGTGAAG CCACAGATGTATAGATAAGCATTATAATTCCTATGCCTACTGCCTCGG

>TGM_shp53_unique

TGCTGTTGACAGTGAGCGCCCACTACAAGTACATGTGTAATAGTGAAG CCACAGATGTATTACACATGTACTTGTAGTGGATGCCTACTGCCTCGG

### Cohort generation

ESC-derived chimeric male mice were generated by injecting KP^LOH^, KC^shCtrl^ or KC^shp53^ backgrounds at the 8-cell or blastocyst stage, as previously described^40^, enabling the synchronous creation of large cohorts of mice bearing all alleles for modeling PDAC initiation and progression. Cohorts were generated by the Mouse Genetics Core Facility at Memorial Sloan Kettering Cancer Center (MSK) or the Rodent Genetic Engineering Core at New York University. Only mice with coat-color chimaerism of over 95% were included for experiments. Due to the sex of our established mESCs lines, the majority of mice derived from them are male. As has been noted ^111^, cohorts from male mESCs can also produce female mice due to Y chromosome loss in culture. Our studies were largely conducted on male mouse cohorts, and we used female mice to complement some of our analyses (e.g., generation of one organoid line). Thus, we lack information regarding the influence of sex on our in vivo studies, which is a limit to the generalizability of our results across sexes.

### shRNA induction

To induce shRNA expression, KC^shCtrl^ and KC^shp53^ mice were switched to a doxycycline diet (200 mg kg^-1^, Harlan Teklad) at 4–5 weeks of age, one week before inducing pancreatitis. KP^LOH^ mice were switched to doxycycline diet at 4–5 weeks of age to allow doxycycline dependent induction of the GFP transgene that reports the presence or absence of the wild-type *p53* allele in this model.

For KC^shp53^ collection cohort 4 (**Data S6**), a subset of mice were switched to doxycycline diet and the remaining animals were fed normal chow. *p53* knockdown led to an increase in progenitor-like cells and expansion of their niche in this comparison. These results not only corroborate previous findings, but also provide an orthogonal negative control, as they show that differences in the abundance of progenitor-like cells track with *p53* knockdown, as opposed to the specific mESC strain used to generate cohorts.

KP^LOH^ mice from batches 2, 7 and 8 (see **Table S1** for details on sample metadata) were administered 625 mg kg^-1^ doxycycline diet (Harlan Teklad) following prior practices in our lab^28^. The remaining cohorts were treated with low dose doxycycline (200 mg kg^-1^) to minimize the potential effects of antibiotic treatment on the microbiome. No differences in the spectrum or dynamics of premalignant states were observed as a function of doxycycline dose.

## ORGANOID MODELS

### Generation of premalignant pancreatic organoids

Pancreatic epithelial cells were FACS-sorted from 5–6-week-old pre-tumor KP^LOH^ mice. Three organoid lines were established: two of them derived from male mice, and one from a female mouse. Mice were placed on doxycycline feed prior to FACS sorting to activate the GFP reporter for wild-type *p53*. Pancreata were dissociated into single cells (see ‘Tissue dissociation for single-cell analyses’), and mKate+/GFP+ (i.e. Kras-mutant, p53-proficient) cells were sorted and seeded in Growth Factor-Reduced Matrigel (Corning). Once matrigel solidified, organoids were covered in complete organoid medium adapted from Huch et al., 2013^112^. Complete organoid medium was composed of Advanced DMEM/F12 (Thermo Fisher) supplemented with 1% GlutaMAX (Thermo Fisher), 1% penicillin:streptomycin (Fisher), B27 (Thermo Fisher), 500 nM A83-01 (Tocris), 50 ng/mL EGF (Invitrogen), 10 nM Gastrin (Sigma-Aldrich), 100 ng/mL Noggin (Peprotech), 100 ng/mL FGF10 (Peprotech), 500 ng/mL R-spondin (Peprotech), 1.25 mM N-acetylcysteine (Sigma-Aldrich), and 10 mM Nicotinamide (Sigma). Medium was replaced every 2–3 days. For initial seeding and up until the first passage, 10 nM Y-27632 (Sigma-Aldrich) and 50 ug/mL Normocin (Fisher) were also added to the medium. Organoids were authenticated through genotyping, and were monitored through passaging to ensure the presence of mKate2 and GFP fluorescence.

## IN VIVO TREATMENTS

### Injury-induced pancreatitis

To assess the spectrum and dynamics of premalignant states and tissue remodeling events in the in the context of oncogenic Kras activation in the premalignant epithelium, we subjected 5–6-week-old male mice with intraperitoneal injections of 80 μg kg^-1^ of caerulein (Bachem) every 8 h for two consecutive days (16 injections total), as previously described^26^. Caerulein dose was adjusted to body weight at the beginning of each day of treatment. We harvested the pancreata at two phases of the injury response: an acute phase, corresponding to the peak of inflammation (days 1 and 2 after the 9th caerulein injection), or a long-term response (3 weeks post-caerulein treatment). All our single cell and spatial datasets on injury-induced pancreatitis have at least 3 replicates per condition. We did not use a statistical criteria for sample size estimation.

### Oncogenic KRAS inhibition

The KRAS^G12D^-specific small molecule inhibitor MRTX1133^62^ allowed us to interrogate the consequences of acute removal of Kras signaling in premalignant cells without directly affecting the tumor microenvironment (**Figs. 6** and **S9**). All our single cell and spatial datasets on oncogenic KRAS inhibition have at least 3 replicates per condition, with the exception of vehicle control tissue profiled with Xenium, for which we interrogated two biological replicates. We did not use a statistical criteria for sample size estimation.

#### Formulation for *in vivo* use

We formulated MRTX1133 for *in vivo* use, as previously described^94^. To prepare a vehicle solution of 10% Captisol, 50 mM citrate buffer pH 5.0e for drug administration, we mixed 20% w/v Captisol (MedChemExpress, HY-17031) in sterile water with 100 mM citrate buffer pH 5.0 (Teknova, Q2443) in a 1:1 ratio. MRTX1133 stock solution was prepared by dissolving powder in vehicle solution at 3 mg mL^-1^. We stored vehicle and MRTX1133 formulations at 4°C protected from light for up to 1 week.

#### Dosing

We fed mice with MRTX1133 at the maximum tolerated dose of 30 mg kg^-1^ through intraperitoneal injection (200 μL for a 20-g mouse) or an equivalent volume of vehicle based on weight. Dosing was twice a day, with an inter-dose interval of 10–12 h. We randomized MRTX1133 and vehicle-treated mice to control for differences in the social structure of individual cages, as well as inter-cage heterogeneity in average mouse size.

#### Short-term consequences of oncogenic KRAS inhibition

We used 5-week-old male KP^LOH^ mice generated through mESC injections as experimental cohorts. For mice profiled with scRNA-seq, we started dosing of MRTX1133 or vehicle concurrently with caerulein treatment, euthanizing mice within 4 h of the last MRTX1133 or vehicle dose (fifth dose, 2 days after the first dose). For mice profiled with Xenium spatial transcriptomics, we started MRTX1133 or vehicle dosing 2 days after the first caerulein dose, euthanizing mice 2 days after the first dose of MRTX1133 or vehicle. These two experimental protocols aimed to assay the role of oncogenic KRAS signaling in inducing or maintaining the progenitor-like state upon acute pancreatitis. In practice, both protocols had equivalent outcomes in terms of the spectrum of premalignant lesions at end-point: depletion of progenitor-like, gastric pit-like and gastric chief-like cells, as well as shifts in the state of acinar-to-ductal metaplasia (ADM) cells, with progenitor-like cells being the subpopulation with strongest dependency on persistent oncogenic KRAS (**Figure 6C** and **S9c**).

#### Long-term consequences of oncogenic KRAS inhibition

We used 5-week-old male KP^LOH^ mice generated through mESC injections as experimental cohorts. One week prior to starting the experiment, we transferred mice to doxycycline diet (200 ppm) to allow interrogation of *p53* loss at the end point. We treated mice with acute pancreatitis as described above (Injury-induced pancreatitis). Two days after the first dose of caerulein, we dosed mice with the maximum tolerable dose of MRTX1133 (30 mg kg^-1^) or vehicle 2 times a day for 3 days, separating doses by no less than 10 h. We kept mice on doxycycline diet, monitoring weekly for signs of ill health and palpating for tumors. Mice that had to be euthanized for health problems (e.g., hydrocephaly, head tilts, distended abdomen due to feces accumulation), but lacked macroscopic PDAC at end point, as determined by gross histological examination, were excluded from the analysis. These included two MRTX1133-treated long-term survivor mice (8.8 and 14 months) that had to be euthanized for reasons other than PDAC. We conducted this experiment independently in two cohorts of KP^LOH^ mice born within 20 days of each other, pooling survival data. Statistical analysis for differences in survival of KP^LOH^ mice as a function of treatment was conducted using the lifelines python package (v0.30.0) (**Figure 6H**).

### Macrophage depletion

To assess the contribution of macrophages to premalignant cell state dynamics, we subjected 4–8-week old mice to twice weekly intraperitoneal injections of 200 μg anti-CSF1R-depleting antibody (Bio X Cell, BE0213) or 200 μg immunoglobulin G2a (IgG2a) isotype (Bio X Cell, BE0090). After one week of treatment, mice were subjected to injury-induced pancreatitis as previously described. Mice were dosed for an additional 3 weeks after pancreatitis with anti-CSF1R or IgG2a antibodies.

## SAMPLE COLLECTION

**Table S1** provides details of scRNA-seq samples collected for this study.

### Experimental endpoints

Pre-tumor stage male KP^LOH^ mice were euthanized at 3–4 months of age. Lack of a macroscopic tumor mass was assessed by palpation before euthanasia, and confirmed by gross histology upon dissection. Tumor stage KP^LOH^ mice were euthanized upon confirmation of the presence of a macroscopic tumor mass by palpation; two animals at 3 months of age, and one at 8 months. Mice subjected to acute pancreatitis in time-course and *p53*-perturbation cohorts were euthanized at 1 day, 2 days or 3 weeks after the second day of the pancreatitis protocol. Mice treated with MRTX1133 or vehicle were euthanized 2 days after the first treatment dose.

### Tissue dissociation for single-cell analyses

For scRNA-seq and bulk RNA-seq collection, we isolated lineage-traced (mKate2+/GFP+ or mKate2+/GFP-) epithelial cells from pancreatic tissues from KP^LOH^, or KC^shRNA^ mice by FACS sorting, as previously described^7,26^. Specifically:

1. Pancreata were finely chopped with scissors and incubated in digestion buffer containing 1 mg mL^-1^ collagenase V (Sigma-Aldrich, C9263), 2 U mL^-1^ Dispase (Life Technologies, 17105041) dissolved in HBSS with Mg^2+^ and Ca^2+^ (Thermo Fisher Scientific, 14025076) supplemented with 0.1 mg mL^-1^ DNase I (Sigma, DN25-100MG) and 0.1 mg mL^-1^ soybean trypsin inhibitor (STI) (Sigma, T9003), in gentleMACS C Tubes (Miltenyi Biotec) for 42 min at 37°C using the gentleMACS Octo Dissociator.
2. Digested samples were washed with PBS and further digested with a 0.05% solution of Trypsin-EDTA (Thermo Fisher Scientific, 15400054) diluted in PBS for 5 min at 37°C. Trypsin digestion was neutralized with FACS buffer (10 mM EGTA and 2% FBS in PBS) containing DNase I and STI.
3. Samples were washed in FACS buffer containing DNAse and STI, and filtered through a 100-μm strainer.
4. Samples were blocked with anti-mouse CD16/CD32 with Fcblock (BD Biosciences, Cat# 553141; Clone 2.4G2) for 7 min at 4°C, followed by incubation with APC-conjugated CD45 antibody (Biolegend, Cat# 103111; Clone 30-F11, 1:200, 10-min incubation).
5. Cells were washed once in FACS buffer containing DNase I and STI, filtered through a 40-μm strainer and resuspended in FACS buffer containing DNase I and STI and 300 nM DAPI as a live-cell marker. Cells were sorted on BD FACSAria I or BD FACSAria III (Becton Dickinson) for mKate2+/GFP+ (*Kras*^G12D^+ *p53*-proficient epithelial cells) or mKate2+/GFP- (*Kras*^G12D^+ *p53*-deficient epithelial cells), excluding DAPI+ and CD45+ cells. FACS-sorted cells were collected in 2% FBS in PBS (see **Data S1.1A** for representative FACS gates for isolating epithelial cells from a pre-tumor stage KP^LOH^ mouse).
6. For scRNA-seq, cells were resuspended at 1000 cells μL^-1^ in 0.04% BSA 1X PBS solution with RNase inhibitor (Thermo Fisher Scientific, AM2684; 1 U μL^-1^ or 1:40 dilution from stock). In the case of pre-tumor *p53*-deficient cell isolation, the low frequency of this subpopulation (200–2000 cells per mouse) limited our ability to resuspend and process sorted cells directly from the sorter. To allow downstream processing of this rare and important cell population, we spiked-in CD45+ cells isolated from the same mouse to reach a threshold of 30,000 cells, resuspending in a final volume of 30 μL for downstream processing.

For *p53* perturbation and acute oncogenic Kras inhibition experiments, we modified this isolation protocol to accommodate pooling of biological replicates in the same encapsulation and sequencing runs via cell hashing, minimizing both costs and batch effects. Following step 2 above:

1. For every sample that would be subjected to cell hashing, we used DNase-free buffers from this point on, as DNase could hamper our ability to recover DNA barcodes.
2. Samples were resuspended in 1 mL ACK-lysis buffer and incubated for 5 min at room temperature to deplete red blood cells, and washed the ACK lysis buffer with 20 mL HBSS.
3. We blocked samples with TruStain Fc block Plus (Biolegend, 156603; clone S17011E, 1:100) for 7 min at 4°C, followed by incubation with a sample-specific TotalSeq cell hashing antibody (Biolegend, 155832, 155833, 155835, 155837, 155839, 155841; clones M1/42; 30-F11). We incubated cell hashing antibodies at a 1:50 dilution for 30 min.

1. We washed samples 3 times with DNAse-free FACS buffer + STI, followed by filtering through a 40-μm strainer and FACS-based isolation of mKate2+/GFP+ cells (*Kras*^G12D^+ pancreatic epithelial cells expressing shp53 or shCtrl).
2. To prepare cells for scRNA-seq, we pooled samples from the same experimental conditions into the same tube, and resuspended cells at a 1000 cells μL^-1^. This strategy ensured our ability to interpret differences between conditions even if deconvolution of biological replicates failed. In practice, we didn’t experience problems with downstream deconvolution of biological replicates in these data.

### Preparation of tissues for histology

For immunofluorescence and Xenium-based analyses, tissues were fixed overnight in 10% neutral buffered formalin (Richard-Allan Scientific) and embedded in paraffin. Formalin fixed paraffin embedded (FFPE) blocks were stored at room temperature, or more recently at 4°C as we started profiling RNA from these tissues using spatial transcriptomic technologies.

For smFISH-based analyses, we followed the fixation protocol by Farack and Itzkovitz^113^. Specifically, we fixed tissue in 4% PFA (Fisher / Electron Microscopy Sciences, 15710) 1X PBS at 4°C for 3 h, followed by overnight incubation in 4% PFA, 1X PBS, 30% w/v sucrose solution, verifying that tissues sank to the bottom of the tube before further processing. We washed tissues with 1X PBS and thoroughly dried them with a Kimwipe before OCT embedding. We incubated tissues for 30 min to 1 h in OCT at 4°C, since we observed that this decreases the likelihood of tissue detachment during sectioning as compared to immediate freezing. Lastly, we completed embedding by placing tissues in a mold, fully covering with OCT, and placing on dry ice for freezing. We stored frozen OCT blocks at −80°C.

## SINGLE-CELL RNA SEQUENCING

### Fresh dissociated samples

Cells were resuspended in 1X PBS and BSA (0.04%) and checked for viability using 0.2% (w/v) Trypan Blue staining (Countess II). All sequencing experiments were performed on samples with a minimum of 80% viable cells. Single-cell encapsulation and scRNA-seq library prep of FACS-sorted cell suspensions was performed on the Chromium instrument (10x Genomics) following the user manual (Reagent Kit 3’ v2 or v3). Each sample loaded onto the cartridge contained approximately 5,000 cells (non-hashed samples) or 15,000 cells (hashed samples) at a final dilution of ∼500 cells μl^-1^. Transcriptomes of encapsulated cells were barcoded during reverse transcription and the resulting cDNA was purified with DynaBeads, followed by amplification per the user manual. Next, the PCR-amplified product was fragmented, A-tailed, purified with 1.2X SPRI beads, ligated to sequencing adapters and indexed by PCR. Indexed DNA libraries were double-size purified (0.6–0.8X) with SPRI beads and sequenced on an Illumina sequencer (R1 – 26 cycles, i7 – 8 cycles, R2 – 70 cycles or higher) to a depth of >50 million reads per sample (>13,000 reads per cell) at MSK’s Integrated Genomics Operation Core Facility.

### Dissociated nuclei from FFPE samples

FFPE samples were preprocessed using a prototype Singulator™ system. Each sample was automatically processed in a NIC+™ cartridge (S2 Genomics, 100-215-389) through two 10-min deparaffinization steps using Deparaffinization Reagent (S2 Genomics), followed by rehydration through successive 1 mL ethanol washes (100%, 100%, 70%, 50%, and 30%). This was followed by two PBS washes. The sample was then centrifuged at 1,000 *g* for 3 min and resuspended in 0.5 mL of Nuclei Isolation Reagent (NIR, S2 Genomics, 100-063-396) containing 0.1 U μL^-1^ RNase inhibitor (Protector™, Millipore Sigma, 3335399001). All subsequent solutions contained RNase inhibitor at the same concentration. The sample was dissociated into single nuclei in a second NIC+ cartridge using “FFPE Nuclei Isolation” protocol, using 0.5 mL of NIR for 12 min of lysis, followed by a 2-mL wash with Nuclei Storage Reagent (NSR, S2 Genomics, #100-063-405). The single-nucleus suspension was centrifuged at 500 *g* for 5 min, resuspended in NSR, and counted using 0.2% (w/v) Trypan Blue staining on a Countess II instrument. This was followed by a second centrifugation at 850 *g* for 5 min.

Nuclei were then resuspended in 1 mL of Fixation Buffer (4% formaldehyde in 1× Fix & Perm Buffer, 10x Genomics, PN-2000517) and incubated at 4°C for 16–24 h. To stop the fixation, nuclei were centrifuged at 850 *g* for 5 min at room temperature and quenched with 1 mL of Quenching Buffer (1× Quench Buffer, 10x Genomics, PN-2000516). Fixed nuclei were then stained with 1 μg mL^-1^ DAPI and sorted for DAPI-positive nuclei.

Up to 300,000 nuclei were processed per hybridization according to 10x Genomics recommendations. Each hybridization was performed in 40 μL of hybridization mix, containing 10 μL of Mouse WTA probes (10x Genomics, PN-2001275) and 2.5 μL of custom probes targeting eGFP and mKate2 for a final concentration of 2 nM per probe. Custom probes were designed following the 10x Genomics technical note on probe design, with particular attention to GC content (see **Data S7** for probe sequences). Hybridizations were carried out at 42°C for 16–24 h.

Following hybridization, samples were diluted in Post-Hybridization Wash Buffer and counted. For each experiment, an equal number of nuclei from each hybridization reaction was pooled to ensure equal sample representation. The pooled nuclei were then washed four times in Post-Hybridization Wash Buffer for 10 min at 42°C. After washing, nuclei were resuspended in Post-Hybridization Resuspension Buffer, filtered through a 30 μm Miltenyi Biotec filter, and counted to determine the appropriate volume for loading onto the Chromium X instrument.

GEM encapsulation was performed following the 10x Genomics Flex GEM-X (PN-1000782) protocol, using their guidelines for cell and reagent volumes per well based on the desired cell recovery. After loading the Chip FX and running it on the Chromium X, GEMs were recovered and processed according to the manufacturer’s instructions. Following GEM processing, the resulting product was pre-amplified and indexed to generate the sequencing library. All libraries were sequenced on an Illumina NovaSeq X+ (R1 – 28 cycles, i5 – 10 cycles, i7 – 10 cycles, R2 – 90 cycles) using standard dual indexing and demultiplexing. Raw BCL files were processed with Cell Ranger (9.0.0), and the resulting FASTQ files were quantified using a custom probe set reference for the mouse genome (GRCm39) within the Cell Ranger pipeline.

**Table S1** provides details of the dissociated samples collected as part of this study.

## SINGLE-CELL RNA + ATAC MULTIOME SEQUENCING

ATAC and gene expression libraries were prepared according to the Chromium Next GEM Single Cell Multiome ATAC + Gene Expression workflow, following 10x Genomics guidelines for ATAC, gene expression, and sequencing preparation. Cells were pelleted (300 *g*, 5 min, 4°C), permeabilized in chilled lysis buffer, incubated on ice for 5 min, and washed in chilled buffer containing RNase inhibitor (Protector, Roche). After centrifugation (500 *g*, 5 min, 4°C), nuclei were resuspended in Nuclei Buffer (10x Genomics), counted, and adjusted to 3,000 nuclei µL^-1^.

For each reaction, up to 15,400 nuclei were diluted in isotonic Tagmentation Buffer with RNase inhibitor and mixed with ATAC Buffer B and ATAC Enzyme B (10x Genomics) for transposition (37°C, 60 min). Samples were loaded onto a Chromium Next GEM Chip J with barcoding reagents, gel beads, and partitioning oil (10x Genomics) and processed on the Chromium Controller to generate GEMs. Reverse transcription and barcoding proceeded at 37°C for 45 min and 25°C for 30 min. Later, GEMs were broken, and the aqueous phase was purified using Dynabeads MyOne SILANE (10x Genomics) and SPRIselect magnetic bead cleanups (Beckman Coulter). Barcoded ATAC and cDNA fragments were pre-amplified and purified with SPRIselect.

ATAC libraries were constructed using sample indexing PCR with 10x Genomics reagents, followed by dual-sided SPRI size selection. cDNA was amplified, cleaned, fragmented, end-repaired, A-tailed, and ligated using 10x Genomics reagents, with SPRI-based purifications throughout. Final gene expression libraries were indexed using 10x Genomics dual-index reagents and purified with SPRIselect. Sequencing was performed according to 10x Genomics recommendations on an Illumina NovaSeq 6000.

## SPATIAL PROFILING

We performed IF to quantify up to 4 proteins in FFPE (**Figs S1I, S2C, S2F, 5E, 6B, 6E, S5A, S5J, 7B, 7F**), the Lunaphore COMET multiplex IF platform to quantify 12 protein markers in FFPE (**Figs S2A and S5D**), and the Leica Cell DIVE multiplex immunofluorescence platform to quantify 50 protein markers in FFPE human tissue (**Fig. 4C**) and 6 protein markers in mouse tissue (**Fig. 3H**). In addition, we used our custom smFISH platform to stain up to 5 mRNAs in fixed frozen mouse tissue (**Figs 1H, 2C and Data S1.2K**). Lastly, we used the 10x Xenium spatial transcriptomics platform to quantify 480 mRNAs in FFPE mouse tissue used throughout the manuscript.

**Data S6** contains source data and sample metadata of tissues analyzed with immunofluorescence. **Table S2** describes details of the 10x Xenium spatial transcriptomics samples we collected for this study.

### Immunofluorescence

Immunofluorescence (IF) was conducted on 5-μm sections of FFPE blocks. Following deparaffinization and antigen retrieval (citrate buffer pH 6.0, Fisher / Vector Biolabs, H-3300-250), slides were blocked with 5% BSA 1X PBS for 1 h at room temperature, followed by overnight incubation with primary antibodies. Following primary incubation, we washed slides 3 times for 10 min with 1X PBS, and incubated with secondary antibodies diluted in blocking buffer for 1h. We washed slides for 10 min with 1X PBS 1 ug mL^-1^ DAPI, followed by two additional washes and mounting. Images were imaged with a Nikon T2i Eclipse system equipped with a 20X Plan APO objective (Nikon, MRD00205) equipped and an ORCA-FusionBT sCMOS camera. For quantification of HMGA2, VIM and TNC, we collected full tissue scans using the Nikon Elements image acquisition software.

For quantification of p53 levels, we stained adjacent tissue sections for the progenitor state marker MSN or p53, as well as GFP as a proxy for *Kras*^G12D^+/*p53*-proficient cells. We selected and acquired fields of view based on the presence of MSN+ and MSN-lesions and subsequently acquired the corresponding regions of interest in the p53-stained slide. We acquired images using a 20X Plan APO objective and the Crest X-Light V2 LFOV25 Spinning Disk Confocal attached to our Nikon T2i Eclipse microscope, collecting fields of view of 2000 x 2000 px.

We used the following primary antibodies: GFP (Abcam, ab13970; RRID:AB_300798, 1:1000), RFP (Evrogen, AB233; RRID:AB_2571743, 1:1000), HMGA2 (Cell Signaling Technology, 8179S; clone D1A7, RRID:AB_11178942, 1:500), Moesin (Proteintech, 26053-1-AP; RRID:AB_2880353, 1:100), E-cadherin (BD Biosciences, 610181; RRID:AB_397580, 1:500), Vimentin (Cell Signaling Technology 5741; RRID:AB_10695459, 1:500), mKate2 (generated in-house, #4007; rat isotype, 1:250), p19 (SantaCruz, sc-32748; RRID:AB_628071 1:100), Tenascin-C (R&D systems, MAB2138; RRID:AB_2203818, 1:250), p53 (Leica Biosystems, p53-CM5P; RRID:AB_2744683, 1:250), Phospho-p44/42 MAPK (Erk1/2) (Thr202/Tyr204) (Cell Signaling Technology, 9101; 1:250). We used the following secondary antibodies as part of this study: Donkey Anti-Mouse IgG Alexa Fluor 750 (Abcam, ab175738; 1:250), Donkey anti Rabbit IgG Alexa Fluor 555 (Invitrogen, A31572; 1:500), Donkey anti-Rat IgG Antibody, Alexa Fluor 647 (Thermo Fisher Scientific, A78947; 1:500), Donkey anti-Chicken IgY Alexa Fluor 488 (Thermo Fisher Scientific, A78948; 1:1000), Donkey Anti-Chicken IgY Alexa Fluor 647 (Sigma-Aldrich, AP194SA6; 1:1000), Donkey anti Mouse IgG Alexa Fluor 488 (Thermo Fisher Scientific, A21202; 1:1000), Donkey anti-Rabbit IgG Alexa Fluor Plus 647 (Invitrogen; A32795, 1:500), Donkey anti-Rat IgG Alexa Fluor Plus 555 (Invitrogen, A48270; 1:500).

### Multiplexed Immunofluorescence using Lunaphore COMET

The Lunaphore COMET platform allowed us to probe for multiple markers of the progenitor-like state in the same tissue slide, overcoming limitations of isotype incompatibility between markers (e.g., MSN and HMGA2 antibodies are both derived from rabbit hosts). Tissue sections (5 µm) were trimmed from a FFPE block and placed at the center of a clean glass slide. The slide was air dried and baked at 42°C for 3 h and stored in a desiccator. Epredia PT Module was used to deparaffinize and retrieve epitopes (Epredia Dewax and HIER Buffer L and Epredia Dewax and HIER Buffer H). The slide was then washed twice with 1X Multistaining buffer (BU06) and loaded onto the COMET. Appropriate volumes of primary antibodies, secondary antibodies, 5 µg mL^-1^ DAPI (Thermo Fisher Scientific, D3571), Multistaining buffer, Quenching buffer (BU08-L), Imaging buffer (BU09), and Elution buffer (BU07-L) were freshly made and loaded into the fluidics compartment of the instrument. Fields of view (FOVs) of 12 mm x 12 mm were captured in a tiled fashion, only where the tissue was auto detected. The primary antibodies were used at the following dilutions: 1:1000 GFP (Abcam, ab13970), 1:300 HMGA2 (CST, 8179S), 1:100 Moesin (ProteinTech, 26053-1-AP), 1:100 p19 (SantaCruz, sc-32748), 1:50 cleaved caspase3 (CST, 9664S), 1:100 Galectin-4 (R&D systems, AF2128), 1:200 Annexin A10 (Abcam, ab213656), 1:1000 E-cadherin (BD, 610181), 1:200 Phospho-p44/42 MAPK (Erk1/2) (CST, 9101), 1:200 Vimentin (CST, 5741), 1:500 CPA1 (R&D systems, AF27765), 1:100 RFP (Genscript, Custom ID 4007). The secondary antibodies were used at the following dilutions: 1: 100 Donkey anti-Rabbit AlexaFluor Plus 555 (Thermo Fisher Scientific, A32794), 1:200 Donkey anti-Rabbit AlexaFluor Plus 647 (Thermo Fisher Scientific, A32795), 1:200 Donkey anti-Rat AlexaFluor Plus 647 (Thermo Fisher Scientific, A48272), 1:200 Goat anti-Chicken AlexaFluor Plus 647 (Thermo Fisher Scientific, A32933), 1:100 Donkey anti-Mouse Plus 555 (Thermo Fisher Scientific, A32773), 1:200 Donkey anti-Goat Plus 647 (Thermo Fisher Scientific, A32849).

### Multiplexed Immunofluorescence using Leica Cell DIVE

We used the Leica Cell DIVE imaging system to conduct multi-IF experiments through cycles of staining, imaging and bleaching of fluorescent stains. This imaging system allowed us to probe for multiple microenvironment markers in the same tissue section while bypassing limitations of isotype incompatibilities, and to leverage the computational removal of autofluorescence. We imaged 5-μm FFPE sections following the manufacturer’s protocol. Briefly, after a 2-step antigen retrieval process, slides were blocked with 3% BSA, stained with DAPI, and imaged unstained to acquire background autofluorescence (AF). Samples were then stained and imaged using DAPI, Cy3, Cy5, and FITC channels on the Cell DIVE instrument with Cell DIVE image acquisition and processing software. Each FOV was imaged in each staining round, followed by AF removal, registration with baseline DAPI, and stitching. Unconjugated primary antibodies were used in the first staining round, followed by secondary antibody staining: GFP (Abcam, ab13970; RRID:AB_300798, 1:1000), HMGA2 (Cell Signaling Technology, 8179S; clone D1A7, RRID:AB_11178942, 1:500) and Tenascin-C (R&D systems, MAB2138; RRID:AB_2203818, 1:250). After imaging, dye inactivation was performed using 0.1 M Na_2_CO_3_ 3% H_2_O_2_ solution for 15 min at room temperature, followed by 1 h blocking with rabbit serum (Sigma-Aldrich, R9133) at room temperature and washing with 1X PBS-T, before starting the next round of AF imaging and staining. Subsequent rounds of staining were conducted with primary antibodies conjugated to a fluorophore, and included the immunosuppressive myeloid cell marker ARG1 (Cell Signaling Technology, 35298; AlexaFluor 555 conjugated, 1:100). All rounds of imaging and slide storage were done in a solution of PBS with 50% glycerol. Staining quality and fluorescence removal were verified after each round. The fully stitched images were imported into HALO® image analysis software (Indica Labs) for visualization.

To quantify features of the progenitor niche in human pancreatic samples, we used the Cell DIVE imaging system to stain a human pancreatitis tissue microarray (TissueArray PA691). This tissue sample spanned cases of mild, acute and chronic pancreatitis, derived from non-neoplastic and tumor adjacent tissue (see **Data S3** for sample metadata). Our panel design leveraged the possibility to conduct standard primary-seconday immunofluorescence in the first round of imaging, allowing quantification of markers of the progenitor niche TNC (R&D systems, MAB2138; RRID:AB_2203818, 1:250), MSN (Proteintech, cat#26053, RRID:AB_2880353, 1:100), and uPAR (R&D systems, AF807, RRID:AB_355618, 1:50), while leveraging the signal amplification that comes with this staining strategy. Importantly, after the first round of staining, we blocked samples with 5% rabbit serum (Sigma-Aldrich R9133), 5% rat serum (Sigma-Aldrich R9759) and 5% goat serum (Sigma-Aldrich G9023) 1X PBS for 1 h at room temperature. Similarly, we added 5% rabbit serum, 5% rat serum and 5% goat serum to the second round of staining. All stainings after the first round were conducted using fluorophore-conjugated primary antibodies.

**Data S3** specifies the staining panel used to generate **Figure 4**. **Single-molecule FISH**

#### Coverslip preparation for smFISH

Coverslips were prepared as described^114^. Briefly, 40-mm–diameter #1.5 coverslips (Bioptechs, 0420-0323-2) were cleaned in batches by arranging in a wafer boat (Entegris, A23-0215) and immersing in a 1:1 mix of 37% HCl and methanol at room temperature for 30 min. Coverslips were then washed twice with Milli-Q water, and once with 70% ethanol, followed by gentle drying with nitrogen gas. Cleaned coverslips were coated with a silane layer to allow stabilization of a polyacrylamide gel during smFISH staining, following published protocols^114^: they were submerged in 0.1% (vol/vol) triethylamine (Millipore, TX1200) and 0.2% (vol/vol) allyltrichlorosilane (Sigma, 107778) in chloroform for 30 min at room temperature, washed once with chloroform, washed once with 100% ethanol and dried using nitrogen gas. Coverslips were stored long-term in a desiccated chamber.

To prepare for staining individual samples, silanized coverslips were coated with 0.1 mg mL^-1^ Poly-D lysine (Thermo Fisher Scientific, A3890401) at room temperature for 1 h in a 6-cm tissue culture plate. They were then washed once with 1X PBS, and 3 times with nuclease-free water. Coverslips were lifted after each wash, using either tweezers or a needle, to ensure that both sides of the coverslips were exposed to the solution. Coverslips were left to dry for at least 2 h in a tissue culture hood before proceeding to tissue sectioning.

#### Tissue sectioning, fixation and permeabilization

Tissue section preparation was conducted following a published protocol^113^. Briefly, 10-μm tissue sections were cut using a cryostat and mounted onto poly-D lysine coated coverslips, then placed face-up on a 6-cm tissue culture dish for all subsequent wash and incubation steps. Coverslips were dried for 5–10 min at 50°C, and placed on dry ice until all samples were sectioned. Next, plates with coverslips were transferred to ice, treated with 3 mL 1X PBS to melt the OCT, and fixed at room temperature with 4% PFA 1X PBS for 10 min. Coverslips were then washed three times with 1X PBS, treated with ice-cold 70% ethanol and maintained at 4°C overnight for permeabilization.

#### Pre-staining treatment of permeabilized tissues

After overnight ethanol incubation, coverslips were rehydrated with 1X PBS on ice for 10 min. To bleach endogenous fluorescence of lineage reporters, tissues were exposed to a bleaching solution of 3% hydrogen peroxide (Fisher, H325-500), 1:600 37% HCl (vol/vol) 1X PBS, and placed under a heat lamp for 1 h^115^, then washed twice with 1X PBS and once with 2X SSC. Next, they were treated with pre-warmed (37°C) digestion solution containing 20 μg mL^-1^ proteinase K (Sigma, 3115836001) in 2X SSC, and incubated at 37°C for 10 min. This step enhances the permeabilization of probes in an optimized protocol for RNA staining in pancreatic tissue^113^. To remove proteinase K, coverslips were washed 3 times with 2X SSC. To prepare coverslips for hybridization, they were treated with pre-hybridization solution, composed of 30% formamide (Thermo Fisher Scientific, AM9344) in 2X SSC and incubated for at least 3 h at 37°C, as previously described^113^.

#### Staining with primary probes

Computational probe design is described below (**Probe design for multiplexed smFISH**). Primary probes were diluted at a 100 nM final concentration per probe in 3H staining buffer, composed of 30% formamide, 10% dextran sulfate (Sigma Aldrich, D8906-50G), 1 mg mL^-1^ yeast tRNA (Thermo Fisher Scientific, 15401029) in 2X SSC^114^. In addition, this staining solution had a final concentration of 2 μM anchor probe, a 15-nt sequence of alternating dT and thymidine-locked nucleic acid (dT+) with a 5′-acrydite modification (Integrated DNA Technologies), designed to anchor all polyadenylated RNAs to a polyacrylamide gel in subsequent steps. Next, hybridization chambers were prepared by attaching parafilm on the surface of a 6-cm tissue culture dish. Upon completion of pre-hybridization incubation, a 100-μL droplet of hybridization solution and probes (100 nM per probe) was placed on the center of the hybridization chamber, and coverslips were placed face down so that the hybridization solution uniformly covered the tissue, taking care of removing bubbles that may have formed in the parafilm–coverslip interface. Hybridization chambers were placed on a 15-cm dish, with a wet Kimwipe used as a humidity buffer, and incubated at 37°C for 36h–48 h.

#### Post-hybridization wash

Upon completing incubation with primary staining solution, post-hybridization wash buffer composed of 30% formamide in 2X SSC was prepared, and pre-heated to 37°C. Coverslips were then washed face-up with post-hybridization wash buffer at 47°C for 30 min. This washing step was repeated for a second 30-min incubation with fresh post-hybridization wash buffer.

Lastly, coverslips were transferred to 2X SSC solution and maintained at 4°C until the next step.

#### Gel embedding

Samples were embedded on a thin layer of polyacrylamide gel, to allow subsequent tissue-clearing through digestion of protein and lipids. To prepare the workspace for gel embedding, microscope glass slides (Premier, 6101) were washed with 70% ethanol and RNAse away (Thermo Fisher Scientific, 21-402-178), placed on a benchtop, and covered with 0.5 mL gel slick (Lonza, 50640), cleaning excess with a Kimwipe. The gel solution was composed of 4% (vol/vol) of 19:1 acrylamide/bis-acrylamide (BioRad, 1610144), 60 mM Tris⋅HCl pH 8 (Invitrogen, 15568-025), 0.3 M NaCl (Boston Bioproducts, R-244), supplemented with the polymerizing agents ammonium persulfate (Sigma, 09913) and TEMED (Sigma, T7024) at final concentrations of 0.03% (wt/vol) and 0.15% (vol/vol), respectively, as described^114^. The solution was then degassed using a vacuum chamber (Thermo Fisher Scientific, 53050609) until bubbles stopped rising to the surface of the solution. Coverslips were rinsed twice with gel solution. A 100-μL droplet of gel solution was placed on a glass slide, and coverslips were placed face-down on the slide so that the gel solution spread evenly at the slide-coverslip interface. Polymerization was completed in 2 h at room temperature, after which gel-embedded coverslips were lifted from the glass slide with the aid of a razor-blade, and transferred to a 6-cm tissue culture dish with 2X SSC.

#### Digestion

Gel-embedded samples were subjected to an overnight treatment with digestion solution, aimed at clearing proteins and lipids from the samples, improving the signal to noise for RNA detection. Digestion solution was composed of 2% SDS (Invitrogen, AM9822), 0.25% TritonX (Acros organics, 327371000), 1:100 dilution of proteinase K (New England Biolabs, P8107S) in 2X SSC. Samples were incubated overnight in digestion solution at 37°C. Following overnight digestion, samples were rinsed once with 2X SSC, transferred into a separate plate with 2X SSC, and washed for 30 min with gentle agitation. The 2X SSC solution was replaced, for a second 30-min wash.

#### Staining with secondary probes

We used readout probes consisting of a 20-bp oligonucleotide conjugated to a fluorophore (Alexa Fluor 488, Cy3B, Cy5 or Alexa Fluor 750) via a disulfide bond. Fluorescent conjugated probes were purchased from Biosynthesis Inc. The secondary staining solution was composed of 5% ethylene carbonate (Sigma Aldrich, E26258-100G) in 2X SSC, and supplemented by 3 nM of a secondary readout probe for each fluorescent color and 1 μM DAPI. Secondary staining was conducted following the same procedure for the primary staining step, with the exception that it was conducted for 20 min at room temperature, covering samples with aluminum foil.

Following hybridization, samples were washed once with a 10% ethylene carbonate 2X SSC solution for 20 min with gentle agitation, and three times with 2X SSC for 5 min per wash.

#### Iterative smFISH imaging

We prepared the following buffers for iterative smFISH imaging: (1) Wash buffer: 10% ethylene carbonate 2X SSC, 2.5 mL per staining round; (2) Cleavage buffer: 10% TCEP (Sigma-Aldrich, 646547-10X1ML) 2X SSC, 3 mL per cleavage round. TCEP in the cleavage buffer allows reduction of disulfide bond linking fluorophores to oligonucleotides in readout probes for rapid extinction of fluorescent signal; (3) Imaging buffer: 10% glucose 2X SSC, supplemented with catalase (Sigma-Aldrich, C3515; 17.5 μg mL^-1^ final concentration) and glucose oxidase (Sigma-Aldrich, G2133; 1.4 mg mL^-1^ final concentration), 2 mL per imaging round. Imaging buffer was stored under a layer of 1.5 mL mineral oil to minimize oxygen in solution during sequential rounds of staining and imaging; (4) 2X SSC, 40–50 mL per experiment. Furthermore, we prepared readout probe mixes for each round of staining. Readout probes were diluted to a final 3 nM concentration per probe, in 5% ethylene carbonate 2X SSC, supplemented with Murine RNAse inhibitor (New England Biolabs, M0314S; 1:400 dilution). Buffers and readout probe mixes were loaded into a custom-build fluidics control system^116^ that can interface with the NIS Elements image acquisition software (v 5.31.02) using custom macros.

Coverslips were mounted in a commercial flow chamber (Bioptechs, FCS2) sandwiched between a 0.75-mm-thick flow chamber gaskets (Bioptechs, 1907-100; DIE# F18524), a micro-aqueduct slide (Bioptechs, 130119-5NC) and a second 0.75-mm-thick flow chamber gaskets (Bioptechs, 1907-100; DIE# 449673-A), as described^117^. We first cut the gel so that it would fit in its entirety within the rectangular opening of the flow chamber gasket. We placed the flow chamber for imaging on a Nikon Ti2 inverted microscope using the FCS2 stage adapter (Bioptechs, 060319-2-2611), and used our fluidics system to flow in 20X SSC into the sample in order to eliminate bubbles in the tubbing and chamber. Next, we flowed imaging buffer into the sample and generated a low magnification map of the entire tissue using a 20X Plan APO objective (Nikon, MRD00205). We then switched objectives to a high magnification 60X Plan APO immersion oil objective (N.A. 1.4, W.D. 0.13 mm, F.O.V. 25 mm, Nikon, MRD01605) to resolve individual mRNAs. We used tape to minimize the movement of the plate holder during sequential rounds of imaging, which was important for preventing positional drift throughout the experiment.

Imaging cycles were conducted using the following parameters:

● Staining. Flow staining buffer for 4 min at a rate of 0.5 mL min^-1^. Incubate for 20 min.
● Wash. Flow wash buffer for 5 min at a rate of 0.4 mL min^-1^.
● Imaging. Flow imaging buffer for 3 min 40 sec at a rate of 0.5 mL min^-1^. Take 7 z-stacks per field of view, using a 1-μm step size, for a coverage of −3 μm to 3 μm around the mid-plane, using perfect focus throughout the entire experiment.
● Cleavage. Flow cleavage buffer for 4 min at a rate of 0.5 mL min^-1^. Flow cleavage buffer for 10 min at a rate of 0.1 mL min^-1^. Incubate for 10 min. Flow 2X SCC for 5 min at a rate of 0.5 mL min^-1^.

We collected images from FOVs without tissue or sources of bright autofluorescence that would allow us to estimate non-uniform illumination and detection profiles in each fluorescent channel, and correct for these in downstream image processing steps.

### Spatial transcriptomics using Xenium

FFPE blocks were sectioned and processed according to 10x Genomics user guidelines (CG000580, CG000582, CG000584). Briefly, 5-µm tissue sections were trimmed from FFPE blocks and placed within the fiducial frame of the Xenium slide (PN-1000460). The slides were air dried, baked at 42°C for 3 h and stored in a desiccator. Tissues were then deparaffinized, rehydrated and de-crosslinked using Xenium Sample Prep Reagents (PN-1000460). Tissues were hybridized overnight using a custom probe set (480 gene panel). The probes were ligated and amplified *in situ*. Tissues were quenched to remove autofluorescence and counterstained with DAPI. Slides with their corresponding decoding file were loaded and imaged on the Xenium instrument.

**Table S2** details the samples we analyzed with the Xenium platform.

## IN VITRO TREATMENTS

### Organoid culture and treatments

Premalignant pancreatic organoids were cultured in complete organoid medium unless otherwise specified. For passaging, organoids were dissociated with TrypLE Express (Fisher) and split at 1:4-1:6 ratios. For TGF-β treatments, organoids were passaged at a 1:4 ratio and cultured for two days in complete organoid medium lacking the TGF-β receptor inhibitor A83-01 (Tocris). After two days, 100 pM TGF-β (R&D Systems) was added and organoids were collected for RNA isolation 24 hours later.

### Quantitative RT-PCR

To collect total RNA from organoids, growth medium was aspirated, and organoids were washed once in 1x PBS. RLT Lysis Buffer (Qiagen) was then added directly to the wells and Matrigel was manually disrupted by pipetting until completely dissolved. RNA was then isolated using the RNeasy Mini Kit (Qiagen) per the manufacturer’s instructions. RNA concentration was measured using a NanoDrop 8000 (Thermo Scientific) and cDNA was synthesized using the AffinityScript cDNA Synthesis Kit with random primers (Agilent). cDNA was diluted with 200–400 µL sterile water, depending on the amount of input RNA (up to 1 µg). Quantitative RT-PCR was performed on a QuantStudio 6 Flex (Applied Biosystems). Each reaction consisted of 4.5 µl cDNA, 5 µl PerfeCTa SYBR Green FastMix (Quantabio), and 0.5 uL mixed forward and reverse primers (500 nM each). *Rplp0* and *Rps13* were used as housekeeping genes. Relative mRNA expression was calculated using the ΔΔCt method. Primer sequences are as follows:

*Rplp0* (ref^118^):

- Forward: 5’-GCTCCAAGCAGATGCAGCA-3’

- Reverse: 5’-CCGGATGTGAGGCAGCAG-3’

*Rps13* (Roche Universal Probe Library):

- Forward: 5’-TGCTCCCACCTAATTGGAAA-3’

- Reverse: 5’-CTTGTGCACACAACAGCATTTA-3’

*Tgfb1* (Origene):

- Forward: 5’-TGATACGCCTGAGTGGCTGTCT-3’

- Reverse: 5’-CACAAGAGCAGTGAGCGCTGAA-3’

*Hmga2* (ref^119^):

- Forward: 5’-AGACCCAGAGGAAGACCCAAAG-3’

- Reverse: 5’-TTCAGTCTCCTGAGCAGGCTTC-3’

*Lgals4* (Origene):

- Forward: 5’-CTTCAGTCCATCAACTTCCTCGG-3’

- Reverse: 5’-GGCAAGGTGTTCATCTGTGGAG-3’

## QUANTIFICATION AND STATISTICAL ANALYSES

## GENE EXPRESSION SIGNATURE DERIVATION FROM EXISTING DATA

### Premalignant state signatures

To derive gene expression signatures for major premalignant subpopulations, we computed pairwise differential gene expression between discretized premalignant states as defined by Burdziak, Alonso-Curbelo and colleagues^7^. We used the wald test in the diffxpy package (v0.7.4, https://github.com/theislab/diffxpy?tab=readme-ov-file) and library size as numeric covariates. We identified upregulated genes using the following thresholds: *qval* < 0.05, log_2_ fold-change > 1, mean expression > 0.05, and defined a signature as the set of genes upregulated in a specific subpopulation in every pairwise comparison between premalignant states.

### SMAD4-dependent TGFβ induced genes

We reanalyzed published bulk RNA-seq data from SMAD4-proficient and SMAD4-deficient PDAC organoids stimulated with TGFβ or vehicle^32^. We used the R DESeq2 package (v1.32.0)^120^ to model gene counts as a function of treatment (TGFβ stimulation of vehicle) and SMAD4 status (SMAD4-proficient or SMAD4-deficient). We identified genes that are upregulated by TGFβ stimulation in a SMAD4-proficient context. Furthermore, we required that upregulation was sensitive to SMAD4 status. We used the following thresholds to identify upregulated genes: *padj* < 0.001 and *log2FoldChange* > 1.5, resulting in a gene signature of 88 genes.

### Glycolysis (Warburg) signature

This signature is composed of a curated list of glycolysis-related enzymes (*Hk1, Hk2, Gapdh, Pgk1, Eno1, Pkm, Ldha*), including glucose, lactate and pyruvate transporters upregulated during the Warburg effect (*Slc16a1, Slc16a3, Slc2a1*)^121^, as well as the hypoxia master regulator *Hif1a*.

### Public transcriptional signatures

The following list specifies the sources and description of public signatures provided in **Data S8**.

● *p53 Fisher* (ref^102^). Curated targets of the tumor suppressive transcription factor p53
● *p53 TSAG* (ref^103^). Effectors of *p53*-dependent tumor suppression that are also bound by p53 in irradiated MEFs
● *p53 restoration* (ref^64^). Upregulated upon *p53* restoration in PDAC cells *in vitro*
● *Senescence UP* (ref^105^). Upregulated in IMR90 human fibroblasts upon *Hras*^V12^-induced senescence.
● *HALLMARK EMT* (ref^104^). Genes defining epithelial-mesenchymal transition, as in wound healing, fibrosis and metastasis.
● *Kras signaling/Fosl1* (ref^34^). Consistently upregulated genes in *Kras* mutant vs wild-type mouse and human tumors.
● *Kras injury* (ref^26^). Genes upregulated in *Kras*^G12D^+ pancreatic epithelial cells, compared to Kras^WT^ cells, both harvested 48h post-acute pancreatitis *in vivo*.
● *GOBP wound healing* (GO:0042060). The series of events that restore integrity to a damaged tissue, following an injury.
● *YAP signature* (ref^122^). Genes activated by YAP overexpression in human mammary cells (MCF10A), and YAP overexpression in mouse liver tissues or in immortalized mouse fibroblasts.
● *IFNγ response* (ref^104^). Genes up-regulated in response to IFNG (Hallmark Gene Sets).
● *p65-dependent* (ref^105^). Upregulated in IMR90 human fibroblasts upon *Hras*^V12^-induced senescence, dependent on p65 proficiency.
● *p63 ChIP-seq targets* (ref^67^). Genes bound by p63 in human breast cancer cells.

## PROCESSING AND ANALYSIS OF SINGLE-CELL DATA

### Data preprocessing and quality control

#### mRNA count matrix generation and demultiplexing

All scRNA-seq datasets were demultiplexed, barcode-corrected, aligned and UMI-corrected with SEQC^123^ using mouse genome mm10 and default parameters for samples generated using the v2 (spontaneous tumorigenesis samples) or v3 (injury-induced tumorigenesis samples) 3’ scRNA-seq kit. We summed all counts from genes that share the same gene symbol.

For samples subjected to cell hashing, we demultiplexed using an in-house method known as SHARP (https://github.com/hisplan/sharp). Hash labels were assigned to either identify a cell as belonging to a specific mouse or as a doublet or low-quality droplet. Doublet calls informed our cluster-based doublet filtering (see below). We excluded non-doublet cells without an assigned hash barcode from sample-specific analyses, but included them in condition-level analyses.

This is possible because cells from distinct conditions (experimental time point x genotype) were sequenced independently, and our cell hashing strategy aimed to distinguish biological replicates within a condition.

#### Empty droplet removal and ambient RNA subtraction

We removed empty droplets using the remove-background function of cellbender (v0.2.0)^124^, with *expected_cells* = 5000 for spontaneous tumorigenesis datasets, and 8000 for injury-induced tumorigenesis datasets (based on the number of cells targeted for encapsulation), *total-droplets-included* = 20,000, *fpr* = 0.01 (default), *learning-rate* = 0.0001 (default) and *epochs* = 150 (default). We excluded droplets with fewer than 100 mRNA counts as input into subsequent quality control (QC) analyses, and used cellbender background-corrected count matrices for downstream applications. Ambient RNA subtraction was important for mitigating the effect of CD45+ cell spike-ins during the collection of rare pre-tumor *p53*-deficient cells, as revealed by inspecting immune-related transcripts in epithelial cells (not shown). Unless otherwise stated, we used the cellbender background-corrected count matrix for downstream analyses. We aggregated counts from genes that share the same gene symbol through summation.

Preprocessed datasets published as part of our study contain raw and cellbender-corrected counts in the same AnnotationData object for ease of comparison.

#### Low-quality cell removal

For each sample, we used an iterative clustering-based approach to identify and remove low quality groups of cells. During each iteration, we applied scanpy (v1.9.1)^125^ to embed single-cell transcriptomes and identify clusters using standard library size normalization (sc.pp.normalize_per_cell), log transformation with pseudocount 1 (sc.pp.log1p), feature selection (sc.pp.highly_variable, *flavor* = ‘seurat’ and default parameters), dimensionality reduction using PCA (*n_comp* = 100), *k*-nearest neighbor (kNN) graph construction (sc.pp.neighbors, *num_neighbors* = 15) and visualization with UMAP (sc.tl.umap). Next, we used PhenoGraph to identify single-cell clusters^126^ (sc.external.tl.phenograph, *clustering_algo* = ‘leiden’) varying the parameter *k* during kNN construction (*k =* 10, 30), resulting in cluster assignments with different levels of resolution.

We removed the groups of cells with lowest summary QC metrics per cluster at each iteration, and stopped excluding when *log_lib_size* reached 7.5 and *percent_mito* fell below 20%. By varying cluster resolution, we could identify small clusters of low-quality cells that would otherwise be merged into large clusters. For some samples, we computed high-resolution clusters using *k* = 5 during the last iteration of cluster-based QC. We found that two or three iterations of this procedure per sample was sufficient to satisfy our bounding criteria, and removed all clusters with outlier QC metrics.

#### Doublet and contaminant identification

We used doubletdetection (v4.2, http://doi.org/10.5281/zenodo.2678041) with default parameters to infer doublets, using raw counts as input. For samples subjected to cell hashing, we consolidated computationally inferred doublets with doublets identified through the detection of two hash ids. Using PhenoGraph cluster assignments with different resolutions (*k* = 5, 10, 30), we identified and recorded clusters in which at least 50% of cells were inferred as doublets. At this preprocessing step, we only recorded doublets, but didn’t exclude them from the dataset. We reasoned that maintaining these annotations would be helpful for cross-sample identification of double-enriched clusters, as we have previously shown^7^.

Although our single-cell data was derived from epithelial cells sorted by fluorescent protein expression, we identified single-cell clusters corresponding to immune and stromal contaminants. We used gene expression markers of major cellular compartments to identify these clusters (*Col1a1* for fibroblasts, *Ptprc* for immune cells, *Pecam1* for endothelial cells, *Des* for pericytes), in combination with the absence of epithelial markers (*Epcam* and *Cdh1*), as well as mRNAs corresponding to fluorescent proteins used during sorting, *GFP* and *rtTA3-IRES-mKate2*. Similar to our strategy with doublet handling, we annotated but did not exclude these clusters at this stage of analysis in an effort to identify contaminants in other samples from the same batch that are too rare to form a single cluster.

#### Gene exclusion for post-cleaning preprocessing

We excluded the following classes of genes for normalization and feature selection: (i) mRNAs corresponding to fluorescent proteins and shRNAs engineered into our mouse model, (ii) mitochondrial and ribosomal transcripts, (iii) the lncRNA *Malat1*, the inclusion of which was previously shown to distort single-cell embeddings in our experimental system^7^. In excluding these genes, we aimed to minimize variation stemming from QC metrics or hard-coded experimental conditions (e.g. inducible expression of a fluorescent protein). In addition, we excluded genes expressed in fewer than 10 cells across all batches. While we excluded these genes during single-cell embedding, we kept them in the count matrix, so that the information they contained could be used in downstream analyses.

#### Within-batch data consolidation

As the final step of QC and preprocessing, we merged count tables and annotations of all samples from the same batch. We merged AnnotationData objects using the concat function of this class with *join* = ‘outer’ (include genes present in any sample) and *fill_value* = 0 (assume that a gene not present in a sample has expression of 0). We computed within-batch single-cell embedding and clustering using the strategy detailed for single sample filtering. We used cluster-level doublet annotations of individual samples to exclude doublet-enriched clusters (those in which >50% of cells were predicted to be doublets). Similarly, we removed clusters enriched in cells annotated as contaminants.

Lastly, we used outlier detection and hard thresholding to exclude a small number of cells with low QC metrics that were not identified using cluster-based exclusion. Specifically, we computed two QC metrics for each cell, log-transformed library size and log-transformed number of detected genes (excluding ribosomal, mitochondrial, fluorescent protein and shRNA mRNA counts). Although correlated, these two metrics provide orthogonal information about transcriptional complexity, as a cell may pass a library size-based threshold even when few genes dominate its mRNA counts. To identify outliers whose QC metrics deviate from those of their nearest neighbors, we used the LocalOutlierFactor function from the sklearn.neighbors package (v1.0.2) with *n_neighbors* = 100 and *contamination* = 0.1 (number of assumed outliers). In the final step, we excluded low-quality cells based on log-transformed library size < 7.5, fraction mitochondrial counts > 0.2, or number of detected genes < 300.

#### Preliminary tumor and premalignant annotations

After integrating samples within batches, we identified clusters that exhibited transcriptional and genomic patterns consistent with being derived from PDAC (see <u>Cell state annotation</u> and <u>Copy</u> <u>number inference</u> for details). Conducting these preliminary annotations within individual batches was important for determining which cells to exclude from subsequent batch correction vector calculations.

### Analysis of KP^LOH^ spontaneous tumorigenesis data

#### KP^LOH^ dataset details

Our spontaneous tumorigenesis data from KP^LOH^ mice charts PDAC progression through the benign-to-malignant transition, and is composed of three conditions: (1) *p53*-proficient cells from mice without a macroscopic tumor (pre-tumor *p53*-proficient), (2) *p53*-deficient cells from mice without a macroscopic tumor (pre-tumor *p53*-deficient), (3) *p53*-deficient cells from mice with a macroscopic tumor (tumor *p53*-deficient) and (4) *p53*-proficient cells from mice with a macroscopic tumor. Tumor *p53*-deficient cells are derived from (i) tumor-bearing KP^LOH^ mice, isolated as GFP-/mKate2+ cells (see <u>Mouse model genetics</u> for details) and (ii) samples from tumor-bearing mice from Burdziak, Alonso-Curbelo and colleagues ^7^. For batch 8, we pooled multiple mice without hashing to minimize the time between harvesting and sorting during single-cell isolation. While we lack cell-to-mouse assignments in this sample, we note that pre-tumor cells generally do not cluster by biological replicate (**Figure S1A**). **Table S1** summarizes the number of cells per sample in our spontaneous tumorigenesis dataset.

Our prior data set^7^ did not include a reporter of *p53* genetic status; thus, a subset of cells from PDAC samples co-embedded with premalignant cells during integration. We also observed that a small fraction of cells sorted as *p53*-deficient from the KP^LOH^ model co-embedded with pre-malignant cells and expressed GFP mRNA, implying that they were indeed *p53*-proficient. We filtered out such contaminants from our dataset before embedding all samples. Note that most premalignant contamination comes from primary tumors rather than metastases, supporting the notion that these were non-cancer cells embedded within the tumor. The following list summarizes the number of cells in pre-malignant clusters within primary and metastatic PDAC single-cell samples:

Primary tumor samples:

● DAC_D020_p5_Epi (Tumor p53 deficient 11): 645 cells in pre-malignant clusters
● Ag-PDAC-PT-Kate (Tumor p53 deficient 10): 180 cells in pre-malignant clusters
● 53_LHRH_PDACreg_KATE (Tumor p53 deficient 7): 14 cells in pre-malignant clusters
● DACC963PT_Kate_plus (Tumor p53 deficient 9): 7 cells in pre-malignant clusters
● PDAC-SP3 (Tumor p53 deficient 6): 5 cells in pre-malignant clusters
● 9268_PHLH_PDAC_SP (Tumor p53 deficient 8): 2 cells in pre-malignant clusters

Metastatic samples:

● Ag-Lung-Mets-Kate (Tumor p53 deficient 10): 2 cells in pre-malignant clusters
● DACC963LIVERmet (Tumor p53 deficient 9): 0 cells in pre-malignant clusters
● DACC963_mKate_plus (Tumor p53 deficient 9): 0 cells in pre-malignant clusters

#### Feature selection and normalization

We computed two embeddings independently to analyze heterogeneity in single-cell gene expression as a function of *p53* genetic status and malignant progression. The first embedding (**Figures 1** and **S1A,B**) excludes *p53*-proficient cells derived from PDAC-bearing mice (embedding 1) and the second includes all samples (**Figure S1C,D**) (embedding 2). Of note, *p53*-proficient cells from tumor-bearing mice co-embedded with *p53*-proficient cells from pre-tumor stage mice (**Figure S1C**), although there were more progenitor-like cells in tumor-bearing animals (**Figure S1F**).

To identify a set of highly variable genes (HVGs) that capture the variability across samples in all 3 batches, we conducted within-batch feature selection using scanpy’s sc.pp.highly_variable_genes function with *n_top_genes* = 3000 and *flavor* = ‘seurat_v3’. The final set of HVGs comprised the union of genes selected in each batch, resulting in 5159 genes (embedding 1) or 5156 genes (embedding 2) capturing variation along PDAC progression.

We conducted a batch-aware normalization approach similar to Haghverdi and colleagues^127^. We estimated per-cell size factors as the total counts after excluding mitochondrial, ribosomal, transgenic and *Malat1* mRNAs (see <u>Gene exclusion for post-cleaning preprocessing</u>), then calculated the median size factor per batch, and rescaled size factors to equalize medians across batches. We normalized data by dividing counts by rescaled size factors, and multiplying by the median rescaled size factor across all cells. Lastly, we applied a log transformation to the count matrix with pseudocount = 1.

#### Dimensionality reduction, batch correction and KNN construction

We computed a batch-corrected latent space for subsequent processing steps using mutual nearest neighbors in the batchelor R package (v1.8.1)^127^ with batches 1 and 2 as reference. These batches contained the majority of cells in the dataset, and spanned all timepoints and genotypes. We used the fastMNN function with *cos.norm* = FALSE, *d* = 100 (number of components to keep), *correct.all* = TRUE and *prop.k* = 0.1 (default), resulting in a corrected latent space of 100 components that capture 45% of reference batch variance for both embedding 1 and embedding 2. Although the inflection point in the cumulative explained-variance curve was at 62 PCs (explaining 42% of variance), we chose to keep more PCs because this dataset was composed of both cancer and premalignant states, and our subpopulations of interest (e.g. progenitor-like cells) were rare (1–2% of premalignant cells).

We computed this latent space on log-normalized counts, using only HVGs. In addition, when calculating correction vectors, we excluded cells corresponding to PDAC clusters; we previously showed that each tumor forms a distinct cluster in this model and reasoned that including PDAC cells could remove and distort true biological heterogeneity in the dataset. We note, however, that these clusters were subjected to batch correction using correction vectors estimated from non-PDAC cells. The count matrix remains unmodified in this approach. Lastly, we constructed a kNN using the batch-corrected latent space as input to scanpy’s sc.pp.neighbors function, using *num_neighbors* = 30.

#### Single-cell data visualization

To visualize our single-cell data, we first applied Uniform Manifold Approximation and Projection (UMAP) for dimensionality reduction using scanpy’s sc.tl.umap function and the precomputed kNN graph as input (*k* = 30). We also computed a force-directed layout (FDL), which captures continuities in data and highlights transitional subpopulations in dynamic systems, using the forceatlas2 python package^128^. We used the diffusion operator of our single-cell data as input to compute force-directed layouts (**see Construction of the diffusion operator and diffusion maps**). This strategy incorporates local changes in distances along the single-cell manifold into graph visualization.

#### Copy number inference

We inferred karyotypes from single-cell transcriptomes to better distinguish PDAC from premalignant status, and to characterize genomic diversification following spontaneous *p53* loss. To infer chromosome-level changes, we used a custom implementation of inferCNV (inferCNV of the Trinity CTAT Project, https://github.com/broadinstitute/inferCNV), as outlined below. inferCNV assumes that changes in copy number cause corresponding changes in gene expression, which are detectable in local genomic neighborhoods despite being subject to variation by cell state and by other factors.

We first selected a *p53*-proficient sample^28^ as a near-diploid reference, and computed the mean expression of each gene from library size–normalized counts (without log transformation). We excluded genes with low expression (*min_threshold* = 0.1), reasoning that they are less likely to reveal robust gene expression differences caused by genomic changes. Next, we ordered genes by genomic coordinates (UCSC mm10). For each gene x cell pair in the KP^LOH^ dataset, we computed the log_2_ fold-change in gene expression (*pseudocount* = 0.1 for both numerator and denominator) over the mean expression of that gene in the reference cell set, clipping log_2_ fold-change estimates to [-3, 3] to limit the effect of outliers. We computed the sliding average log_2_ fold-change over a window of consecutive genes in the same chromosome (*window_size* = 100), trimming chromosome ends. Lastly, we recentered average log_2_ fold-change expression profiles by subtracting the values of each cell by their median, resulting in our final proxy for copy number changes. To cluster inferred karyotype profiles, we computed a simplified matrix, in which each cell is described by the average log_2_ fold-change expression of each chromosome. We clustered this simplified matrix using hierarchical clustering (*method* = ‘ward’, *metric* = ‘euclidean’) implemented in the cluster.hierarchy module of scipy (v1.7.3).

Our approach incorporated two modifications to the standard approach. First, our initial examination of inferred copy number profiles revealed that small groups of biologically related genes could distort estimates. For example, we identified a cluster of carboxypeptidases (*Cpa1, Cpa2, Cpa5, Cpa4*) on mouse chromosome 6 that are expressed at high levels in acinar and ADM cells, causing spikes in inferred copy number that could be mistakenly interpreted as focal amplification. We therefore removed all such gene groups from copy number inference through manual inspection of spikes in gene smoothed gene expression profiles with smaller window sizes (5–20 genes), as well as ribosomal and mitochondrial genes (see **Data S8** for excluded genes). Our approach prioritizes robust estimation of chromosome-level copy number changes over the identification of more focal alterations and highlights an important opportunity for feature selection in developing copy number inference strategies.

Second, we incorporated iterative copy number inference to mitigate batch effects during karyotype estimation. We could use within-sample diploid references for pre-tumor stage inference, because we compared *p53*-proficient samples (expected to be near diploid) with *p53*-deficient samples. Notably, the only recurrent change in pre-tumor *p53*-proficient cells was the gain of chr6, as reported in the KP^LOH^ model^28^. The distribution of the average smoothed log_2_ fold-changes in chr6 was bimodal, providing a natural threshold to identify cells that gained this chromosome (average log_2_ fold-change > 0.17). We used cells without this event as diploid references within each mouse, reducing variability in inferred copy number profiles.

To classify pre-tumor *p53*-deficient cells as genomically ‘quiet’ or ‘rearranged’, we computed a simplified karyotype matrix in which each entry is the average copy number change of each chromosome in each cell. Next, we binarized this matrix by identifying entries > 0.16 (indicating gains) or < −0.16 (indicating losses). We selected these thresholds based on the distribution of average copy number changes across all cells and chromosomes in the dataset. We defined genomically ‘quiet’ cells as those with fewer than 9 gain or loss events, and ‘rearranged’ cells as those with more than 9 events. These thresholds captured differences in transcriptional states that distinguished premalignant-like and cancer-like clusters (**Figures 1** and **S1**) in the presence of noisy karyotype inference from transcriptomes.

#### Refinement of condition assignment

We used copy number profiles and transcriptome-based PhenoGraph clusters to refine assignment of individual cells to cancer-like or premalignant. First, we identified a rare group of pre-tumor cells sorted as GFP+, but that lacked GFP mRNA expression and chr11 loss (containing the *p53* locus). Given that these are criteria for detecting *p53* deficiency in this mouse model, we re-assigned them as pre-tumor *p53*-deficient cells (*n* = 4 cells reassigned).

#### Annotation of premalignant states

To annotate cell states in the premalignant pancreas, we first used the scanpy implementation of PhenoGraph sc.external.tl.phenograph (*k* = 30, *clustering_algo* = ‘leiden’) with gene expression signatures from our published dataset^7^ (see **Premalignant state signatures**). First, we standardized our count matrix by computing the z-score expression of each gene across all cells. To calculate a signature score per cell, we averaged the z-scored expression of signature genes in each cell. We aggregated signature scores at the cluster level by averaging, and standardized such average scores across clusters. Lastly, we used the searborn (v0.11.2) clustermap function to guide manual cluster annotation.

We noted that while some cell states were clearly separated from the bulk of the premalignant epithelium (e.g. ADM, tuft and neuroendocrine cells), the majority of epithelial cells varied along a phenotypic continuum linking gastric-like and progenitor-like states. To capture continuity between these states, we used diffusion component analysis^42,129^, as diffusion components represent axes of variation in the data and can describe successive cell-state transitions along the phenotypic manifold (see **Construction of the diffusion operator and diffusion maps**). In our spontaneous tumorigenesis dataset, the second diffusion component (DC2) correlated with single-cell progenitor-like scores (**Data S1.1E**). To identify a boundary between gastric-like states and progenitor-like states along this phenotypic continuum, we used the triangle method to identify a threshold in the distribution of cell densities along DC2 (**Data S1.1F**). Cell-state discretization in light of this continuum was helpful to interpret cell-state-specific consequences of spontaneous *p53* loss.

Genes upregulated in malignant cells in the KP^LOH^ model We leveraged our *p53*-proficient and *p53*-deficient single-cell data in the KP^LOH^ background to derive a signature of genes upregulated in PDAC compared to premalignant cells. By adopting a pseudobulking approach for differential gene expression, we could quantify changes in average transcript expression between premalignant and malignant cells in a manner agnostic to the subpopulation structure of each sample, while leveraging inter-replicate variability for the derivation of a robust expression signature. We restricted our analysis to samples from batches 1 and 2 (see **Table S1**), which were collected simultaneously and sequenced using the same reagents to avoid variation from technical sources. Furthermore, we excluded metastasis samples to focus on pancreas-derived cells. In total, we analyzed 4 pre-tumor *p53*-proficient samples and 6 tumor-derived *p53*-deficient samples. To identify differential expression, we aggregated unnormalized, cellbender-corrected counts per sample to construct a pseudobulk count matrix with genes as rows and samples as columns. Next we used the R package DESeq2 (v1.42.1) to test for differential gene expression between PDAC and premalignant conditions, using *design* = “∼ condition” to model counts. We identified upregulated genes as those with padj < 0.001 and log_2_(fold change) > 1.5, resulting in a signature of 941 genes.

#### Simultaneous visualization of multiple signatures

To visualize multiple signatures in the same single-cell layout (**Figure 1E**) we used a previously implemented signature-based pseudo-coloring strategy^7^. We normalized scores for each signature by subtracting the minimum signature value and dividing by the 99^th^ quantile of such scores. We define the signature matrix *S*_*n*×*m*_ such that *S*_*i*,*j*_ is the normalized score for signature_j_ in cell_i_, and we define a color-encoding matrix *W*_*m*×_*_3_* where *W*_*j*,·_is the RGB vector representation of the color associated with signature_j_. The pseudo-coloring of a cell is defined by the matrix multiplication *S* × *W*. Because RGB components are bounded between [0,1], we clip values to 1 after matrix multiplication. This approach is most effective in simultaneously visualizing multiple phenotypically distinct subpopulations and a limited number of mixed subpopulations, as pseudo-colors can saturate due to the effect of summation and clipping.

#### Construction of the diffusion operator and diffusion maps

Throughout this work, we make use of diffusion maps as a latent representation of single-cell expression data that is analogous to a non-linear version of principal components. A diffusion map captures dominant directions of diffusion along a single-cell manifold, understood as a random walk in which cells are allowed to reversibly transition between similar transcriptional states. We follow the computation of diffusion maps as previously described^42,43,129^.

Let *M*^*n*×*m*^ be a matrix representing the log-normalized expression of *m* genes in *n* cells. We define the k-nearest neighbor graph (kNN) *G*^*n*×*n*^ as a sparse matrix such that *G*_*i*,*j*_ is the Euclidean distance in principal component space between cell_i_ and cell_j_ when cell_j_ belongs to the k-nearest neighbors of cell_i_, and *G*_*i*,*j*_ = *0* for every other cell_j_ in the data.

We compute the cell–cell affinity matrix *A*^*n*×*n*^ by applying an adaptive Gaussian kernel to the kNN matrix:

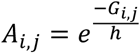

where *h* varies for each cell_i_ and is defined as the distance to its *k′* closest cell (we use *k^′^* = *10* for a kNN graph with *k* = *30*). The width *h* of the Gaussian kernel determines the rate at which *A*_*i*,*j*_ decays as a function of distance. Thus, an adaptive width allows controlling for heterogeneity in cell density along the single cell manifold.

To compute the diffusion operator *T*^*n*×*n*^, we symmetrize the cell-cell affinity matrix, set the diagonal to 0, and normalize by row, resulting in a row stochastic matrix. *T*_*i*,*j*_ can be interpreted as the transition probability from cell_i_ to cell_j_. Further exponentiation of the diffusion operator results is diffusion, a random walk over the kNN graph that results in long-range connectivities between single cells based on short-scale phenotypic transitions.

Although Euclidean distances on PC space captures cell–cell similarities at the local level, and is routinely used to construct a kNN graph during manifold estimation, they fail to capture distances over long ranges due to non-linearities in the phenotypic manifold. Diffusion distance—intuitively understood as the result of a diffusion process or a random walk over the kNN graph—captures long-range cell-cell connectivities while respecting such non-linearities. A diffusion map results from the eigen-decomposition of the diffusion operator. The eigenvectors of such decomposition, termed diffusion components, provide a new representation of the data that can be used to approximate diffusion distance. Ordering eigenvectors by their corresponding eigenvalue, and keeping the top *L* eigenvectors allows estimation of the diffusion distance between two cells:

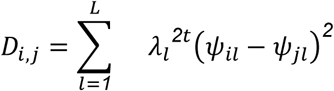

Where *λ*_*l*_ is the top *l*^*th*^ eigenvalue of the diffusion operator, *ψ* it’s associated eigenvector, and *t* is the number of diffusion steps (*t* = *3* in our analysis). Because the 0^th^ eigenvector of the diffusion operator is a constant vector, we exclude this from the diffusion map, following work from Haghverdi and colleagues^43^.

#### Diffusion distance analysis

To quantify transcriptional similarity between premalignant cells and PDAC, we used diffusion distance, a quantity that allows estimation of long-range cell–cell connectivities while respecting non-linearities in the phenotypic manifold (see **Construction of the diffusion operator and diffusion maps**). To compute diffusion distance between premalignant and malignant cells (**Figure 1F**), we used the eigenvectors associated with the 17 highest eigenvalues of the diffusion operator, based on the second eigengap as the threshold criterion. To calculate the similarity between pre-tumor *p53*-proficient (premalignant) and cancer cells, we computed the diffusion distance from every premalignant cell to the closest cancer cell (annotated as *p53*-deficient tumor or microtumor).

#### Differential gene expression

To identify gene expression programs selectively upregulated in progenitor-like cells, we computed differential gene expression between pretumor *p53*-proficient progenitor-like cells and other pretumor *p53*-proficient epithelial cells. We used the wald test in the diffxpy package (v0.7.4, https://github.com/theislab/diffxpy?tab=readme-ov-file), with raw counts as input, batch as a categorical covariate, and library size as a numeric covariate. We filtered out genes with mean expression < 0.05 before testing for significance of enriched gene sets using the GSEA implementation of gseapy (v1.1.2), using the MSigDB Hallmark 2020 database, log_2_ fold-change estimates as the rank variable, and FDR < 0.1 as a significant threshold.

To assess cell state-dependent consequences of *p53* loss, we computed differential gene expression between pretumor *p53*-proficient cells and pre-tumor *p53*-deficient cells with ‘quiet’ genomes (see <u>Copy number inference</u> for details). We used the wald test in the diffxpy package, with raw counts as input and library size as a numeric covariate. We identified downregulated genes upon *p53* loss as those with qval < 0.05, log_2_(fold change) < −1 and mean expression > 0.05. To visualize gene expression as a function of cell state and *p53* status (**Figure 1G**) we *z*-scored log-normalized counts using the mean and standard deviation of pre-tumor *p53*-proficient cells as reference. This standardization strategy aims to highlight deviations in gene expression attributed to *p53* deficiency and doesn’t depend on inclusion or exclusion of tumor-stage samples.

**Data S9** contains differential gene expression results. Quantification of p16^INK4A^ and p19^ARF^ isoforms The *Ckdn2a* locus encodes two structurally and functionally distinct gene products, p16^INK4A^ and p19^ARF^, both of which mediate tumor suppression through different mechanisms. cDNAs for these gene products result from alternative use of the exon 1α (p16^INK4A^) or exon 1β (p19^ARF^), spliced into the shared exon 2. Most reads captured through 3’-end sequencing are unable to distinguish between these two gene products, but we found rare reads that spanned isoform specific splice junctions, allowing unambiguous determination of *Cdkn2a* isoforms. To identify such reads, we used our custom scRNA-seq processing pipeline SEQC to scan aligned bam files from pre-tumor *p53*-proficient samples for reads that (1) fell within the boundaries of exon 1β and exon 2 of the *Cdkn2a* locus (chr4:89276895-89276975), and (2) showed evidence of splicing as evidenced by a gap flag in the CIGAR string. Next, we aligned reads to the spliced p16^INK4A^ or p19^ARF^ sequences, to assess the isoform associated with each read.

### Processing and analysis of *p53* knockdown data

#### Dataset details

Our injury shp53 cohort contained dissociated single-cell data from *Kras*^G12D^+ epithelial cells collected from KC^shp53^ or KC^shRen^ mice 3 weeks after injury to induce pancreatitis. This dataset, composed of two batches with 2–3 mice per genotype per batch, formed the basis of our investigations on the cell-intrinsic consequences of *p53* loss in the context of pancreatic injury. **Table S1** summarizes the number of cells per condition in our *p53* perturbation dataset.

#### Cell filtering

Preliminary embeddings of filtered and merged objects during data cleaning (see **Data preprocessing and quality control**) revealed a PhenoGraph-defined (*k* = 30) subpopulation of 210 premalignant cells that coexpressed divergent cell-type markers—*Cpa1* for acinar, *Msn* for progenitor-like, *Muc6* for gastric-chief-like and *Anxa10* for gastric-pit-like cells. This subpopulation was present in only one batch, and 95% of its cells were derived from a single biological replicate. We excluded these cells from further analysis because the cluster was not reproducible between biological replicates.

#### Normalization, feature selection and dimensionality reduction

As we did not detect strong batch effects during exploratory analysis, we merged the two batches into the same AnnotationData object and computed embeddings as follows: (1) size factor estimation from library sizes, excluding *Malat1* and ribosomal, mitochondrial and transgenic mRNAs; (3) normalization through division by size factors, scaling by the median size factor estimate, followed by log transformation with *pseudocount* = 1; (4) selection of top 3000 HVGs in each batch using the scanpy sc.pp.highly_variable_genes function with *flavor* = ‘seurat_v3’; (5) dimensionality reduction using the scanpy PCA function sc.pp.pca., keeping 68 PCs (explaining 54% of total variance) based on the inflection point in the cumulative explained-variance curve; (6) kNN construction with *k* = 30; and (7) visualization using UMAP and FDL (see <u>Single-cell data visualization</u>). Our embedding recapitulated the structure that we previously identified in our published injury-induced tumorigenesis dataset^7^.

#### Visualization of cell state density in two-dimensional representation

To gain an intuition of how individual cell distributions change along the phenotypic manifold upon *p53* knockdown, we computed two-dimensional densities in the coordinates of our layouts. For each condition (*p53* knockdown or control), we computed a histogram summarizing cellular frequencies at different coordinates of the 2D projection of the UMAP embedding (100 bins for x and y axes). We smoothed the histogram using a 2D Gaussian kernel with *bandwidth* = 1 bin, and visualized the estimated distributions in a contour plot (**Figure 7C**). We emphasize that this procedure does not accurately estimate cellular densities in high dimensional space; however, we found it useful for communicating results of high-dimensional computations (e.g. the accumulation of progenitor-like cells with mesenchymal properties upon *p53* knockdown).

#### Differential gene expression

To assess the consequences of *p53* knockdown in premalignant epithelial cells during injury, we adopted a pseudobulking approach. PhenoGraph (*k* = 30) grouped progenitor-like cells into three clusters—one from shRen samples (shRen progenitor 1) and two from shp53 samples (shp53 progenitor 1 and 2) (**Figure S6D–F**). Diffusion component analysis suggested that different progenitor-like subpopulations lie along a phenotypic continuum and that *p53* knockdown facilitates persistence or progression of the more advanced progenitor 2 state.

We reasoned that comparing cluster pairs would allow us to distinguish direct effects of *p53* knockdown that are due to target gene activation from secondary effects that are due to changes in cell state. We therefore asked which gene expression programs change upon *p53* knockdown (1) for all progenitor-like cells, (2) for regions of the progenitor continuum with similar shp53 and shRen cell densities, and (3) that specifically characterize progenitor 2 cells.

We grouped cells by genotype (shp53 or shRen) and progenitor-class (progenitor 1, progenitor 2 or all progenitor-like cells) combination, then summed unnormalized counts to generate a pseudobulk sample. Our approach is conceptually similar to marker-based cell sorting followed by bulk RNA sequencing, and illustrates how differential gene expression results change depending on the resolution at which single-cell communities are computationally or experimentally isolated. To compute differential expression, we used DESeq2 (v1.42.1) with pseudobulk counts as input, and genotype or cell state as contrasts. We used the GSEA implementation of gseapy (v1.1.2) to query gene sets differentially expressed between different progenitor-like clusters in shp53 cells, using the MSigDB Hallmark 2020 database, log_2_ fold-change estimates as the rank variable, and FDR < 0.1 as a significance threshold. **Data S9** contains differential gene expression results.

#### Condition-aware imputation and gene signature scoring

We used MAGIC-imputed counts^129^ to compute gene expression signatures. MAGIC uses the diffusion operator (see **Construction of the diffusion operator and diffusion maps**) to share gene expression information in local neighborhoods in cell state space, mitigating the effect of dropouts in sparse single-cell datasets. Formally, the imputed count matrix is obtained by exponentiation of the diffusion operator, and multiplication with the count matrix. The exponent of the diffusion operator *t* corresponds to the number of diffusion steps (*t* = *3* in our specific implementation). Increasing *t* increases the distance over which gene expression information of any given cell influences imputed counts of another cell in the phenotypic manifold.

To avoid sharing gene expression information between cells from different genotypes, we carried out imputation separately for shp53 and shRen samples, using the same PC space to construct the kNN graph (*k* = 30) as input to the imputation process. To score gene signatures, we first computed a *z*-scored imputed gene expression matrix. We used the mean and standard deviation of imputed counts in shRen samples for *z*-scoring. This strategy aims to highlight deviations in shp53 cells relative to control cells.

#### Diffusion component analysis

Diffusion component analysis captures continuous, non-linear variation in data; we applied it to identify dominant axes of variation in our premalignant pancreas data. We constructed a diffusion operator using a kNN graph (*k* = 30) built on cells from both shp53 and shRen samples. The third eigenvector of the diffusion operator (diffusion component 3 or DC3) captured variability between the two most abundant subpopulations in the premalignant pancreas, consisting of gastric-like and progenitor-like states. To plot gene signatures as a function of diffusion component, we discretized DC3 into 100 equally sized bins and computed the average signature score for each bin, separately for shp53 and shRen genotypes (**Figure 7D**). We excluded bins with fewer than 10 cells from visualization.

### Analysis of Kras inhibitor data

We aimed to systematically quantify changes in the premalignant epithelium from acute oncogenic Kras inhibition by MRTX1133 treatment in the context of pancreatic injury. This unbiased characterization complements our targeted analyses of the effects of this treatment on the abundance of progenitor-like cells (**Figure 6B**). The three or four biological replicates from each condition (vehicle or MRTX1133-treated) were pooled, encapsulated and sequenced together, followed by sample deconvolution using cell hashing.

#### Single-cell embeddings

Starting from QC-filtered count matrices (see **Data preprocessing and quality control**), we merged data from two conditions into a single object, and generated single-cell embeddings by: (1) size factor estimation from library sizes, excluding *Malat1* and mitochondrial, ribosomal and transgenic mRNAs; (2) standard library size normalization, followed by scaling by median library size; (3) log transformation using pseudocount = 1; (4) selection of top 3000 HVGs using scanpy’s sc.pp.highly_variable function on cellbender counts with *flavor* = ‘seurat_v3’ (excluding mitochondrial, ribosomal, transgene and *Malat1* mRNA); (5) dimensionality reduction using the scanpy PCA function sc.pp.pca., keeping 57 PCs (explaining 51% of total variance) based on the inflection point in the cumulative explained-variance curve; (6) kNN construction using *k* = 30; (7) UMAP visualization using sc.tl.umap and default parameters; (8) computation of diffusion operator on the kNN graph (Gaussian kernel width determined adaptively based on distance to each cell’s 10th nearest neighbor) and (9) FDL visualization using the diffusion operator as input and initialization using UMAP coordinates. We visualized conditions and cell states as 2D density maps projected on our FDLs (**Figure S5B**). **Table S1** summarizes the number of cells contained in our single-cell object.

#### Cell type annotation

We used PhenoGraph clustering and marker gene expression to annotate premalignant subpopulations. Most subpopulations relied on highly specific established markers (e.g., *Cpa1* for ADM cells, or *Pou2f3* for tuft cells, or *Syp* for neuroendocrine cells). For gastric-like states, we used a refined set of markers, since oncogenic KRAS inhibition led to shifts in the spectrum of these states. To refine gastric-like states, we also used a smaller than typical *k* (*k* = 10) for kNN construction when inferring PhenoGraph clusters. We used the following markers to annotate premalignant states:

● Progenitor-like (*Hmga2, Msn, Itga3, Nes*)
● ADM (*Cpa1, Rbpjl, Nr5a2*)
● Tuft (*Pou2f3, Alox5, Ptgs1*)
● Neuroendocrine (*Syp, Scg5, Chga, Chgb*)
● Cycling (*Mki67, Bub1, Cdk1*)
● Duct (*Rgs5, Cp, Prox1*)
● Gastric-general (*Dmbt1*)
● Gastric pit-like (*Anxa10, Tff1*)
● Gastric chief-like (*F5, Muc6*)

Differential abundance analysis.

To test for differential abundance of distinct premalignant subpopulations as a result of MRTX1133 treatment, we used the Milo algorithm^63^. This method first identifies communities of cells on a kNN graph that partially overlap between conditions (Milo transcriptional neighborhoods), then models the cell counts from different experimental conditions in each neighborhood using a generalized linear model with negative binomial residuals. This allows testing for differences in the abundance of cells from different conditions within granular cellular states in the data. We used the miloR implementation with a precomputed kNN graph (*k* = 30) and PCA (*n_pcs* = 57), using the makeNhoods function for Milo neighborhood construction with *prop* = 0.01 and *refined* = TRUE. This approach uncovered a set of granular cell states that are either enriched or depleted (SpatialFDR < 0.1) in the premalignant pancreas upon acute oncogenic KRAS inhibition. To visualize these results, we annotated transcriptional neighborhoods by their most common cell state label, and plotted their estimated log-fold change in MRTX1133-treated vs vehicle-treated mice as a function of cell state (**Figure S5C**). Furthermore, it showed that the progenitor-like state is most dependent on persistent *Kras* signaling among premalignant subpopulations.

#### Differential gene expression analysis

We reasoned that dissecting gene expression changes *within* premalignant states as a consequence of acute oncogenic KRAS inhibition could reveal programs dependent on persistent KRAS signaling and cell-state transitions mediated by loss of oncogenic signaling. We focused on ADM cells, as this subpopulation was not depleted, but showed a marked shift in transcriptional state. We used the wald test in the diffxpy package (v0.7.4, https://github.com/theislab/diffxpy?tab=readme-ov-file) to compute differential gene expression between MRTX1133 and vehicle-treated ADM cells, using library size as a numeric covariate, and filtering to only include genes with mean expression > 0.05 that are expressed in at least 20 cells. These filters aimed to exclude low-expressed genes with potentially large fold changes that do not represent the strongest biological differences between states. To identify molecular programs upregulated or downregulated upon MRTX1133 treatment, we used the GSEA implementation of gseapy (v1.1.2), using log_2_ fold-change estimates as the rank variable, and FDR < 0.1 as a significant threshold. We independently queried multiple databases, including the MSigDB_Hallmark_2020 database for general shifts in cellular signaling and the TF_Perturbations_Followed_by_Expression database to reveal molecular regulators that may mediate cell-state shifts following oncogenic Kras inhibition. Our focus on these gene sets was motivated by the fact that chronic genetic or pharmacological inhibition of oncogenic Kras signaling has been shown to restore a normal pancreatic histology in the premalignant pancreas^130,131^.

### Processing and analysis of premalignant tumor microenvironment data

We used Flex Gene Expression (10x Genomics) to gain insights into transcriptome-wide heterogeneity in gene expression across cellular compartments in the premalignant pancreas. These data allowed us to contextualize compositional and molecular properties of cellular states associated with the progenitor niche, as identified with Xenium-based measurements, including information regarding the expression of mRNAs that were not measured in our Xenium panel, but that have important roles in myeloid and stromal compartments in the context of tissue injury and cancer.

Our dissociated single-cell data was composed of samples from KC^shp53^ mice (*n* = 2) or KC^control^ mice (*n* = 2, KC^shRen^ or KC^shp53^ without doxycycline induction of shp53). All samples came from tissue harvested 3 weeks post-pancreatitis, followed by nucleus isolation from FFPE blocks (see **Dissociated nuclei from FFPE samples**). Data from these samples was subjected to (1) QC and embedding of single-cells from all samples, and (2) computation of compartment-specific embeddings.

#### Cell and gene filtering

We used cellbender (v0.3.2)^124^ for ambient RNA correction and prediction of empty droplets. Examination of library sizes of cellbender-filtered cells revealed a unimodal distribution of ln(library size) with mean = 7.04 and standard deviation = 1.17. We conservatively filtered out cells with ln(library size) < 5, corresponding to the bottom 3.6% of the dataset. We excluded mitochondrial and transgene mRNAs (GFP and mKate2) from influencing normalization and feature selection. Furthermore, we used doubletdetection v4.2, http://doi.org/10.5281/zenodo.2678041 for prediction of doublets, removing any PhenoGraph cluster (*k* = 30 or 10) in which at least 50% of cells within the cluster were predicted to be doublets. **Table S1** summarizes the number of cells included in this dataset for each biological replicate.

#### Embedding of single cells from all samples

We generated single-cell embeddings as for other datasets, by (1) size factor estimation from library sizes, excluding *Malat1* and mitochondrial, ribosomal and transgenic mRNAs; (2) standard library size normalization, followed by scaling by median library size; (3) log transformation of count matrix with pseudocount = 1; (4) selection of top 3000 HVGs using scanpy’s function sc.pp.highly_variable with *flavor* = ‘seurat_v3’ (excluding mitochondrial, ribosomal, transgene and *Malat1* mRNA); (5) dimensionality reduction using the scanpy PCA function sc.pp.pca., keeping 100 PCs (explaining 53% of total variance). We used many more PCs than the knee point of the cumulative explained-variance curve because we sought to capture both intercompartment and intracompartment heterogeneity in this dataset, containing the full diversity of premalignant cell types; (6) kNN construction using *k* = 30; (7) Computation of PhenoGraph clusters with different resolutions (*k* = 30 or 10), using Leiden for clustering of the Jaccard similarity matrix; (7) visualization using UMAP. This strategy resulted in a consolidated dataset for the extraction of select cellular compartments for further analysis.

#### Compartment and condition-specific embeddings

Our main goal in collecting and analyzing this dataset was to provide a transcriptome-wide contextualization of the gene expression changes that we identified in Xenium-based spatial transcriptomics data. Thus, it was important to compute compartment-specific and condition-specific embeddings. We focused specifically on fibroblast and myeloid compartments, which represent the two most abundant microenvironmental cells in the premalignant pancreatic parenchyma, and those in which we identified the strongest changes in gene expression as a function of which niche they were encountered in. To compute these embeddings, we (1) isolated subpopulations based on condition filtering and identification of PhenoGraph clusters labeled by marker genes; we followed the strategy outlined above (<u>Embedding of single cells</u> <u>from all samples</u>), except that we used an adaptive strategy for determining the number of PCs to retain at each step, based on the inflection point in the cumulative explained-variance curve. **Data S10** summarizes the intermediate processing steps that we took to compute compartment and condition-specific embeddings.

### Analysis of human pancreatic epithelial data

#### Computation of *p53* activity and gene expression signatures

To investigate *p53* activity in human data, we used the scRNA-seq dataset published in Carpenter, Elhossiny, Kadiyala et al.^2^. We split the data into acinar or epithelial cell subsets, as determined by cell type annotation. To calculate transcription factor activity score for TP53, we used run_viper() function in decoupleR R Package (v2.6.0) using default parameters (**Figure 4B**).

#### Signature-based annotation of epithelial subpopulations

For signatures derived from mouse datasets, we converted mouse symbols to their corresponding human orthologs using convert_mouse_to_human_symbols() function from nichenetr R package (v2.0.1). We used AUCell R Package (v1.22.0)^132^ to score the gene sets of interest in each cell using default parameters except for aucMaxRank which was set to include 10% of the number of genes in the rankings. This approach led to our initial identification of rare human epithelial states expressing a gene program analogous to the murine progenitor state (**Figure 4A**). Second, we used z-score based estimation of gene signatures, using ADM/duct-like cells from cancer-free donors as reference for computation of the mean and standard deviation of each gene, and excluding genes that were not expressed in this population (leading to zero standard deviation). For each cell, signature scores were computed as the average z-score of signature genes. To visualize signatures across cell populations, we first computed PhenoGraph clusters (k=30) over all ADM/duct-like cells in the data and defined acinar cells and PDAC cells as independent groups for reference. We used these clusters and groups to aggregate signature scores by averaging. **Figure 4B** shows the resulting aggregated signature score matrix, standardized per signature.

## PROCESSING AND ANALYSIS OF SINGLE-CELL MULTIOME DATA

Our previous work established that adoption of distinct premalignant transcriptional states is coupled to genome-wide remodeling of chromatin accessibility landscapes^7^. Integration of chromatin accessibility information and gene expression changes allows inference of regulatory mechanisms underlying the defining features of premalignant states. Given that we found that the tumor-suppressive transcription factor p53 is maximally active in progenitor-like cells, we reasoned that integration of chromatin accessibility data would uniquely reveal the transcriptional consequences of p53 activation during autochthonous tumor suppression.

### Data preprocessing and quality control

Our data consists of two batches of single-cell RNA+ATAC multiome profiling of *Kras*^G12D^+ pancreatic epithelial cells isolated from mice 2 days after receiving the first dose of caerulein to induce pancreatic injury and inflammation. Cells from batch 1 were isolated from one mouse, while cells from batch 2 were isolated from two independent mice. This independent data collection allowed more generalized characterization of chromatin accessibility changes defining each premalignant subpopulation, and accessibility of p53 motifs in progenitor-like cells:

● DACE657_mKate2_multiome: KPfC, Injury day 1 (batch m1)
● p489c_shRen_caer_48h_multiome_1: KC^shRen^, Injury day 1 (batch m3)
● p489c_shRen_caer_48h_multiome_2: KC^shRen^, Injury day 1 (batch m3)

Gene expression count matrix construction and quality control.

We constructed a cell-by-gene count matrix on cellbender ambient-corrected counts, keeping cells that passed both cellbender and emptyDrops filtering. We first preprocessed and filtered each sample independently to remove low quality cells, identify doublets and stromal/immune contaminating cells, as described above (see <u>Low-quality cell removal</u> and <u>Doublet and</u> <u>contaminant identification</u>). Next, we applied hard thresholds to remove cells with low quality control statistics. Specifically, we removed cells with high percentage of mitochondrial mRNA counts (> 20%) or low library size (ln[library size] < 7 for ADM and tuft cells. For other premalignant states, we removed cells with > 20% mitochondrial mRNA counts or ln(library_size) < 7.75. We used these different thresholds to avoid complete depletion of ADM and tuft cells, which had overall lower library sizes than other premalignant states.

#### Chromatin accessibility matrix construction and quality control

We used ArchR v1.0.3^133^ to process raw fragment files obtained through CellRanger Arc, using the createArrowFiles function (*minTSS* = 1, *minFrags* = 100, *maxFrags* = 1000000). We used such permissive thresholds in this first filtering step to allow downstream determination of QC thresholds through the analysis of distributions of QC statistics. Following construction of an ArchR project, we filtered the chromatin accessibility dataset to include only cells that passed QC filters in the gene expression modality. Next, we excluded low-quality cells based on transcription start site (TSS) enrichment and number of fragments in the ATAC modality: log_10_(number of fragments) < 3.23 or TSS enrichment < 6.88, determined from the joint distribution of these two QC statistics in order to capture the density of cells that had many fragments and were enriched for those mapping to TSSs. Our final dataset includes only cells that passed both gene expression and chromatin accessibility QC metrics (**Data S10**).

#### Embedding of gene expression modality

We chose to define subpopulations and to assign cell types based on gene expression, given our better understanding of feature selection and dimensionality reduction criteria, and of marker gene expression in premalignant subpopulations. We processed each batch independently, selecting 3000 HVGs using Scanpy’s sc.pp.highly_variable function (*flavor* = ‘seurat_v3’) on raw cellbender counts. Next we computed PCA, selecting the top 100 PCs explaining 56% of variance in batch m1 and 45% of variance in batch m3. Lastly, we computed a kNN graph (*k* = 30), followed by PhenoGraph clustering (*k* = 30) and visualization with UMAP.

#### Cell typing

Used transcriptional signatures from our previous work^7^ to assign PhenoGraph clusters to states. Specifically, we scored premalignant signatures (progenitor, gastric, ADM, tuft, neuroendocrine), as the average z-scored expression of signature genes in each cell, and visualized and assigned discrete cell-state labels to each cluster. For batch m3, we noted that there were clusters that co-expressed gastric and progenitor signatures, suggesting intermediate states in the continuum. To keep this information in our discretized cell type assignment, we labeled these cells as transitional. Lastly, we identified a small subpopulation of premalignant cells with features of both progenitor-like and ADM cells, labeling these as undefined. The following table summarizes the cell-type fractions in each sample. We note that while we could identify all premalignant states in each dataset, batch m3 is strongly enriched for progenitor-like cells (**Data S10**).

#### Computation of metacells

Data sparsity is a major challenge in single-cell chromatin accessibility analysis, which can be overcome through aggregation at the level of features^134,135^ or cells^108^. Clustering can capture broad differences between populations, but misses heterogeneity between finer cell states. We used SEACells^108^, a density-aware aggregation strategy that aims to partition a single-cell manifold into compact neighborhoods, or metacells^136^, representing discrete, fine-grained cell states, such that intercell variability within each metacell is primarily attributable to technical noise. We computed SEACells v0.3.3 metacells on each batch (*build_kernel_on* = ‘X_pca’, *n_waypoint_eigs* = 10), considering differences in the total number of cells in each dataset (*n* = 50 metacells for batch m1, and *n* = 100 metacells for batch m3). For batch m1, the median metacell size was 46 cells (interquartile range of 28–64 cells). For batch m3, the median metacell size was 63 (interquartile range of 33–94 cells).

### p53 motif analysis

#### Overview of p53 motif

The well-studied canonical p53 responsive element^137,138^ is composed of two direct-repeat decamers (half-site) with a core CATG motif, flanked by three purine-rich and three pyrimidine-rich nucleotides on its 5’ and 3’ ends, respectively^139^. The consensus sequence of a p53 half-site is a palindrome. Thus, each half-site is composed of two adjacent quarter-sites, such that highest binding affinity is achieved when each subunit of a p53 tetramer binds a single quarter site^140,141^.

p53 can also bind variants of the canonical p53 responsive element, including two half-sites separated by a spacer of up to 20 bp (the frequency of validated motifs decays with spacer length)^139^. To capture both binding modes, we used homer^142^ to scan for (1) canonical p53 responsive elements, as defined by the homer database position weight matrix (PWM), and (2) p53 half sites as defined by the ENCODE PWM. While the homer threshold for identifying a full p53 site is 8.59, we used a score of 5 to identify a single half-site (score would be 10 if we added the scores of two adjacent half-sites). This more stringent threshold allowed us to focus on high quality half-sites when predicting non-canonical elements.

#### Tile-based scanning of p53 motif

A common strategy for identifying motifs in chromatin accessibility data is to first call peaks (with default length of 500 bp in ArchR), and then scan these peaks for the presence of DNA binding motifs. We identified two factors that may yield false positive and false negative motif calls using this approach. First, not every motif identified within a peak call is actually accessible, likely because accessible regions may be narrower than the fixed peak length; for example, 12% of canonical motifs that we found within peaks did not meet our subsequent accessibility criteria. In addition, we noticed that peak calls could also miss accessible genomic regions—for example, when proximity between two peak calls precludes the identification of a third peak between them.

To overcome this challenge, we adopted a peak-free motif scanning strategy. Our approach incorporates two principles: we rely on the identification of accessible tiles for motif scanning^143^, and we conduct follow-up analyses of accessibility at the motif level, as opposed to the peak level. To identify accessible tiles, we segmented the genome into 100-bp tiles, quantified the number of insertions in nucleosome-free fragments within each tile (fragment length <= 147 bp), and aggregated accessibility into SEACells metacells, resulting in a metacell-by-tile accessibility matrix. To normalize this matrix, we divided it by the total number of reads in TSS across all cells in each metacell, multiplying by an arbitrary factor of 10,000. Next, we computed the maximum accessibility of each tile over all metacells. We defined accessible tiles as those with a maximum accessibility of at least 0.01. We used this permissive threshold to capture as many p53 motifs as possible, prior to filtering by motif-specific accessibility information.

We noted that some p53 motifs bound in ChIP-seq datasets^65^ were not identified after scanning this set of accessible tiles, and realized that this could occur whenever a motif spanned adjacent tiles. We thus extended each accessible tile by 20 bp at each end, allowing for overlap between adjacent tiles. This final set of genomic regions was used to scan for the full and half p53 sites with the annotatePeaks tool from homer, using the unmasked genome mode.

#### Identification of canonical and non-canonical motifs

To consolidate canonical and non-canonical p53 motifs, we first eliminated redundancies in our motif calls due to the palindromic nature of the p53 full and half-sites, keeping unique motif calls at a genomic coordinate with the highest score, irrespective of strand. We defined two types of non-canonical p53 motifs: (1) pairs of half-sites separated by at most 11-bp spacers, and (2) isolated half-sites, and restricted these to the highest confidence subset by requiring that they localize within 100 bp of p53 ChIP-seq peaks as defined by Kenzelman-Broz et al.^65^. Our consolidated motif table contains 59,133 canonical and 5,036 non-canonical motifs.

#### Estimation of motif coverage per cell type and differential motif coverage analysis

To further refine our motif calls, we computed per-motif chromatin accessibility profiles. For each sample x cell_type pair, we obtained the number of nucleosome-free fragments (fragment length <= 147 bp) that partially or totally overlapped with each motif. We kept raw counts for downstream analysis. To identify differentially accessible motifs, we used DESeq2 on pseudobulked raw read counts per motif in sample x cell type pairs. We compared accessibility between gastric-like and progenitor-like cells: padj < 0.05 and abs(log_2_(fold-change)) > 1.

Overall, we found more motifs that were significantly more accessible in progenitor-like cells, as opposed to gastric-like cells. Furthermore, motifs that were differentially open in progenitor-like cells were enriched in motifs with evidence for p53 binding in public ChIP-seq data^65^, as compared to those with increased accessibility in gastric-like cells (p = 4.80E-38, Fisher’s exact test).

#### Identification of motif accessibility threshold

To identify a threshold for motif accessibility in progenitor-like cells, we used triangle thresholding on the distribution of normalized log-transformed motif accessibility scores in progenitor-like cells (pseudocount = 0.00001). This resulted in a threshold of 0.01175 normalized insertion counts on pseudobulked progenitor-like cells.

### Identification of *in vivo* p53 program

#### Motif to gene mapping

To identify genes potentially regulated by p53 in progenitor-like cells, we computed a motif-to-gene mapping. We first identified a consensus TSS associated with each gene symbol in Ensembl GRCm38 102, considering transcripts annotated as protein_coding, lncRNA, IG_C_gene, Mt_rRNA, Mt_tRNA, TEC, TR_C_gene, TR_D_gene, and TR_J_gene, thus excluding transcripts annotated as pseudogenes or predicted to yield nonsense-mediated decay products. Then we identified the median genomic coordinate corresponding to the transcript start (for transcripts in the + strand), or end (for transcripts in the - strand) over all transcripts corresponding to a gene symbol, thus identifying a consensus coordinate for the TSS. Lastly, we identified all p53 motifs accessible in progenitor-like cells that lie within 20 kb of each TSS. Based on the accessibility threshold identified above (Identification of motif accessibility threshold), a total of 8,181 p53 motifs were identified as accessible in progenitor-like cells.

**Data S4** contains the list and metadata of the p53 motifs that we identified.

#### Differential gene expression

We leveraged our differential gene expression results (see Processing and analysis of *p53* knockdown data - Differential gene expression) to identify genes upregulated in *p53*-proficient progenitor-like cells relative to *p53*-proficient gastric-like cells. Using a threshold of log_2_(fold-change) > 0.5 and adjusted p < 0.005, we identified 2,090 genes upregulated in progenitor-like cells relative to gastric-like cells.

Next, we identified genes downregulated in progenitor-like cells upon *p53* knockdown. To focus on the direct consequences of *p53* knockdown within a given cell state, as opposed to differences linked to cell-state transitions, we compared *p53-*proficient and deficient cells at equivalent positions along the G–P diffusion component (see Processing and analysis of *p53* knockdown data - Differential gene expression and **Figure S6E**). Using a threshold of log_2_(fold-change) < −0.5 and adjusted p < 0.005, we identified 298 genes downregulated in *p53-*deficient cells upon *p53* loss, that were also present in our Ensembl annotation table (for example, the gene encoding for our shCtrl RNA was excluded).

#### Integration with public datasets

Beyond the data generated in this study, we examined multiple sources of evidence to assess the list of candidate p53 targets: genes that we previously showed to be upregulated in PDAC cells *in vitro* upon *p53* restoration^64^; genes bound by p53 in irradiated MEFs^65^; and p53 targets reported in multiple publications^65,84,102,103,139,144–147^, allowing us to identify potentially novel p53 targets in our experimental system.

#### Categorization of p53 targets

By integrating three sources of information (presence of accessible p53 motif in progenitor-like cells, upregulation in progenitor-like cells relative to gastric-like cells, and upregulation upon *p53* restoration *in vitro*), we categorized genes that decrease in expression upon *p53* knockdown into 8 different classes (**Figure S6G**). Note that there are previously reported p53 targets in each category, including those groups that lack evidence for p53 motif accessibility in progenitor-like cells, suggesting that some genes may be regulated by accessible motifs that lie beyond the 20 kb distance threshold that we used for motif-to-gene mapping.

#### Pathway analysis

To further interpret our list of candidate p53 targets, we conducted pathway analysis using Enrichr^107^ and the ChEA_2022 database, which compiles results from ChIP-seq datasets across transcription factors (**Figure S6I**). This analysis highlighted enriched p53 binding in all groups of genes for which we identified a candidate accessible motif in their genomic neighborhood. In addition, p63 binding appeared as a secondary signal, consistent with the shared motif features between p53 family members^107,148^.

## QUANTIFICATION OF qPCR DATA

We used a two-tailed paired t-test with default parameters in Prism v10.6.1 to test for differences in mRNA expression in premalignant organoids treated with recombinant TGF-β, relative to untreated ones (*n* = 3 paired biological replicates per organoid line and condition).

## SPATIAL DATA PROCESSING AND ANALYSIS

### Immunofluorescence image processing and quantification

Our immunofluorescence image collection and analysis aimed to quantify signaling proteins in progenitor-like or other premalignant cells (i.e., p53 and phospho-ERK (p-ERK)), and progenitor-like epithelial cells in response to perturbing the premalignant pancreas (i.e., HMGA2- or VIM-positive cells). For the latter, we collected whole tissue scans, reasoning that this strategy would most faithfully quantify the progenitor-like cell population, by minimizing bias and variability stemming from spatial heterogeneity in progenitor-like lesions (documented in **Figure S7C**). To quantify how premalignant states differ in p53 abundance, we selected FOVs containing lesions rich in progenitor-like cells (MSN+) and spatially adjacent lesions devoid of this state (MSN-), providing an internally controlled setup for comparison. All analyzed tissues were collected at 20x magnification with a pixel size of 0.34 μm px^-1^.

To process and analyze whole tissue scans, we divided images in non-overlapping FOVs of 500 x 500 px for *p53* perturbation, HMGA2 and VIM datasets, or 1000 x 1000 px for KRAS inhibitor datasets. We used smaller FOVs for the *p53* perturbation cohort to minimize the inclusion of large empty regions in images derived from small fractions of the pancreas.

#### Generation of nuclear and whole-cell segmentation masks

To quantify HMGA2-positive cell abundance, p53 protein and p-ERK intensity, we used nuclear segmentation masks generated with Mesmer, implemented in the deepcell python package (v 0.12.9)^149^. This deep learning method was pretrained on a large number of manually annotated cell and nuclear masks from diverse imaging modalities and biological sources, aimed at generalizing mask prediction in new data, and it uses boundary predictions with a watershed approach to refine segmentation masks. For nuclear segmentation, we used the standard rolling ball background subtraction approach (*radius* = 10 px) on DAPI images, which we then fed to Mesmer with *compartment* = ‘nuclear’, *preprocess_kwargs* = {*’percentile’*: 99.9, *’threshold’*: True, *’normalize’*: True, *’kernel_size’*: 10}, and postprocess_kwargs_nuc = {*’maxima_threshold’*: 0.001, *’maxima_smooth’*: 1, *’interior_threshold’*: 0.1, *’interior_smooth’*: 0, other parameters as default}. Non-default parameters were manually tuned to recover accurate nuclear segmentation results in our datasets.

As VIM localizes outside the nucleus, we undertook whole-cell segmentation using Cellpose (v3.0.6)^150^ with DAPI (nucleus) and E-cadherin (membrane) stains to quantify VIM+ epithelial cells. Visual inspection revealed that Cellpose generated better whole-cell segmentation masks than Mesmer in our data. Like Mesmer, Cellpose is a deep learning method that is pretrained on a large collection of ground-truth masks; it leverages both cytoplasmic and nuclear stains to predict cell boundaries. Although we were interested in cells with mesenchymal properties, we noticed that progenitor-like cells with upregulated VIM expression retain membrane-localized E-cadherin. This suggests that our cells of interest lie at an intermediate point in the epithelial-to-mesenchymal spectrum, and enables the use of an epithelial marker such as E-cadherin to generate masks. Following background subtraction with the rolling ball approach (*radius* = 10) on DAPI and E-cadherin channels, we normalized images (saturating at the 99th quantile of the intensity histogram), and blended these channels into an RGB image. Next, we applied the pretrained segmentation model (*model_type* = ‘cyto3’) to generate segmentation masks using the model.eval function of the cellpose package with *diameter* = None, *channel_axis* = 0, *normalize* = True, *channels* = [2,3], *flow_threshold* = 0, and *cellprob_threshold* = 0.

Visual inspection of both Mesmer and Cellpose masks confirmed that these approaches better adapt to heterogeneous cellular densities and signal intensities in the tissue, compared to traditional approaches that rely on thresholding, morphological operations and watershed-based segmentation.

#### Signal quantification and quality control

We used segmentation masks to quantify fluorescent signal intensity, applying standard rolling ball background subtraction for markers with subcellular and stereotyped localization (e.g., HMGA2, VIM and p53). We avoided this background subtraction for p-ERK and TNC, as a single radius may fail to adequately capture the diverse sizes and geometries of epithelial and stromal structures that they mark; instead, we measured raw intensities at this step, and leveraged the full distribution of measured states to conduct background subtraction at the whole-slide or single-FOV level (see below).

In addition to measuring fluorescence intensity in the segmentation mask, we quantified geometric parameters such as area and solidity (a measure of roundness, calculated as the fraction of the convex hull of each segmented object covered by the actual mask). Geometric parameters are particularly helpful for QC, as they provide information about putative segmentation errors. They were computed using the regionprops function of the measure module in the python scikit-image package (v0.22.0). The resulting multidimensional representation of each cell in the dataset consisted of both intensity and geometric measurements, allowing downstream QC and *in silico* isolation and quantification of specific cell subpopulations.

Geometric features can be used to identify segmentation errors; for example, pairs of adjacent cells that segmentation fails to separate tend to have increased area, and decreased solidity. We used morphological parameters to identify putative low quality cells, filtering out cells with low solidity (< 0.8) or high nuclear area (>750 px), which excluded an average of 3% of cells in the data. **Data S6** summarizes results from QC filtering in whole-slide scan datasets.

#### Identification of epithelial cells

In our mouse models, RFP and GFP are activated for lineage tracing upon Cre-mediated recombination and doxycycline induction of shRNA alleles, thus serving as proxies for oncogenic KRAS activation (see **Mouse model genetics**). To identify premalignant epithelial cells, we computed the distribution of log-transformed (pseudocount = 1) average nuclear GFP or RFP intensity. Log-transformation of intensity values compresses the tails of the distributions, making bimodality more apparent and facilitating the identification of thresholds to isolate marker-positive cells. We used otsu thresholding on the log-transformed intensity to binarize the signal, resulting in the identification of premalignant epithelial cells (29% of cells, on average, across datasets). **Data S6** summarizes results of epithelial cell identification in whole-slide scan datasets.

#### Quantification of HMGA2- and VIM-positive cells

We used HMGA2 and VIM expression to quantify the abundance of progenitor-like premalignant cells in premalignant tissues. Similar to our treatment of GFP and RFP signals, we log-transformed (pseudocount = 1) the intensity distributions of HMGA2 and VIM followed by signal binarization using the Otsu thresholding method, capturing the second mode of the distribution. We summarized HMGA2 and VIM measurements as the average number of positive epithelial cells per field of view in each slide, and tested for differences between experimental conditions (KC^shp53^ and KC^shCtrl^) with a two-tailed Wilcoxon rank sum test. **Data S6** provides detailed information regarding sample metadata and source data shown in **Figures 6B** and **7B,E**.

#### Quantification of p-ERK levels

We assessed the effectiveness of oncogenic KRAS inhibition *in vivo* through staining and quantification of p-ERK, a canonical downstream effector on MAPK signaling. To do so, we calculated the average p-ERK signal and non-epithelial cells per tissue. We used the average p-ERK intensity in non-epithelial cells as our estimate of the background signal, subtracting it from our measurements of p-ERK intensity in epithelial cells. The use of non-epithelial cells to estimate a global background per sample is motivated by selective p-ERK upregulation in the *Kras*^G12D^+ cells; on the other hand, non-cell-autonomous effects leading to upregulation of MAPK signaling in non-epithelial cells would lead to underestimation of signal intensity in the epithelium, and thus our results should be interpreted as a lower bound estimate of the difference in p-ERK engagement between vehicle and MRTX1133-treated samples. We tested for differences between experimental conditions using a two-tailed Wilcoxon rank sum test.

**Data S6** provides detailed information regarding sample metadata and source data shown in Figure S5A.

#### Quantification of p53 levels

Our single-cell data revealed the selective upregulation of p53 target genes in progenitor-like premalignant cells during both spontaneous and injury-induced tumorigenesis (**Figures 1G,K, 7D,G** and **S6D–I**). Protein stability is a key mechanism by which p53 is regulated^151^; thus, we quantified protein levels in premalignant cells, staining for p53 and the progenitor-like marker Msn separately in serial sections to avoid antibody isotype incompatibility. We first identified progenitor lesions (Msn+) in one section, considering only epithelial structures and remaining blind to p53 signal, and manually annotated the equivalent region in the adjacent section. Next, we quantified p53 abundance in epithelial cells from all Msn+ or all Msn-lesions. As p53 staining has a lower signal-to-noise ratio than cell-state markers such as Hmga2, Msn, Vim and GFP, we conducted local signal normalization to control for systematic intensity bias across the tissue. Specifically, for each FOV, we normalized nuclear p53 intensity in epithelial cells by its average intensity in non-epithelial cells, then computed the average normalized signal in MSN+ or Msn-epithelial cells for each tissue sample. We used a two-tailed Wilcoxon rank sum test to identify significant differences in p53 levels between *Msn*+ and *Msn*-cells. **Data S6** provides detailed sample metadata and source data information for **Figure S2C.**

#### Quantification of TNC levels

Acute oncogenic Kras inhibition led to collapse of the progenitor niche within 48 h of treatment, as evidenced by depletion of microenvironmental transcriptional states associated with progenitor-like cells (**Figure 6C,D**). To test whether remodeling events were reflected at the protein level at these early time points, we quantified TNC, an injury- and cancer-associated ECM component that is upregulated in activated myofibroblasts in the progenitor niche. We opted for segmentation-free pixel quantification in the entire tissue section, given that TNC generates fibers not strictly associated with cellular or nuclear segmentation masks.

Furthermore, we normalized by GFP-positive signal, as it represents the area of pancreatic parenchyma in the image. To identify positive and negative pixels, we analyzed intensity distributions of TNC and GFP per tissue section. First, we set a hard threshold of ln(pixel intensity) > 5.5, empirically determined to avoid empty areas in the image for both markers. Next, we computed the histograms of log-transformed TNC and GFP intensity values. We fit a Gaussian centered on the first mode of the intensity distributions using scipy.optimize to determine a background distribution. Using the mean and standard deviations of our background estimates, we standardized the intensity distributions, effectively equalizing the background distribution in all tissue slides. Lastly, we used triangle thresholding to identify pixels positive for TNC or GFP. We normalized the fraction of TNC-positive pixels by the fraction of GFP-positive pixels, and tested for differences between conditions using a two-tailed Wilcoxon rank sum test. **Data S6** provides details of sample metadata and source data in **Figure 6E**.

#### Quantification of the effect of macrophage depletion in premalignant pancreas

We used immunofluorescence to quantify the outcome of anti-CSF1R treatment on the abundance of macrophages in the premalignant pancreas. For each biological replicate (IgG control *n* = 4, anti-CSF1R *n* = 3), we collected five 2000 x 2000 px FOVs of premalignant tissue stained for F4/80, a macrophage marker, and GFP, a proxy for activation of oncogenic Kras in the premalignant epithelium. We first normalized DAPI images by clipping the pixel intensity distribution at the 10th and 99th percentiles, followed by scaling to [0,1]. Next, we used Cellpose-SAM to obtain nuclear segmentation masks for each FOV and quantified the average intensity of GFP and F4/80 in each nucleus, using a rolling ball background subtraction strategy (*disk size* = 10 px) to enhance the signal-to-noise ratio in the F4/80 channel. We excluded nuclei with areas less than 20 px or greater than 750 px, and with solidity below 0.85. In addition, we used triangle thresholding to remove nuclei at the lower tail of the log-transformed (pseudocount = 1) DAPI intensity distribution. These filters aimed to mitigate the effect of false-positive nucleus identification or segmentation errors (merged cells). Following measurements of intensities, we used triangle thresholding on the log-transformed GFP intensity distribution to identify premalignant epithelial cells. Next we used triangle thresholding on F4/80 nuclear intensity distributions (pseudocount = 1) derived from GFP negative cells to identify F4/80 positive cells. We computed the fraction of non-epithelial cells that were positive for F4/80 per biological replicate, and tested for differences in the median fraction of F4/80 positive cells as a function of treatment (IgG or anti-CSF1R) with a Wilcoxon rank sum test (**Figure 5F**).

To assess the effect of anti-CSF1R treatment on HMGA2-positive cell prevalence, we quantified the fraction of this subpopulation in the entire tissue sample (for reliable quantification, large tissue regions need to be analyzed to overcome the spatial heterogeneity of progenitor-like cells). We followed the strategy above (Image processing and quantification) for generating image patches and nuclear masks using Mesmer, and for identifying GFP-positive premalignant epithelial cells. We noticed that the mode of log-transformed HMGA2 nuclear intensity distributions differed subtly between biological replicates. Since at the time of harvesting HMGA2-positive cells are rare in the premalignant pancreas (3 weeks post-injury; <10% of cells following caerulein-induced pancreatitis), these subtle differences can cause thresholding methods, which aim to capture the upper tail of the distribution, to produce false-positive or false-negative calls for HMGA2-positive cells. To equalize the modes of these distributions, we fit a Gaussian to the first mode of the HMGA2 log-transformed intensity distribution for each biological replicate using the curve_fit function from the scipy.optimize module (v1.11.4) and standardized the distribution of each replicate based on the mean and standard deviation estimates. Following this correction, we quantified the fraction of HMGA2-positive cells out of all the GFP positive cells per biological replicate, and tested for differences in the median HMGA2 fraction using a Wilcoxon rank sum test (**Figure 5G**).

Visual inspection of the imaging data also revealed that HMGA2-positive cells tended to occur in larger clusters in IgG control samples, as opposed to the single-cell patterning that prevailed in anti-CSF1R treated samples. To systematically analyze the effect of anti-CSF1R treatment on the cluster size distribution of HMGA2-positive cells, we computed a spatial neighbor graph, connecting nuclei that lied within 20 µm, thus capturing the immediate vicinity of each cell. Next, we identified connected components in the spatial neighborhood graph, and annotated each HMGA2-positive cell with the size of their connected component. To visualize the distribution of neighborhood sizes, we binned all HMGA2-positive cells according to the size of their HMGA2 cluster, using powers of 2 as bin breaks, and we tested for differences in the distribution of neighborhood sizes associated with each HMGA2-positive cell with a Kolmogorov-Smirnov test (**Figure 5H**).

### Processing and quantification of COMET data

The Lunaphore COMET platform can stain up to 40 proteins in the same tissue slide, and allows multiple antibodies with the same host IgG isotype to be used, as it relies on cycles of primary and secondary staining as opposed to fluorophore-conjugated primary antibodies. This capability was critical for our study, since multiple rabbit antibodies (MSN, HMGA2, ANXA10, cleaved caspase 3, phospho-ERK and VIM) together accounted for important dimensions of cell-state heterogeneity and signaling in the premalignant pancreas.

#### Dataset details

Our dataset is composed of three tissues from premalignant stage mice treated with the oncogenic KRAS inhibitor MRTX1133, and euthanized 4 h after inhibitor dose, and two from corresponding vehicle-treated mice. Data was collected both during *in vivo* experimentation and imaging in two batches:

Batch 1:

● JR-sp-15-010 (4h after MRTX1133 30mg/kg)
● JR-sp-15-011 (4h after vehicle)

Batch 2:

● JR-sp-40-11 (4h after vehicle)
● JR-sp-40-13 (4h after MRTX1133 30mg/kg)
● JR-sp-40-17 (4h after MRTX1133 30mg/kg)

Each sample captures 12.5 mm x 12.5 mm of tissue, resulting in a 44643 x 44643 px image with a pixel size of 0.28 μm px^-1^. All downstream processing, including data inspection and interaction, was conducted on images resulting from built-in background subtraction in the COMET system. This procedure relies on subtracting an autofluorescence image acquired from capturing unstained tissue with the same wavelength as the stained image, thus improving signal-to-noise ratios. No additional background subtraction was conducted thereafter.

#### Segmentation

We prioritized the accurate identification of an epithelial cytoplasmic mask during segmentation, given our focus on this cellular compartment. Specifically, we aimed to quantify the correlation of progenitor-like state protein markers, and the fraction of cells that undergo apoptosis upon oncogenic KRAS inhibition as functions of premalignant cell state. We first used the SpatialData^152^ framework to rapidly extract images from different channels for the same tissue, storing multiscale images in the zarr format. Next, we used Cellpose-SAM to segment epithelial cells in a two-dimensional image composed of a nuclear signal (DAPI) and an epithelial membrane marker (E-cadherin). As input to the Cellpose model, we used images downscaled by a factor of 0.5, which improved computation speed and memory usage, while maintaining accuracy in segmentation, as assessed visually. We clipped DAPI and E-cadherin images on the 10th and 99th percentiles of their pixel intensity distributions, normalizing pixel intensities to [0,1]. We used Cellpose-SAM (v4.0.7) with the following parameters: *batch_size* = 64, *flow_threshold* = 0.4, *cellprob_threshold* = 0, *normalize* = {“tile_norm_blocksize”: 0} to obtain a segmentation mask. Note that cytoplasmic masks are only valid for epithelial cells; other cellular compartments had a nuclear mask only. Lastly, we rescaled the masked image to match the original image size and saved it as a compressed numpy object. In parallel, we computed a segmentation mask for nuclei only, using the DAPI image as input to Cellpose-SAM, with the same parameters. The cytoplasmic and nuclear segmentation masks we obtained for each tissue image are labeled images; i.e., pixels associated with each cell have a specific value which acts in lieu of a cell identifier.

#### Consolidation of nuclear and cytoplasmic masks

Because some markers largely localized to the nucleus (e.g., HMGA2, p19) and others to the cytoplasm and membrane (E-cadherin, MSN, CC3), we reasoned that it was important to quantify signals in both cellular compartments, while maintaining consistent cell labels between segmentation masks. To consolidate nuclear and cytoplasmic masks, we first binarized the nuclear segmentation labeled image, so that pixels corresponding to a nucleus had a value of 1, and non-nuclear pixels had a value of 0. Next, we multiplied this binarized nuclear mask with the labeled cytoplasmic mask. This resulted in a new labeled mask image representing a nuclear segmentation with identifiers consistent with our cytoplasmic mask. Since some cells may disappear through this operation, e.g., those for which a nucleus was not independently identified during segmentation of the DAPI image, we removed all cells from the original cytoplasmic mask that lacked a corresponding cell in the new labeled nuclear mask. This process resulted in pairs of nuclear and cytoplasmic masks for each tissue sample, with consistent identifiers in both segmentation modalities.

#### Intensity quantification

We used the regionprops function of the scikit-image package (v0.25.2) to quantify the mean nuclear and cytoplasmic intensity for each fluorescent channel, as well as morphological properties, such as area and solidity, keeping a record of the centroid of each cell. In the case of p19, which forms discrete foci in the nucleus, we extracted additional parameters that allowed us to identify the presence of intense foci within a mask. Specifically, we obtained the median pixel intensity within each mask, as well as the 95th percentile of the masked pixel intensity distribution (termed ‘top pixels’). Cells with foci will have a higher top pixel to median pixel intensity ratio.

#### Censoring low quality regions and lymph nodes

During raw data inspection, we noticed several properties of our images that could distort downstream analysis. First, imperfect alignment between autofluorescence images and stained images led to remnant wedge-shaped autofluorescence patterns, particularly associated with red blood cells. Second, densely-packed lymph nodes also had non-negligible remnant autofluorescence signal. Cells within the lumen of epithelial structures, which were also more likely to exhibit cleaved caspase 3 activation, had autofluorescence patterns that made them hard to distinguish between epithelial and non-epithelial cells. Lastly, out-of-focus regions, tissue folds and bubbles could also influence downstream quantification. To clean up our data, we manually identified these regions and applied a binary censor mask.

#### Cell typing

To identify epithelial cells for downstream analysis, we leveraged the multiplex nature of our data, rather than relying on single intensity thresholds for epithelial markers such as E-cadherin or GFP.

- Feature selection: Similar to Cell DIVE images, we conducted coarse cell typing on a sample-by-sample basis. We used log-transformed and z-scored intensity values of a select feature set: DAPI_nucleus, dpERK_cyto, dpERK_nucleus, MSN_cyto, HMGA2_nucleus, LGALS4_cyto, LGALS4_nucleus, E-cadherin_cyto, GFP_cyto, GFP_nucleus, ANXA10_cyto, ANXA10_nucleus, VIM_cyto, VIM_nucleus, p19_nucleus_intensity_ratio, p19_nucleus, CC3_cyto, CC3_nucleus, CPA1_nucleus, CPA1_cyto. Of particular importance was the inclusion of DAPI intensity in the feature set, as we consistently identified a cluster of cells with DAPI-low or negative signal that may correspond to regions without cells.

- Dimensionality reduction and clustering: We conducted PCA, keeping the top PCs based on kneepoint (typically 7 PCs capturing up to 90% of the variance), followed by PhenoGraph clustering on a kNN with *k* = 30 constructed on PC space.

- Cell type assignments: We examined the average expression of select protein markers in each PhenoGraph cluster to determine those that expressed epithelial-specific markers (E-cadherin, GFP, LGALS4, ANXA10, HMGA2 and CPA1). We excluded clusters that expressed E-cadherin but lacked GFP expression, indicating cells without *Kras*^G12D^ expression. This identified epithelial cells within each tissue region.

#### Identification of epithelial subsets

Because our dataset was gathered in two batches, we combined samples from the same batch for refined cell typing, reasoning that it would improve consistency in cluster assignment between samples while mitigating variability due to batch effects. We conducted clustering using the procedure described in the previous step (<u>Cell typing</u>), with the exception that we excluded VIM_cyto and VIM_nucleus from our feature set during embedding. Visualization of the average marker expression per cluster allowed us to identify subclusters of progenitor-like and gastric-like states. Specifically, we identified a set of gastric-like cells that expressed LGALS4, but not ANXA10, as well as a set of clusters that expressed both markers. In addition, we identified a set of clusters that expressed MSN, but relatively low levels of HMGA2 and p19, as well as a set of clusters that expressed the three markers. While we could distinguish between these cell states in our clustering, different types of progenitor-like cells still colocalized to the same lesions, suggesting that they are phenotypically related.

#### Correlation between progenitor markers

To assess the correlation of progenitor and gastric-like markers at the protein level, we computed and clustered a correlation matrix on six epithelial markers that spanned major premalignant cell states: progenitor (MSN, HMGA2 and p19), gastric (LGALS4, ANXA10) and ADM (CPA1). This analysis confirmed that, similar to patterns at the mRNA level (**Figure S1E**), these markers correlated in single cells at the protein level (**Figure S2A,B**).

#### Quantification of cleaved caspase 3 after oncogenic KRAS inhibition

We used triangle thresholding on log-transformed z-scored intensity distributions to identify CC3-positive cells in each sample. Next, we computed the fraction of each epithelial cluster or subcluster that were either CC3-positive or negative. We tested for differences in CC3 positivity between cell states with a one-sided Wilcoxon rank sum test. We note that the maximum significance achievable for this test with two groups of *n* = 3 is 0.049 (single asterisk in **Figure S5E**).

### Quantification of human pancreatitis Cell DIVE data

We sought to quantify the prevalence of KRT17+ cells and microenvironmental properties of the 67 high quality tissue cores from patients with pancreatitis in our Cell DIVE dataset. Our panel emphasized epithelial and immune markers, which work robustly with this technology in our experience, as well as fibrosis and reactive stromal markers such as TNC, aSMA, Collagen I and PDPN. Given the high quality of segmentation achievable in both acinar and non-acinar epithelial cells with markers such as beta-catenin, we decided to first characterize segmented epithelial cells and then conduct cell-type-free quantification of marker expression throughout the tissue in relation to epithelial heterogeneity.

Our analysis consisted of (1) identifying individual tissue cores; (2) segmenting cells; (3) quantifying signal intensity per core; (4) applying cluster-based identification of cell types; (5) identifying KRT17+ epithelial cells; (6) censoring low quality regions and quantifying TNC in a spatially restricted manner; and (7) applying distance-based quantification of marker gene expression relative to epithelial cell states.

#### Identification of individual tissue cores

We conducted segmentation and signal quantification on each tissue core separately, allowing us to control for differences in signal intensity across samples, which is particularly important for commercial tissue arrays. We measured the typical radius of a tissue core as 2950 px and downscaled the DAPI image from our first imaging cycle by a factor of 200, then blurred this image using a fast fourier transform convolution with a disk whose radius corresponded to the downscaled radius of a tissue core. We identified the centroid of each tissue core as the local maximum of the blurred DAPI image, visually verifying that there was one centroid per patient sample, and defined a square bounding box centered at each centroid. The width and height of the bounding box were 6300 px, accounting for the diameter of each core, and a 200-px rim on each side to buffer against heterogeneity in the size of each sample. Lastly, we used these bounding boxes to extract an image for each tissue core and channel combination, saving them as numpy objects.

#### Cell segmentation

We used Cellpose v4.0.7 to segment cells. This version of the Cellpose deep learning framework incorporates the backbone of the Segment Anything Model (SAM) foundation model (https://doi.org/10.48550/arXiv.2304.02643). We used an RGB image with nuclear and cytoplasmic signal as input to the Cellpose model. The nuclear signal was derived from the maximum projection of all DAPI images throughout imaging cycles. To account for differences in image intensities between cycles, we normalized image intensities prior to computing the maximum projection, by clipping each image at the 50th and 97.5th pixel intensity percentiles, followed by rescaling pixel intensities to [0, 1]. The cytoplasmic image was constructed similarly, but from the maximum intensity projection of two channels (beta-catenin and CD45)—chosen because they exhibited high signal-to-noise, sharp cytoplasmic boundaries, and labeled all epithelial and immune cells with similar intensities. This stood in contrast to other markers, such as E-cadherin, which differed in intensity between acinar and non-acinar epithelial cells.

Following the generation of a nuclear/cytoplasmic RGB image, we used the Cellpose SAM model to identify cell boundaries (*flow_threshold* = 0.4, *cellprob_threshold* = 0.0). The segmentation for other major cell types (e.g., fibroblasts, endothelial, mural cells) was restricted to nuclear masks because we lacked membrane stains for them.

#### Quantification of signal intensity per core

We quantified the average intensity of each fluorescent channel for each segmented cell. Since the Cell DIVE platform obtains the autofluorescence signal for each channel right after each dye inactivation round, and subtracts this signal from the subsequent imaging round, we decided not to conduct any further background subtraction. In addition, we recorded the centroid information from each cell, keeping a record of the cell position within each core (local coordinates) and in the whole slide (global coordinates).

#### Cluster-based identification of cell types

Our dataset is composed of a cell-by-marker intensity matrix. To identify major cell types, we applied methods developed for single-cell transcriptomics. First, we visually inspected the signal from each marker, identifying those that showed robust and variable expression throughout the tissue. Our final feature set included the following markers: Beta-catenin, CD56, CryAB, Ecad, EGFR, Glut1, H3me3K9, HLAdr, HMAG1, IRF1, KI67, KRT17, MHCI, MSN, NaKatpase, p16^INK4A^, p21, p53, PanCK, pERK, pSTAT3, SOX2, TUBB3, uPAR, YAP, ARG1, aSMA, CC3, CD11b, CD11c, CD163, CD20, CD31, CD3e, CD4, CD45, CD68, CD8a, COL1A1, GZMB, IBA1, MPO, PD1, Podoplanin, Pselectin, TNC, VIM. Next, we log-transformed our intensity matrix (pseudocount = 1), and z-scored intensities over each marker. We computed PCA on our standardized intensity matrix, selecting the number of top PCs based on the kneepoint criterion. The median number of PCs selected were 11 PCs, explaining 78% variance (**Data S3** specifies the number of PCs selected and the variance they explained in each sample). Lastly, we constructed a kNN graph on the cell-by-PCs matrix (*k* = 10) and identified clusters using PhenoGraph (*clustering_algo* = ‘leiden’). We grouped cells by PhenoGraph cluster, aggregated the z-scored intensity of each marker, and assigned cells to the following groups:

- Epithelial beta-catenin positive (high for beta-catenin, low for pan-cytokeratin), marking acinar cells and islets. Identification of this group of epithelial cells was important for downstream analyses, as it allowed true fluorescence to be distinguished from high autofluorescence in acinar cells.

- Epithelial pan-cytokeratin high (PanCK^high^) (high for beta-catenin, high for pan-cytokeratin), marking ductal and metaplastic epithelial cells.

- Immune (expressing combinations of CD45, MPO, IBA1, CD68, CD20, CD3e, CD8a)

- Stromal (expressing high levels of Collagen I, aSMA or CD31).

In addition, we identified a group of cells with high levels of GLUT1 that otherwise showed mixed cell type phenotypes and excluded these cells from subsequent analyses.

Following assignment of cell types, we visualized their spatial organization as a scatter plot, verifying that it matched the global patterning observed in the raw microscopy image.

#### Identification of KRT17+ epithelial cells

KRT17 is a marker of basal-like cells in human PDAC with high signal-to-noise in IF assays. Since progenitor-like cells upregulate basal-like signatures, and KRT17 was upregulated in a cluster of human epithelial cells that shared transcriptional features with progenitor-like premalignant cells, we reasoned that this marker could anchor our investigations of phenotypic heterogeneity in human tissues. To identify KRT17+ and KRT17-epithelial cells, we used triangle thresholding on the log-transformed intensity distribution of this marker over all PanCK^high^ cells in the dataset. We note that restricting the identification of KRT17+ cells to the PanCK^high^ epithelial fraction was important to avoid inclusion of KRT17-acinar cells that nonetheless passed the intensity threshold due to autofluorescence.

#### Spatially restricted quantification of TNC

Consistent with its role as an extracellular matrix component, TNC showed clear fibers in IF images. However, it suffered from low signal-to-noise, and acinar cells in particular contained autofluorescence regions with intensities in the range of true TNC signal. Given the salience of this marker in the mouse progenitor niche, we conducted a refined quantification to minimize autofluorescence signal capture. Specifically, we manually outlined tissue regions with TNC fibers in QuPath, blinding the specification of these regions to any other fluorescent marker, and exported these annotations as a binary mask. Next, we extracted pixels in the TNC image residing within these binary masks, computed the distribution of log-transformed pixel intensity, and used the triangle method on the pixel intensity distribution to identify a threshold for TNC positivity. This resulted in a binary image where non-zero pixels were deemed TNC positive. We used the spatial join operation (sjoin) from the geopandas package (v1.1.1) to count the number of pixels within a 30-μm radius of a cell centroid, thereby quantifying the abundance of TNC immediately surrounding a cell.

#### Censoring low quality regions

Certain regions of tissue contained folds, low quality DAPI staining, or artifacts resulting from misalignment between sequential imaging rounds for specific channels. To avoid analyzing these regions, we created binary censor masks in QuPath, and excluded cellular centroids from them. In total, we censored 3% of cells in the dataset within 10 tissue cores. The number of censored cells ranged from 4% to 30%.

#### Local neighborhood composition around KRT17+ or KRT17-epithelial cells

Our data exploration revealed that KRT17+ epithelial cells had a tendency to localize at the edge of acinar lobes, and thus lay closer to stromal and immune cells. To test this systematically, we asked whether this fraction was higher in the neighborhoods of KRT17+ PanCK^high^ cells (hereafter termed KRT17+ neighborhoods), relative to the neighborhoods of their KRT17-counterparts (neighborhood radius = 60 μm). Because the prevalence of KRT17+ epithelial cells differed widely between tissue cores (**Figure 4D**), we subsampled cells such that for each patient, we compared an equal number of KRT17+ or KRT17-neighborhoods. In addition, we excluded tissue cores with fewer than 15 KRT17+ epithelial cells, to minimize noisy signals stemming from low cell numbers (6 cores excluded). For each tissue core, we computed the average immune/stromal fractions of KRT17+ or KRT17-neighborhoods, and tested for differences in the median immune/stromal fractions between these two types of neighborhoods across all patients with a Wilcoxon rank sum test (**Figure 4E**).

#### Distance-based quantification of marker gene expression relative to epithelial cell states

We aimed to quantify protein marker levels as a function of distance from KRT17+ or KRT17-epithelial cells, but—because KRT17+ cells were located at the edge of pancreatic lobes—wanted to avoid capturing expression differences due to the location of the epithelium in relation to the stromal/parenchyma boundary. We thus grouped cells into tumor microenvironment (TME) bins based on the stromal/immune fraction of each PanCK^high^ epithelial cell neighborhood (neighborhood radius = 60 μm), with bins ranging from 0 to 1 in 0.1 increments. For each core, we subsampled either KRT17-cells or KRT17+ cells (whichever were more numerous) to render their numbers equivalent for each TME bin, and discarded KRT17+ cells which lacked a KRT17-cell at an equivalent TME bin within the same patient. Collectively, this allowed us to compare equal numbers of KRT17+ and KRT17-cells from each patient, while also comparing cells in these two groups that had similar local microenvironment composition at the coarse cell type level.

We then constructed a spatial neighbor graph connecting all cells with centroids within 200 μm, and computed the average expression of each marker as a function of distance for each individual cell, generating an n_cells x distance x protein_marker matrix. Averaging the distance x marker matrix over all KRT17+ or KRT17-cells resulted in two matrices that summarized distance-dependent changes in protein expression in these two epithelial cell types. We visualized these matrices after jointly standardizing KRT17+ and KRT17-matrices over all distances for each protein marker (**Figure 4G**).

### Quantification of progenitor niche markers in Cell DIVE data

We quantified key epithelial and microenvironmental markers in the progenitor niche at the protein level to complement our comprehensive spatial transcriptomic data and to determine niche decay as a function of distance from progenitor-like cells. This effort yielded a quantitative characterization of the progenitor niche and the typical length scale over which it operates, in both premalignant and malignant mouse tissue.

We used the Cell DIVE platform to quantify HMGA2, TNC, ARG1 and uPAR—four markers of epithelial and TME subpopulations enriched in the progenitor niche (**Figures 3F** and **S3E,F**). In addition, PanCK staining allows identification of epithelial cells, and GFP positivity serves as a proxy for *p53-*proficient premalignant cells. We collected three samples from KP^LOH^ premalignant tissue 2 days post-pancreatitis, and compared it to tumors (*n* = 2) and microtumor (*n* = 1) that emerged spontaneously in the same model.

#### Segmentation

We used Cellpose-SAM (v4.0.7) with the following parameters: *batch_size* = 64, *flow_threshold* = 0.4, *cellprob_threshold* = 0, *normalize* = {“tile_norm_blocksize”: 0} to obtain a nuclear and a cytoplasmic segmentation mask. In the case of tumor samples, we manually identified regions of interest using QuPath in an effort to reduce the size of input images to Cellpose. Collectively, these regions of interest encompassed the entire tissue sample. As inputs, we used DAPI (nucleus) and PanCK (cytoplasmic) images preprocessed by clipping the pixel intensity distribution at the 10th and 99th quantiles, followed by minmax normalization. We consolidated nuclear and cytoplasmic masks by first creating a binary mask representing pixels in nuclear regions. We multiplied this binary image and our cytoplasmic mask, effectively creating a new nuclear mask that shared labels with the cytoplasmic one. Following removal of objects with area < 20 px, we identified and removed cells in the cytoplasmic mask that were not associated with a nucleus. This resulted in a pair of nuclear and cytoplasmic segmentation images with harmonized labels.

#### Quantification

We used a rolling ball background subtraction approach prior to quantification, with the radius *r* empirically determined on a per marker based on their staining pattern: e.g., we used *r* = 10 pixels for HMGA2, that shows pan-nuclear stain, and decreased it to *r* = 5 for uPAR and TNC, which localize to the membrane stain. A smaller radius tends to enhance the signal of smaller objects, but can subtract true signal from larger ones. Following background subtraction, we used the regionprops function from the skimage.measure module to extract the mean intensity of each protein marker in either the nuclear or cytoplasmic masks.

#### Filtering

We filtered cells with nuclear areas outside the range of 20–1000 px in premalignant samples, and 20–1500 px in malignant samples. The larger nuclear area threshold for malignant samples was motivated by the inspection of nuclear size distributions, which were consistent with nuclear enlargement during cancer development. In addition, we removed objects with solidity < 0.85 in premalignant samples, and solidity < 0.8 in malignant samples, again motivated by the decrease in morphological regularity that can emerge during cancer development. Lastly, we used triangle thresholding to remove the lower tail of the log transformed DAPI intensity distribution in order to remove potentially spurious nuclei.

#### Identification of epithelial and HMGA2+ cells

We determined thresholds for positivity of GFP and HMGA2 on a per-sample basis. We used otsu thresholding on the log-transformed cytoplasmic GFP intensity distribution to identify GFP positive (*p53*-proficient) cells. Next, we used triangle thresholding on the linear HMGA2 intensity distribution or GFP positive cells to identify HMGA2+ cells. We decided to use the linear scale of the distribution as the thresholds identified through this method better matched our visual assessment of HMGA2 positivity, as opposed to the ones identified in the log-transformed intensity distribution.

#### Comparison between premalignant and malignant niches

We aimed to quantify how the intensity of HMGA2, TNC, ARG1 and uPAR differed in the vicinity of HMGA2+ cells, as compared to their HMGA2 negative counterparts, as a function of tumor progression (premalignant and PDAC) (**Figure S2K**). For each premalignant sample, we identified all GFP+/HMGA2+ cells, and randomly sampled an equal number of GFP+/HMGA2-cells. In the case of malignant samples, we identified 5000 GFP-/HMGA2+ (PDAC) cells, and 5000 GFP+/HMGA2+ (premalignant) cells. Next, we identified the set of cells with a nuclear centroid within 60 μm of each of these anchor centroids, computed the average intensity of each marker across cells in each neighborhood, and averaged across neighborhoods centered on either HMGA2+ or HMGA2-cells. We normalized intensities in HMGA2+ neighborhoods by the average intensity in GFP+/HMGA2-neighborhoods on a per-sample basis to control internally for differences in intensity emerging during tissue fixation/preparation.

#### Quantification of expression as a function of distance from HMGA2+ cells

To quantify how the expression of protein markers change as a function of distance from HMGA2+ or HMGA2-cells, we adopted the same strategy described above (<u>Comparison</u> <u>between premalignant and malignant niches</u>), except that we computed neighborhoods up to a 200 μm radius, and kept information regarding the distance of each neighbor cell to the anchor centroid (**Figure S2L**). This allowed us to quantify the average expression of each marker as a function of distance from a given cell. Next, we averaged these distance-intensity functions across all cells from each premalignant sample.

### smFISH probe design and data quantification

#### Probe design

We built upon published software^117^ to design custom panels for multiplexed smFISH. This design strategy relies on pre-computation of all possible 30mer sequences found in mouse cDNAs (Ensembl GRCm38.p6), augmented with coding sequences of fluorescent proteins engineered into our mouse model. We excluded pseudogenes from the potential pool of mRNAs used for probe design. We compute multiple scores for each 30mer, including Tm, GC content, and potential for hybridization with rRNAs and tRNAs. We used the following parameters to include a 30mer into our candidate probe-set: GC-content (43–63%), Tm (66–76°C), excluding 30mers that contain at least a 15mer present in an rRNA or tRNA.

In addition, we computed expression-informed penalties to estimate the specificity of each candidate probe. We adapted published software to include single-cell information into the estimation of specificity scores, reasoning that it would decrease the chances of selecting probes with off-target binding to highly-expressed genes in rare cell populations. To do so, we considered our single-cell data from epithelial and immune compartments of the injured pancreas. Furthermore, we leveraged a published single-cell time course of *Kras*-driven transformation in pancreatic tissue to incorporate information about fibroblast, pericyte and endothelial gene expression^29^. Since our spatial analysis is focused on the premalignant-stage of pancreatic tumorigenesis, we excluded cancer-associated samples for the purpose of computing specificity scores.

To summarize single-cell gene expression as a function of cell state in distinct cellular compartments (epithelial, immune, fibroblast, pericyte and endothelial), we aggregated cells into metacells, representing discrete fine-grained cell states, using SEACells v0.2.0^108^. For each cellular compartment, we first used standard log library-size normalization and dimensionality reduction using PCA (*n_pcs* = 100) for preprocessing. We then ran SEACells using *n_waypoint_eigs* = 10 (default) and *waypoint_proportion* = 0.9 (default). We selected the number of metacells per compartment such that the median number of cells per metacell ranged within a similar range:

● Epithelial: 300 metacells, median size = 79 individual cells
● Immune: 150 metacells, median size = 69.5 individual cells
● Fibroblasts: 100 metacells, median size = 88 individual cells
● Pericytes: 15 metacells, median size = 63 individual cells
● Endothelial: 50 metacells, median size = 85.5 individual cells

Next, we computed a summarized gene expression matrix *X* of dimensions *n* x *m*, where *n* is the number of metacells across all cellular compartments, and *m* is the number of genes in the dataset. In this matrix, *x*_ij_ is the average normalized linear counts of gene *j* across individual cells in metacell *i*. We normalized *X* by total counts per metacell, and scaled by an arbitrary factor of 2000. Lastly, we identified the maximum expression per gene across all metacells, and used these to compute specificity penalties during probe design. This strategy penalizes off-target binding to highly expressed genes, even when such high expression occurs in rare cell subpopulations.

We computed transcription-wide specificity as published^117^, except that we assumed all isoforms of a gene contribute uniformly to its total expression (our single-cell data lack isoform-specific information). To compute the specificity score, each 30mer was represented as a collection of overlapping 17mer sequences in sliding windows with 1-bp shift. For each 17mer, we calculated the fraction of occurrences in the on-target gene (any isoform) out of all occurrences in the transcriptome, weighted by gene expression to penalize off-target binding to highly-expressed genes. We computed a final specificity score ranging from 0 (no occurrence from on-target gene) to 1 (all occurrences from on-target gene) for each 30mer by averaging the scores of its constituent 17mers, and selected 30mers with scores above 0.75 as candidates for our panels.

Candidate 30mers were used to compile primary probes for a set of query genes. We selected Ensembl canonical isoforms to design probes targeting a particular gene, aiming for 92 non-overlapping probes per gene. Whenever this was not possible due to transcript length, homology to other genes, or other sequence properties, we allowed a maximum overlap of 20 bp between probes. The use of overlapping probes was previously reported to maximize smFISH signal^153^ due to the probabilistic nature of probe–mRNA binding. Lastly, we appended readout sequences to each probe, which serve as recognition sequences for fluorescently labeled readout probes. In the case of genes for which we were not able to generate at least 75 probes, we added two or four copies of the selected readout sequence in order to amplify the fluorescent signal coming from such probes. Sequences of probes used in this study are included in **Data S7**.

#### Quantification of Msn+ cells with smFISH

Iterative smFISH imaging is subject to subtle drift in the x-y plane due to the physical interaction between the 60x oil objective and the plateholder. To correct for such drift, we used rigid body registration on maximum intensity projection of DAPI images corresponding to distinct imaging cycles. Since drift was more pronounced in the first imaging rounds and then stabilized, we manually identified a reference imaging cycle that minimized overall signal loss after alignment. For each mouse, we analyzed five fields of view, randomly chosen at the time of data acquisition, blinding their selection to the presence of *Msn* signal. We quantified the prevalence of *Msn*+ cells and the size of their spatial clusters by manually identifying epithelial nuclei (*Gfp*+ cells) with ImageJ, generating binary masks where each dot corresponds to an individual cell centroid. In addition, we manually traced *Msn*+ epithelial regions in the image to allow computational classification of epithelial centroids as *Msn*+ or *Msn*-.

To quantify the fraction of *Msn*+ cells in each tissue sample, we identified individual cell centroids locating local maxima in distance-transformed binary mask images. Next, we used the shapely (v2.1.2) and geopandas (v1.1.1) libraries to convert *Msn*+ region masks into polygons, with which we intersected epithelial centroids. **Figure 1I** shows the fraction of *Msn*+ epithelial cells across all FOVs in each biological replicate (*n* = 4).

To quantify the size of Msn+ epithelial clusters in smFISH data, we first determined the average distance between adjacent epithelial cells, by computing a kNN graph on centroid coordinates (*k* = 3). The median distance between adjacent cells was 47.5 pixels. Next, we computed a spatial neighbor graph considering only *Msn*+ centroids, connecting cells within a 47.5 pixel radius.

Next, we identified connected components in this spatial neighbor graph. **Figure 1J** shows the distribution of the size of connected components that each Msn+ cell in our dataset belongs to, quantified for each biological replicate (*n* = 4).

## SPATIAL TRANSCRIPTOMICS DATA: PANEL DESIGN AND DATA ANALYSIS

### Custom probe set design for spatial transcriptomics

Spatially resolved transcriptomics provides the opportunity to localize the rich transcriptional heterogeneity uncovered by single-cell experiments in tissue context. In the premalignant pancreatic epithelium, this includes transcriptional gradients that connect cellular states and key signaling axes such as *Kras* and *p53*. The premalignant epithelium undergoes dramatic microenvironmental remodeling, including the formation of fibrotic and inflammatory niches rich in myeloid cells. We sought to understand (1) the interplay between the adoption of distinct pancreatic epithelial states and changes in the microenvironment, (2) the spatiotemporal dynamics of niche transitions, and (3) what intercellular signaling circuits may mediate the formation and stabilization of progenitor niches. Thus, we designed a custom Xenium probe panel targeting 480 genes (**Data S2**) to capture:

1. Transcriptional heterogeneity in major cellular compartments, including cluster-level heterogeneity and transcriptional gradients connecting disparate cellular states.
2. Activation of major signaling programs (e.g., *p53*, *Kras*, *Yap*, interferon) operating in the benign-to-malignant transition in pancreatic cancer.
3. Genes that may construct intercellular circuits through juxtacrine and paracrine interactions.

#### Reference single-cell datasets for spatial expression profiling

Marker selection for imaging-based spatial transcriptomics should consider gene expression estimates for all cell populations in the target tissue. Optical crowding from abundant mRNA species not only hampers their accurate quantification, but also that of other mRNAs in the vicinity, due to the combinatorial encoding of mRNA identity through sequential imaging^116^. To inform our custom Xenium probe set, we leveraged the premalignant pancreas expression atlas we compiled for smFISH probe design (see **Probe design for multiplexed smFISH**), which includes epithelial and immune cell profiles in response to injury (with or without *p53* perturbation) as well as fibroblast, endothelial and mural cell data (excluding tumor-stage samples to better match premalignant cell stated distributions) from Schlesinger, Yosefov-Levi, Kolodkin-Gal *et al*.^29^.

Our compiled dataset also leveraged SEACells to generate more robust estimates of gene expression distributions across cell states, including rare cell states^108^. We reasoned that information from rare cell states would help us to retain biologically important mRNAs expressed in a small fraction of cells, and to exclude abundant mRNAs prone to optical crowding in rare cells and adjacent cells. To this end, we used the maximum expression per gene across all SEACells to guide marker selection for imaging-based spatial transcriptomics.

#### Gene expression constraints on marker selection

We avoided selecting mRNAs that lacked discrete foci due to optical crowding in smFISH staining of the premalignant pancreas (**Data S1J,K**; e.g. *Cpa1* in acinar cells, *Muc6* in gastric chief-like premalignant cells, *Tff1* in gastric pit-like premalignant cells or *Acta2* in activated stroma, *Fn1* in progenitor-like cells). Moreover, we identified ideal expression thresholds based on genes that we successfully imaged using multiplex smFISH (e.g. *Ptprc* in immune cells, *Adgre1* in macrophages, *Msn* in progenitor-like cells, *Anxa10* in gastric pit-like cells). Guided by experience with individual markers, we set the lower bound (max expression across metacells > 0.5), upper bound for subpopulation markers identified through unbiased clustering (max < 2.5), upper bound for gene-set derived markers such as communication genes (max < 4) and upper bound for biologically curated markers (max < 10), though we did make exceptions for certain markers of biological interest (e.g., expression of type I interferons, *Pecam1* endothelial marker).

#### Marker selection for cellular compartments and microenvironmental subpopulations

We used a combination of prior knowledge and unbiased marker gene identification, guided by expression constraints as described above. For example, we selected well-recognized markers of epithelial (*Cdh1*), immune (*Ptprc*), macrophage (*Adgre1*, *Csf1r*), fibroblast (*Pdgfra*, *Col5a1, Vim*), endothelial (*Pecam1*) and pericyte (*Des*, *Pdgfrb*) cells. Although excluded from our premalignant atlas, we included markers for a rare subpopulation of glial cells (*Fabp7*, *Plp1*) identified by Schlesinger, Yosefov-Levi, Kolodkin-Gal, *et al*.^29^, as well as *Fabp4*, an adipocyte marker.

To dissect heterogeneity within cellular compartments, we computed differential gene expression to identify markers of manually curated cell subpopulations (immune dataset^7^) or PhenoGraph clusters (*k* = 30) (fibroblast, endothelial and mural cells). We prioritized known markers of subpopulations (e.g. *Tnc*, *Dpt* or *Gli1* in fibroblasts) or communication pathways (e.g., *Csf1* in Gr-MDCs, *Lifr* in fibroblasts, *Kitl* in endothelial cells or *Csf2rb* in mural cells), as well as genes that independently mark subpopulations in distinct compartments (e.g. *Prox1* as a marker of normal duct cells and lymphatic endothelial cells). Collectively, this marker selection strategy allowed us to identify subpopulations across multiple compartments, while maintaining the biological interpretability of gene expression patterns in our spatial data.

#### Marker selection for premalignant epithelial cells

We aimed to capture the major premalignant states that we identified in dissociated single-cell data, including the cluster of progenitor-like cells with mesenchymal phenotypes that accumulate upon *p53* loss. We started with markers that have been reproducibly identified in independent single-cell datasets, including acinar (*Rbpj1*, *Nr5a2*), tuft cell (*Pou2f3*, *Dclk1*), neuroendocrine (e.g. *Syp*, *Ascl1*), gastric-like (*Elf3*, *Onecut2*)^29^, and progenitor-like (e.g., *Msn, Hmga2, Nes*) cell-state markers. We next used differential expression in PhenoGraph clusters to select markers that capture heterogeneity in gastric and progenitor-like states; for example, distinct subpopulations of gastric-like cells that correspond to chief-like (*F5*) or pit-like (*Anxa10*) states. In addition, we included *Cp* and *Rgs5*, two genes expressed in premalignant cells most resembling ducts. Lastly, we included a number of genes that transiently increased at the boundary of gastric-like and progenitor-like states (e.g. *Msln*, *F3*), as well as genes that progressively increased in expression along the progenitor axis (e.g. *Zeb2*, *Grem1* and *Piezo2*).

We made the conservative decision not to probe for mRNAs of fluorescent proteins to distinguish premalignant and normal cells in our samples, due to the potential of these constitutively expressed mRNAs to hamper decoding the rest of the panel. Despite this, we note that we were able to identify structural and molecular features corresponding to normal ducts and islets, allowing us to focus on lesions consistent with premalignant phenotypes.

#### Marker selection for signaling pathways and biological processes

We manually selected markers for major signaling pathways, prioritizing negative feedback genes known to constitute some of the earliest transcriptional responses to pathway activation. Specifically, we probed for *p53* (e.g., *Mdm2, Cdkn1a, Bax*), MAPK (e.g., *Dusp4, Dusp6, Dusp5, Spry1, Spry2*), interferon (e.g., *Socs1, Socs3, Irf7, Oasl1*), YAP (*Ccn1, Ccn2*) and TGF-β (e.g., *Smad7, Il11, Has2*) signaling. In addition, we probed for cell cycle regulators, including major cyclins and cyclin-dependent kinases (CDKs) (e.g., *Ccnd1, Ccne1, Ccna1, Cdk1, Cdk2, Cdk4, Cdk6*) as well as CDK inhibitors (e.g., *Cdkn1b, Cdkn1c, Cdkn2a, Cdkn2b*).

#### Selection of genes involved in cell–cell interactions

To prioritize genes involved in cell–cell communication, we leveraged our previously identified communication modules—groups of ligands and receptors that are selectively upregulated upon *Kras* signaling in the premalignant pancreas and are predicted to form multiple interactions between cell states^7^. These communication modules involve a plethora of cytokines and receptors with known and predicted roles in tissue remodeling during tumor initiation (e.g., *Il33, Il18, Il1a, Ccl2, Csf2, Lif, Vegfa, Fgf*) and their cognate receptors. In addition, we probed for ligands and receptors from *Wnt*, *Shh*, *Bmp* and *Notch* signaling, due to their roles in development and tissue repair. Lastly, we included genes involved in cell adhesion (e.g., claudins, cadherins, integrins) and juxtacrine signaling (e.g., ephrin and semaphorin signaling), prioritizing genes based on gene expression constraints and HVG status (*n* = 3000) within each cellular compartment.

#### Computational design of probe set

We used 10x Xenium Designer (10x Genomics) with vendor assistance to generate isoform-specific probes for *Cdkn2a*. We used three independent reference datasets to compute cell-type utilization scores, which inform the potential for optical crowding as a function of target mRNA abundance in specific cellular states. Specifically, we used in-house single-cell data for premalignant epithelial cells after injury with or without *p53* knockdown (this work), immune cells in the injured premalignant pancreas^7^ and a PDAC progression atlas that includes all cellular compartments^29^. Given that these datasets were generated using different Ensembl annotation versions, we mapped *gene_symbols* to *ensembl_ids* to match those of the Ensembl build Mus_musculus.GRCm38p6.102. We excluded any gene above the recommended cell utilization score recommended by 10x Xenium Designer.

### Spatial transcriptomics embedding and annotation

In our spatial data, each cell is represented by three primary features: (1) a transcript count matrix, (2) an x-y spatial coordinate (in μm), and (3) a polygon describing its segmented nucleus. In addition, we derive secondary features describing the morphological and molecular properties of the local niche of a cell, such as the structure and size of premalignant lesions, and the average gene expression of a given cell type in the vicinity of a given cell. These multiple viewpoints allow us to identify molecular and morphological correlates between the intrinsic state of a cell and its microenvironment. Our data is composed of 33 tissue samples, spanning 9 conditions, including different collection timepoints in the response to acute pancreatitis, as well as pharmacological inhibition of oncogenic KRAS, and *p53* knockdown in premalignant cells. In total, this comprises 9,463,399 cells (excluding cells with mixed phenotype), with an average of 92.83 mRNA counts per cell, and 57.92 detected genes in each cell.

In this section, we first describe our strategy to compute cell-level and niche-level representations from single-cell-resolved spatial transcriptomics data. Then we describe how we integrate these different viewpoints to study the dynamic interplay between premalignant epithelial cells and their microenvironment.

#### Transcript-to-cell assignments

Assigning transcripts to cells is a critical first step in recovering comprehensive and accurate cell states from imaging-based spatial transcriptomics data. Because cell boundary estimates based on pure geometric constraints, such as nuclear segmentation expansion followed by Voronoid tessellation (default 10x Xenium processing), result in pervasive transcript cross-contamination from adjacent cells, we opted to construct a count table composed only of transcripts that overlap nucleus masks (10x nucleus transcripts). This approach successfully eliminated inter-cell transcript contamination, as evidenced by our recovery of Leiden clusters with reasonable cell type purity based on marker expression (**Data S1.2L-P**). However, it also resulted in an average loss of 57% of transcripts per slide (interquartile range 53%–60%). Although the limited sensitivity of this conservative approach is not ideal for all contexts, it was sufficient to recover expected cell-state heterogeneity in every cellular compartment (e.g., myofibroblast, myeloid, lymphoid and endothelial subpopulations) in our data (**Data S1.2L-P**), including transcriptional gradients, such as the gastric–progenitor continuum in the premalignant epithelium (**Figure 3E,F**). This is likely due to our use of probes targeting highly expressed marker genes in our panel, and to the fact that we focused on cell types that were not exceptionally small and therefore contained relatively high transcript counts.

#### Gene filtering, cell filtering and normalization

We excluded negative control probes from library size estimation and dimensionality reduction steps, but left them in the data for possible use in subsequent processing steps. Furthermore, we excluded cells with fewer than 25 mRNA counts. On average, we excluded 9.5% of cells per slide (interquartile range 7–10%).

For normalization, we estimated size factors per cell as the total 10x nucleus counts per cell. We normalized count matrices by dividing by size factors, and scaling by the median size factor in the data. We used linear, as opposed to log transformed counts, as this led to faster and reproducible convergence during the computation of UMAP embeddings. Linear counts may be more appropriate for this data modality given that our panel is enriched in communication genes, signaling proteins and transcriptional regulators, gene classes that tend to have lower expression than the abundant house-keeping genes in transcriptome-wide datasets.

#### Single-cell embeddings for spatial transcriptomics

We computed an initial single-cell embedding using concatenated count matrices from all tissue slices. To compute this embedding, we used GPU-based implementations of PCA, kNN graph construction and UMAP functions in rapidsai (v24.2.0). We conducted dimensionality reduction using PCA on linear normalized counts, keeping the top 136 PCs that captured 75% of variance. In addition, we noted that the use of more than 100 PCs was important to isolate molecularly and spatially distinct subpopulations that were nonetheless rare in the dataset (i.e., adipose, glial and non-pancreatic epithelial cells). Use of a smaller number of PCs collapsed these subpopulations into stromal and epithelial compartments, hindering the purity of our initial cell type annotation. We constructed a kNN graph using the GPU-based nearest-neighbor graph implementation of cuML python package (v24.02.00). We computed the kNN graph using euclidean distance on PC space (*k* = 30) as our metric, followed by UMAP for visualization (*n_neighbors* = 10), and computation of clusters using GPU implementations of leiden (*resolution* = 1.0). We used this integrated embedding to annotate coarse and refined cell states (see below). These annotations were transferred into any condition-specific embedding to maintain consistency in downstream analyses.

#### Major compartment annotation

We annotated major cellular compartments before determining more refined cell states (details below). We leveraged spatial patterning to inform cell type annotation—for example, by identifying cellular states associated with lymph nodes, the edge of the tissue (e.g., mesothelial) or the pancreatic parenchyma.

To identify major cellular compartments, we annotated clusters based on known markers, then refined cell states within cell types by re-embedding and clustering cellular subsets. Specifically, we first computed clusters using the Leiden algorithm on a kNN graph (*k* = 30) constructed on PC space, with *resolution* = 1.0. We manually assigned clusters to major compartments based on markers in our panel: epithelial (*Cdh1, Itgb4, Cpa1, Onecut2, Prox1, Cp, Krt7*), endothelial (*Pecam1*), immune myeloid (*Ptprc, Adgre1, Csf1r, Csf3r, Cd68, Itgam, Itgax*), immune lymphoid (*Ptprc, Cd3g, Cd4, Cd8a, Foxp3, Cd19, Cd79a, Lman1, Gata3, Rora, Trdc*), fibroblast (*Col5a1, Pdgfra, Dpt, Pdpn*), mural cell (*Des, Rgs5, Tagln, Fhl1*), adipose (*Fabp4*), mesothelial (*Msln*) and glial/nerve (*Plp1, Fabp7, Ncam1, Ncam2, Syp, Ngfr*).

### Refined cell-state annotation and condition-specific embeddings

To identify more granular cell states, we re-embedded cells and recomputed clusters within each compartment, then annotated clusters based on the expression of marker genes from the literature, manually curated single-cell atlases of the premalignant pancreas, or unbiased identification of cluster marker genes using scanpy’s sc.tl.rank_genes_groups. The latter approach highlighted genes with higher expression in a cluster relative to all other clusters; among these, we selected markers that would provide the most interpretable cell-state label (e.g., *Cd55* in endothelial cells or *Ccn2* in myofibroblasts). In addition, we computed condition-and subpopulation-specific embeddings for downstream analyses. For example, **Figures 2,3** and **5** focus on variation in epithelial cells and their niches in unperturbed samples to gain insights into tissue remodeling around progenitor-like cells.

To compute refined single-cell embeddings, we first excluded negative control probes and genes expressed in fewer than 1% of cells within a compartment, so that they would not influence normalization and dimensionality reduction. Next, we normalized the data by dividing counts by library size in each cell and re-scaling by the median library size, applied PCA on normalized linear counts, and kept the top PCs that captured 75% of total variance. We constructed a kNN graph on Euclidean distance in PC space (*k* = 30) and visualized our embeddings using UMAP (*n_neighbors* = 10, *min_dist* = 0.1).

Next, we computed clusters at different resolutions to dissect heterogeneity within cell types. We used Leiden clustering on the kNN graph with *resolution* = 1.0 for initial cluster annotation, then computed PhenoGraph clusters on the kNN graph by leveraging GPU implementations of Jaccard similarity matrix construction and Louvain clustering at multiple resolutions (*resolution* = 1.0, 0.75, 0.5) in the cugraph module of rapidsai. The use of multiple resolutions allowed us to distinguish closely related cell states during cell type refinement, while avoiding noise from over-clustering.

See **Data S10** for all cell-type or condition-specific embeddings in our study; methods for annotating specific cell states are presented below.

#### Epithelial state annotation

We first annotated Leiden clusters based on major premalignant subpopulation markers identified during PDAC initiation (^7,29^ and this work): progenitor1 (*Msn, Hmga2, Itgb4*), progenitor2 (*Vim, Piezo2, Tnc*), gastric-like (*Elf3, Onecut2, Lgals4*), gastric chief-like (*F5*), gastric pit-like (*Anxa10*), tuft (*Pou2f3, Dclk1*), neuroendocrine (*Syp, Hepacam2*), duct-like (*Cp, Prox1, Rgs5*), ADM (*Rbpjl*), and cycling cells (Mki67).

To distinguish normal ducts from premalignant cells with a duct-like phenotype, we re-embedded the duct subpopulation using the above strategy. We identified clusters corresponding to normal ducts based on *Rgs5* and *Prox1* expression^29^. Cells in this cluster formed morphological structures characteristic of normal duct, providing support for our annotation.

#### Fibroblast state annotation

We used fibroblast signatures from cancer contexts^53,154^, together with the spatial organization of stroma in the premalignant pancreas, to assign Leiden clusters:

1. myCAFs. Most fibroblasts in the premalignant pancreas (86%) resembled myofibroblast cancer-associated fibroblast (myCAF) states (*Sdc1, Tagln, Tgfb1, Tnc, Epha4, Igf2, Itgbl1*). These cells populated the pancreatic parenchyma, and were in close proximity to premalignant epithelial cells (**DataS1L**). We identified marker genes expressed in myofibroblast subclusters using the scanpy function sc.tl.rank_genes_groups, and used these markers to label specific subpopulations (e.g., *Tnc+, Ccn2+, Gli1+, Igf2+*).
2. iCAFs. A group of cells were consistent with the inflammatory cancer associated fibroblast (iCAF) phenotype (*Cxcl1, Cxcl12, Tnxb*) and expressed the universal fibroblast marker *Dpt*^155^. These fibroblasts were excluded from the parenchyma (**DataS1L**).
3. Trf+ CAFs. A small subset of fibroblasts (0.2%) expressed antigen-presenting CAF (apCAF) markers (*Trf, Sdc4*). We conservatively labeled these as Fibroblast_Trf given that our panel did not include additional apCAF markers.

#### Myeloid state annotation

We used our single-cell immune atlas of the premalignant pancreas^7^ to group myeloid cells into three broad categories, followed by cell-type refinement (**DataS1M**):

1. Monocyte/macrophages. Most myeloid cells in the premalignant pancreas (91%) resembled monocyte/macrophages (*Csf1r* and *Adgre1*). Spatially patterned clusters within this compartment guided the annotation of refined states. Specifically, *Cd163*+ cells localized outside the pancreatic parenchyma and *Nlrc5*+ cells localized to lymph nodes; *Maf* and *Itgax* characterized clusters at two extremes of the continuum of macrophage states; and a cluster of *Itgax*+, *Cd274* (PD-L1)^high^ cells appeared upon *p53* knockdown in the premalignant epithelium. Thus, granular myeloid subsets were characterized by spatial patterning in addition to transcriptional variation in the premalignant pancreas.
2. Granulocyte–myeloid derived cells (GrMDCs). A cluster defined by high *Csf3r* and low *Csf1r* expression characterized GrMDCs.
3. Dendritic cells. Dendritic cells were defined by *Itgax* (CD11c) expression, lack of *Csf1r* and *Adgre1*, and expression of dendritic cell markers in the panel (*Ifi205, Itgae, Jak2*).

#### Lymphoid state annotation

‘We used markers from our single-cell immune atlas of the premalignant pancreas^7^, along with cluster refinement and marker identification using the scanpy sc.tl.rank_genes_groups function to define lymphoid cells groups. We re-embedded non-B-cell-related lymphoid cells (see <u>Refined cellular states and condition-specific embeddings</u>), followed by PhenoGraph clustering and cell-state annotation to define the following populations: (1) B cells (*Cd79a, Cd19*) and plasma cells (*Lman1*), (2) parenchyma associated lymphoid cells, including Tregs (*Cd4, Foxp3*), Th17 cells (*Il23r*), gdT cells (*Trdc*), innate lymphoid cells (*Gata3*), NK cells (*Ncr1*) and mast cells (*Kit*), and (3) lymph-node-associated CD4 and CD8 T cells (**DataS1N**).

#### Endothelial and mural state annotation

Endothelial cells were broadly divided into vascular (*Pecam1+/Prox1-*) or lymphatic (*Pecam1+/Prox1+*) subpopulations. Within the vascular subpopulation, we labeled clusters based on known biology (e.g., activated endothelial cells expressing *Selp*) or marker expression (e.g., *Piezo2*+ endothelial cells). We annotated mural cell clusters based on markers identified using the scanpy sc.tl.rank_genes_groups function. Endothelial and mural cells exhibited their expected spatial colocalization, and clusters of these cellular compartments also showed spatial patterning related to vessel size (**DataS1O,P**).

#### Mixed cell-state annotation

Some clusters showed evidence of cross-contamination between cell types, particularly between fibroblasts and myeloid cells (e.g., expression of macrophage marker *Csf1r* with fibroblast-specific *Igf2*). These artifacts can emerge from errors in nuclear segmentation or from 2D projection of transcripts, both of which are expected to increase in tightly packed tissues such as the premalignant pancreas. We flagged clusters in these mixed states, kept them in the dataset for visualization of tissue architecture, but excluded them from biological analyses involving cell type proportions or changes in gene expression between niches. Altogether, mixed cell states encompassed 3.1% of our data, and were dominated by cells sharing fibroblast and myeloid profiles (1.7% of all cells). The following list summarizes the absolute number and fraction of cells in our dataset corresponding to mixed states:

● mixed_myeloid_stroma: 166,979 cells (1.7%)
● mixed_epithelial_stroma: 70,906 cells (0.72%)
● mixed_endothelial_immue_stroma: 33,375 (0.34%)
● mixed_lymphoid_other: 22,790 cells (0.23%)
● mixed_mural_fibroblast: 8,310 (0.08%)
● mixed_lymphoid_myeloid: 3,338 (0.03%)
● mixed_mural_immune: 2,750 (0.02%)
● mixed_mural_other: 1,233 (0.012%)

#### Compartment-aware gene censoring

Despite restricting our analyses to transcripts called by 10x Genomics nucleus segmentation and excluding clusters with mixed cell states, we observed some transcript contamination between cell types. For example, *Igf2* was expressed in myeloid cells, despite its absence in dissociated reference datasets^7^. As expected, these effects were larger for highly expressed markers of specific subpopulations or cellular compartments, which can result from projecting expression onto two dimensions from a 3D section (https://doi.org/10.1101/2025.03.14.643160, https://doi.org/10.1101/2025.01.20.634005), or from transcript diffusion.

To mitigate the effect of contaminants in downstream analyses, we leveraged dissociated reference datasets^7,29^ to identify the fraction of cells within each cellular compartment with non-zero counts for any given gene. For downstream analyses, we censored genes in each cellular compartment that were expressed in < 1% of cells in every reference dataset. This threshold was meant to exclude genes with little evidence for expression in single-cell data, suggesting that their presence in our spatial dataset could be due to cross contamination between cell types. On average, 303 (67%) of the genes included in our panel passed this filter.

To complement filtering based on summary statistics from dissociated cells, we identified robustly expressed genes based on the fraction of cells within a subpopulation of a cellular compartment that express a given gene (gene detection rate) in our spatial data. Assessing per-subpopulation rather than per-compartment statistics ensures that no positive populations are missed.

For a specific cell type, this resulted in a matrix *P*^cxm^ where *c* is the number of subpopulations within the cell type, and *m* is the number of genes. To identify a ‘robust expression’ threshold within a cell type, we flattened the matrix *P*, and computed the histogram of subpopulation-level gene detection rate (*n_bins* = 25). Next, we used triangle thresholding to identify a cutoff that distinguished genes with high or low detection rates. On average, this strategy nominated genes as robustly expressed if they were detected in at least 9% of cells within a compartment subpopulation. We applied it to all coarse cell types in our data to identify compartment-specific gene sets with evidence of robust expression. Across cell types, an average of 262 (54%) of genes were classified as robustly expressed. For a given cell type, an average of 240 (50%) of genes passed both the spatial and dissociated data detection thresholds. We used this high-confidence gene set to analyze within-cell-type spatial heterogeneity in expression (**Figures 5A,B, S4E** and **S5A–D**). We note that for future work, newer methods can correct transcript misassignment resulting from contamination in spatial transcriptomics data (https://doi.org/10.1101/2025.01.20.634005).

**Data S10** summarizes the results of compartment-aware gene censoring across all cellular compartments.

### Gene expression and cell-state gradients from spatial transcriptomics data

Our single-cell characterization revealed that premalignant epithelial phenotypes do not exist as discrete states. Rather, we and others observe pervasive continuity in the transcriptional heterogeneity of individual premalignant cells (**Figure 1E**)^7,29^. These continuums are reflected by the presence of cells with mixed states (**Figure 1E**), which suggest cellular plasticity.

Embracing continuity in cell states allowed us to connect these features of plasticity at the single-cell level with tissue remodeling events encoded in spatial transcriptomics data. We took two complementary approaches to characterize transcriptional gradients in premalignant cells: (1) we used gene signatures of canonical premalignant subpopulations^7^ to visualize their continuous variation along an epithelial-specific phenotypic manifold and (2) we used diffusion component analysis to identify the major axes of transcriptional variation in epithelial states.

#### Visualization of gene expression signatures in epithelial cells

We asked whether epithelial cells in our spatial data recapitulated the spectrum of premalignant transcriptional states we observed in dissociated scRNA-seq. We used markers of premalignant subpopulations in our Xenium panel as transcriptional signatures (**Data S2**), and computed a signature score matrix X^nxz^ where *n* is the number of epithelial cells in non-perturbed samples (*n* = 1,388,199) and *z* is the number of epithelial states (*z* = 6, progenitor-like, gastric-like, duct-like, neuroendocrine, tuft, adm). To compute the signature score matrix, we first standardized our log-transformed count matrices (*pseudocount* = 1) by computing the *z*-score of each gene over all epithelial cells. The signature score *x*_i,j_ is the average standardized expression of genes in *signature*_j_ for *cell_i_*. We normalized the signature score matrix column-wise, such that signature scores ranged between [0,1], saturating the signature score at the 90th quantile. We visualized signatures by plotting epithelial cells in both spatial and transcriptional (UMAP) domains, pseudocoloring each cell by aggregating RGB color vectors weighted by each signature score (see <u>Simultaneous visualization of multiple signatures</u>). This visualization strategy showed that our spatial data captured the transcriptional heterogeneity in epithelial states in single-cell data (compare to Figs. 1e, 3e, 4b in ref. ^7^). Furthermore, the mixture of signatures at intermediate points connecting extreme cellular states highlighted continuity between subpopulations of premalignant states, such as the presence of epithelial cells with mixed gastric and progenitor signatures.

#### Diffusion component analysis in spatial data

To identify the major axes of continuous variation in premalignant states, we used diffusion maps, which model gradual cell-state transitions along the phenotypic manifold as a diffusion process in a kNN graph. Starting from a kNN graph (*k* = 30) constructed on a single-cell embedding for gastric-like or progenitor-like cells from unperturbed samples (see *Refined cellular states and condition-specific embeddings*), we constructed a diffusion operator, followed by eigendecomposition to identify diffusion components (**see Construction of the diffusion operator and diffusion maps**). The first diffusion component, representing the dominant axis of transcriptional variation among gastric-like and progenitor-like cells, captured gradual changes in gene expression linking gastric-like epithelial states on one extreme, and progenitor-like on the other, including mixed phenotypes at intermediate points (**Figure S2E**). This approach positioned each of the 838,581 gastric-like or progenitor-like cells in our data along a continuous axis of variation between these subpopulations.

### Spatial transcriptomics niche analyses

#### Definition and identification of cellular niches

Spatial transcriptomics makes it possible to represent a cell not only by its intrinsic properties, but also by the properties of its cellular neighbors and surrounding tissue structure. The spatial niche framework provides a simple and elegant approach to connect a cell with its surroundings^156,157^. In general, a niche can be defined as a set of cells in shared physical space, represented by a spatial neighborhood graph *Gn* × *n*, where *n* is the number of cells in the data, and *G*_i,j_ is a measure of spatial distance between *cell*_i_ and *cell*_j_.

We used 2D Euclidean distance as our metric, and defined a niche as the set of cells within a radius *r* of a reference cell, which we refer to as the anchor cell. In using a fixed radius to define a niche, we can ensure biological length scales that are interpretable and represented by well-defined physical units; dissociated single-cell data, by contrast, does not report physical distance, with implications for our understanding of the spatial extent of cellular influences. We use a radius of 60 μm throughout this work as this length scale captured glandular structures in the premalignant pancreas, establishing individual lesion size as a unit of analysis. However, we recognize different choices of *r* are bound to highlight different emergent properties of intercellular communities, such as the formation of large spatial domains of progenitor-like cells upon *p53* knockdown (**Figure S7C**).

We extracted three types of features from our spatially defined niches: (i) morphological parameters, (ii) cell composition vectors, and (iii) compartment-specific locally averaged niche expression matrices (detailed below). These features enabled us to characterize tissue remodeling events that are coupled to gradual changes in premalignant cell states.

#### Quantification of structural and morphological parameters

To study morphological changes associated with the adoption of distinct premalignant cell identities, we quantified three local tissue parameters in individual niches: lesion size, epithelial fraction, and luminal area. Our computational approach relied on graph and pixel-based representations of epithelial structures, enabling tissue properties to be estimated from cell centroids alone.

To quantify lesion size, we defined a lesion as a contiguous set of physically adjacent epithelial cells in the tissue and built a spatial neighbor graph connecting all epithelial nuclei centroids within 20 μm. The use of this length scale, which is within the range of a cell diameter, maximized the probability that two cells are interacting physically in the tissue. Next, we identified connected graph components in our spatial neighbor graph. We defined epithelial lesion size as the number of cells in each connected component.

#### To quantify epithelial fraction, we computed the fraction of epithelial cells out of all cells in a niche

To quantify luminal area, we adopted classical image processing procedures aimed at identifying holes in the tissue surrounded by epithelial cells. We identified (1) non-empty regions covered by tissue; (2) closed holes within the tissue; and (3) empty space that was not closed and may have resulted from epithelial disruption during tissue processing:

1. To identify empty space, we generated a tissue image starting from single-cell centroids. For each tissue slice, we constructed a 5 μm px^-1^ mesh grid covering all cellular centroids using numpy (v1.23.4) linspace and meshgrid functions. Next, we calculated the number of cell centroids that overlapped each xy coordinate of the mesh grid, resulting in a bitmap-based representation of the tissue, where values correspond to the cell density at each pixel. Using a 2D Gaussian filter with an isotropic kernel of 3-px bandwidth (15 μm), we denoised cell density estimates in the image. Lastly, we applied an empirically determined threshold of 0.01 to identify pixels with high cell density (foreground) from empty pixels (background).
2. To identify closed holes surrounded by epithelial cells, we computed a bitmap-based representation of epithelial cells. Starting from the mesh grid in step 1, we set any pixel overlapping with an epithelial centroid to 1, and all other pixels to 0. To close small gaps between pixels associated with a single glandular epithelial lesion, we dilated our binarized image of the premalignant epithelium using the dilate function in the cv2 package (v4.10.0) with an 10-px disk kernel and *n_iterations* = 1. Next, we filled holes in the image using the cv2 floodFill function, followed by erosion using the erode function to counterbalance the original dilation, returning the boundaries of epithelial lesions to their original scale. Luminal pixels correspond to empty points that lie within filled epithelial masks.
3. To identify empty luminal regions that are not enclosed, we used a pixel propagation procedure, whereby luminal pixels are identified as empty points within 20 μm of an anchor epithelial cell. We used this propagation strategy iteratively, such that luminal points identified in the first iteration lead to the identification of the next layer in the lumen. The use of a 20-μm threshold was empirically determined to prevent classification of the tissue edge as lumen due to the presence of epithelial cells close to the tissue edge.

We defined lumen area as the number of pixels classified as lumen within a radius *r* of a cell (*r* = *60 μm*) for **Figure 2H**.

To quantify how structural features in the vicinity of premalignant cells change along the gastric–progenitor DC, we discretized this DC into 11 equally sized bins, and computed the distribution of niche morphological parameters as a function of an epithelial cell’s DC bin (**Figure 2G–J**).

#### Quantification of cell-state proportions in niches

The distribution of discretized cellular states in niches provide one description of the compositional heterogeneity of cellular communities in the tissue. To quantify the distribution of cell states across niches, we constructed a cell-state count table, represented as a *n* x *m* matrix, where *n* is the number of niches (centered at every single cell) and *m* is the number of discretized cell states in the dataset. Then, we computed cell-state distributions at two resolutions, coarse cell-type and cellular compartment.

#### Compartment-specific niche expression matrices

The active molecular programs within a niche provide an additional description of the compositional and transcriptional heterogeneity of cellular communities. The first challenge in characterizing spatial patterns of transcriptional variation at the single-cell level is sparsity; each cell in our data expressed only ∼60 transcripts on average (**Table S2**). Moreover, genes in positive cells (cells with at least one mRNA count for that gene) had a median of 1.3 raw counts (interquartile range, 1.2–1.6). Sparsity persisted even when restricting our analysis to genes with known robust expression in a specific cell state. For instance, *Msn* had a median of 2 raw counts (interquartile range of 1–3) in *Msn*-positive cells. This contrasts with smFISH measurements of the same gene, which average 55 transcripts per progenitor-like cell (**Figures 1H, 2C**). Dropouts, or unobserved transcripts, dilute differences between cell subpopulations and limit the estimation of gene–gene covariance matrices^108^. We and others have developed approaches to overcome sparsity in dissociated data, including gene count imputation through signal sharing in kNN graphs^129^ and aggregation of gene counts in metacells—sets of cells that occupy the same transcriptional state, with minor residual variation between cells due to technical as opposed to biological sources^108,136^.

To overcome sparsity in Xenium spatial transcriptomics, we spatially aggregated mRNA counts, leveraging coarse cell-type annotations to obtain a compartment-specific summarized gene expression vector for each cellular niche. To do so, we first constructed a cell type x niche x gene count table, where each entry corresponded to the number of mRNA molecules in cells from a specific cell type in every niche in the dataset. We computed size factors as the total number of mRNA molecules in cells from the specified cell type in the niche, and used them to normalize our compartment-specific count matrix, followed by scaling by the median library size. Using this approach, each niche is represented by *m* independent count matrices, where *m* is the number of coarse cell types queried in the niche.

Spatial aggregation overcomes sparsity while preserving information about coarse cell types that express a given gene, facilitating the analysis of cell-type-specific expression differences between niches. On the other hand, aggregation loses single-cell resolution, and may mix heterogeneous states. However, our analyses at single-cell resolution showed concerted shifts in cell-state distributions within cell types when comparing niches dominated by gastric-like or progenitor-like cells (**Figure 3E**), suggesting that our spatial aggregation approach is well suited to capture the average changes in gene expression that accompany such shifts. In addition, while heterogeneity in cell density can introduce artifacts in these compartment-specific count matrices, as denser neighborhoods are more likely to have fewer collective dropouts in gene expression, our choice of *r* = *60 μm* mitigates this problem due to the large number of cells (> 100 on average) within each niche.

#### Ordering cells and niches by average gastric–progenitor DC

Oncogenic KRAS activation leads to the transcriptional diversification of the pancreatic epithelium, as well as profound tissue remodeling events linked to inflammatory responses^18–20^.

We hypothesized that the identity and local distribution of premalignant cells in tissue profoundly impact the morphological, cellular and molecular properties of their surrounding microenvironment. To investigate the relationship between cell-state changes in the premalignant epithelial cells and remodeling of the surrounding microenvironment, we integrated two viewpoints of our spatial data: (1) our quantification of niche features (see <u>Definition and identification of cellular niches</u> and subsequent sections) and (2) our characterization of gastric–progenitor cell-state continuum in premalignant cells (see <u>Diffusion</u> <u>component analysis in spatial data</u>). This strategy allowed us to connect the gradual transcriptional changes in premalignant cells with gradual changes in morphological, cellular and molecular properties of their niche.

We noted that gastric and progenitor-like cells within the same niche were at similar positions in the gastric–progenitor DC axis, suggesting spatial coordination in the adoption of divergent premalignant cell identities. The spatial coupling of premalignant heterogeneity allowed us to average the gastric-progenitor DC value of gastric or progenitor-like cells in the niche, resulting in a niche analog to the gastric-progenitor DC axis that we defined at the single cell level. In order to minimize variation due to averaging of small numbers of cells, we restricted our analyses to niches anchored at any cell with at least 10 gastric or progenitor-like cells in their niche. This newly constructed axis positioned each of the 2,772,533 niches in our data along a niche continuum linking the canonical gastric or progenitor niches.

### Summarization of niche features along the gastric–progenitor continuum

We used three complementary approaches to quantify how the cell-state composition of niches changes as a function of variation along the average gastric–progenitor DC, which we discretized into 100 uniform bins.

For our first approach, we quantified cell-type proportions in niches as a function of the DC axis bin, by computing the median and interquartile range of the relative frequencies of coarse cell types (**Figure S2G**), or cell states within a cell type (**Figure S2H**). For cell states, we included niches harboring at least 10 cells of the specified coarse cell type in the niche.

For our second approach, we disregarded discrete cell-state labels and instead visualized changes in the distribution of transcriptional states within cellular niches as a function of the DC axis bin (**Figure 3E**). Given niche matrix *Gr* × *n*, where *r* is the number of niches, *n* is the number of cells in the data, and *G_i_*_,*j*_ = 1 if *cell_j_* belongs to *niche_i_*, otherwise 0, we:

1. Extracted cell-state composition vectors for niches in each gastric–progenitor bin. We identified all niches that fall in a given bin in *G*, and extracted all cells that appear in these niches. This set of cells represents the neighborhood composition vector of niches in a specified bin along the gastric–progenitor axis.
2. Visualized cell-state densities in UMAP representations. We projected 2D cell density estimates onto a UMAP of tumor microenvironmental cells (**Figure 3E**) (see <u>Visualization of cell state density in two-dimensional representation</u>). As a reference, we visualized the density of gastric- or progenitor-like epithelial cells in the bin (**Figure 3E**). This approach provided a label-free visualization of changes in niche cell-state distributions coupled to changes in epithelial anchor-cell states.

For our third approach, we quantified gene expression changes in specific cellular compartments as a function of the DC axis bin (**Figure 3F**). Given compartment-specific niche expression matrix *X^c^* ^x^ *^n^* ^x^ *^m^*, where *c*, *n* and *m* represent the number of cell types, niches containing at least 10 gastric- or progenitor-like cells, and genes, respectively, and x*_s_*_,*i*,*j*_ is the average expression of *gene_j_* in cells of *type_s_*in *niche_i_* (see <u>Compartment-specific niche</u> <u>expression matrices</u>), we:

1. Standardized compartment-specific niche expression matrices. We log-transformed the niche expression matrix (*pseudocount* = 1) to stabilize variance and reduce the impact of outliers in downstream quantification. We standardized *X* by *z*-scoring log-transformed niche expression over all niches, for every cell type and gene pair. The resulting matrix represents cell-type-specific heterogeneity in gene expression across niches.
2. Summarized gene expression along niche bins. For a given cell type, bin and gene, we averaged our standardized compartment-specific niche expression matrix over all niches in the bin, resulting in a summarized niche expression matrix *X^c^* ^x^ *^b^* ^x^ *^m^*, where *b* is the total number of bins along the DC axis (*b* = 100). The resulting matrix represents the continuous gene expression shifts along the gastric–progenitor niche continuum in each cell type, analogous to the quantification of gene trends in pseudotime analysis^42,45^.
3. Visualized gene trends. To examine trends in niche gene expression, we first sorted genes based on how early their expression changed along the gastric–progenitor DC axis—when it reached a maximum value, minimum value, or changed in average *z*-score sign. We chose to visualize markers of gastric-like (e.g., *Anxa10, F5, Onecut2, Elf3*) and progenitor-like (e.g., *Msn, Hmga2, Vim*) cells, as well as myeloid (e.g., *Maf, Mab, Itgax*) and fibroblast (e.g., *Tnxb, Dpt, Postn, Tnc*) subpopulations that changed in abundance along the DC axis. In addition, we visualized genes that suggest shifts in signaling along the niche axis (e.g., *Oasl2*, *Irf7* for interferon signaling; *Dusp4, Dusp6, Dusp5* for MAPK signaling; *Mdm2, Cdkn1a* for p53 signaling). Lastly, we selected genes related to communication and wound healing that were upregulated in distinct cellular compartments of the progenitor niche.

These three approaches reveal changes in cell-state frequencies, cell-state densities, and compartment-specific gene expression as niches progress from gastric to progenitor-like epithelial anchor states, providing complementary viewpoints on the spatiotemporal dynamics of tissue remodeling during progenitor niche formation.

#### Robustness of results to niche size

To assess the robustness of gene trends along the G–P niche continuum with respect to niche radius, we repeated the procedure described above while systematically varying the niche radius between 15 μm and 200 μm, with a step size of 5 μm. For each radius, our procedure included (i) construction of compartment-specific niche expression matrices, (ii) binning niches along the G–P niche pseudotime, and (iii) computation of compartment-specific gene trends along the gastric-progenitor niche pseudotime. Next, for each gene in **Figure 3F**, we computed the Pearson correlation between gene trends computed for any given radius, with those identified using a 60-μm radius used throughout the main text (**Figure S2I**). Most comparisons showed strong correlation (Pearson correlation coefficient > 0.8), especially in the epithelial and fibroblast compartments. While these correlations were largely positive in myeloid cells, we identified stronger radius-specific differences in gene trends in this cellular compartment. On rare occasions, we found negative correlations between gene trends for specific radii, in relation to those identified using our reference radius. Two salient examples are the expression of *Csf1r* and *Adgre1*, two markers of macrophage maturation (**Figure S2I**). These results show that while most gene trends in our data are not strongly dependent on the size of the niche (up to a 200-μm radius), a systematic analysis varying niche size can reveal scale-dependent gene expression patterning in tissues.

#### Comparison of spatial niche expression with dissociated data

Our spatial analyses revealed a coupling between epithelial states and fibroblast and myeloid states within corresponding niches along the gastric–progenitor continuum (**Figure 3F**). These results were based on a handful of markers in our Xenium panel. To gain insights into broader programs across the full transcriptome, we reasoned that scRNA-seq data from our premalignant samples (see **Analysis of premalignant tumor microenvironment data**) should reflect similar axes of continuous variation, and thus searched for these axes in non-epithelial cell types in the niche microenvironment.

In the fibroblast compartment, we used our myofibroblast cell embeddings, as these are the major stromal components of the pancreatic parenchyma (see **Data S10** for embedding details). We applied diffusion component analysis to the scRNA-seq data (*k* = 30, adaptive Gaussian kernel with distance to tenth neighbor as bandwidth) to identify the major axis of transcriptional variation (DC1). We ordered cells along this axis and grouped them into 50 bins of equal cell number, then averaged *z*-scored gene expression over all cells in each bin to compute gene trends along DC1. Plotting genes in this orthogonal dataset (**Figure 3F**) revealed concordant trends: progressive downregulation of genes associated with inflammatory fibroblasts and universal fibroblast state (e.g., *Tnxb, Dpt, Cxcl12*), induction of activated myofibroblast genes near the progenitor-like cell extreme (e.g., *Tnc, Tgfb1*), and upregulation of Shh-related signatures in the middle of the axis (e.g., *Gli1, Ptch1, Ptch2*). By leveraging transcriptome-wide information, we confirmed the induction of additional fibroblast activation markers at the end of DC1, including *Acta2, Timp1* and *Spp1*, as well as tumor suppressor *Cdkn2a*, recently highlighted as a marker of senescent myofibroblasts that promote pancreatic cancer progression^53^ (**Figure S2J**).

In the myeloid compartment, we examined PhenoGraph clusters expressing *Maf* or *Itgax*, which each mark an extreme point in our niche continuum, and exist at intermediate levels along this axis in our spatial data. Notably, diffusion component analysis revealed that the *Maf*–*Itgax* dichotomy also captured the major axis of variation in scRNA-seq data (DC1). Projection of genes upregulated alongside *Itgax* and *Cd274*, markers of myeloid cells in the progenitor niche, revealed the engagement of a plethora of genes associated with immune suppressive and pro-fibrotic myeloid subsets (e.g., *Spp1, Arg1, Il1b, Fn1*) (**Figure S2J**). Taken together, this analysis allowed us to contextualize cell-state changes in the progenitor niche with transcription-wide information, some of which are known to mediate fibrotic responses and cancer progression.

#### Discretization of canonical gastric and progenitor niches

We found it useful to identify a single canonical progenitor and gastric-like niche, in order to simplify the computation of gene expression differences between these two communities (in wound-healing and communication gene programs, for example). We defined the canonical progenitor niche as the set of niches belonging to niche bins in which the median proportion of progenitor-like cells was higher than the median proportion of gastric-like cells. This corresponded to the top 22% of niches along the average gastric–progenitor DC niche axis. We defined the canonical gastric-like niche as the bottom 22% of niches ordered along the same axis. For simplicity, we term these sets gastric niche and progenitor niche.

#### Wound-healing signatures in the progenitor niche

We noticed that a plethora of genes with roles in wound healing processes (e.g., *Tnc* and *Postn* ECM remodeling, *Pdgfb* signaling, plasminogen processing) were upregulated in the progenitor niche across cellular compartments, motivating us to systematically test for wound-healing gene upregulation in our curated gene panel. We used all genes in our panel that appear in the Gene Ontology wound-healing response gene set (GO:0042060) as a wound healing signature, grouped niches by canonical gastric or progenitor label and by biological replicate, and averaged compartment-specific *z*-scored niche expression of these genes. Differences in signatures (**Figure 3G**) or individual signature genes (**Data S1.3R**) between gastric and progenitor niches were tested using a two-tailed Wilcoxon rank sum test per compartment. This strategy identified differences in wound healing responses between gastric and progenitor-niches and the top genes that drive this signature in each cellular compartment.

#### Communication module analysis

To identify potential channels of intercellular communication in the progenitor niche, we first identified ligands and receptors whose expression correlated with the position of a niche along G–P niche pseudotime. Specifically, we calculated the Spearman correlation between compartment-specific niche expression trends (see third approach in <u>Summarization of niche</u> <u>features along the gastric–progenitor continuum</u>) and their bin along the G–P niche pseudotime. We identified genes with a Spearman rank correlation > 0.9 as those that increase in the progenitor niche. This approach resulted in a set of candidate communication channels that satisfied both spatial co-occurrence in the progenitor niche, and upregulation in progenitor niches, relative to gastric niches (**Data S11**). To visualize concerted changes in the expression between cognate receptor–ligand pairs that may mediate heterotypic crosstalk in the progenitor niche (**Figures 5A–C** and **S3G**). Specifically, we plotted the expression of a ligand in a specified cellular compartment across different niches. Our data suggested that collective upregulation of multiple communication genes in spatially co-occurring cell states may contribute to the formation and stabilization of the progenitor niche.

### Tissue-level consequences of genetic and pharmacological perturbation

Our characterization of the injured premalignant pancreas revealed the tissue remodeling events that accompany the formation of progenitor niches at the morphological, cellular and molecular levels. These cancer-like wound-healing niches centered around progenitor-like cells, a subpopulation that simultaneously exhibited the highest engagement of tumor suppressive and oncogenic transcriptional signatures in the premalignant pancreas. Perturbation of *Kras* and *p53* signaling in epithelial cells profoundly impacted the abundance and state of this unique premalignant subpopulation: *p53* inactivation led to its expansion and adoption of advanced mesenchymal phenotypes (**Figure 7B–E**), whereas KRAS inhibition rapidly depleted this cellular state (**Figure 6B**). Next, we investigated how perturbation-induced changes in progenitor-like cells impacted non-epithelial cell types, particularly those associated with progenitor-like cells in the absence of additional interventions (e.g., *Tnc*+ myofibroblasts, *Itgax*+ macrophages/monocytes).

#### *In silico* dissection of pancreatic parenchyma

An immediate challenge for analysis was the sample-to-sample heterogeneity in tissue structures outside of the pancreatic parenchyma, including lymph nodes and adipose tissue. These structures were larger and more prevalent when derived from whole-pancreas FFPE blocks than from samples comprising a fraction of the pancreas (collected as part of a cohort with multiple types of readout). Given that interlobular spaces and cell-dense lymph nodes could introduce technical variability into downstream analyses (due to tissue size and dissection strategy), we used spatial information to focus only on cells directly associated with the pancreatic epithelium. This *in silico* dissection is analogous to the manual removal of mesenteric lymph nodes in our prior work with dissociated cells, which was critical to avoid having the sheer density of their immune cells dominate cell-state quantification. Our strategy was to:

1. Define the pancreatic parenchyma as the set of all epithelial cells, plus neighboring cells within a 60-μm radius. Any cell outside this set was ignored.
2. Computationally identify lymph nodes. Starting from centroids corresponding to lymph-node-associated immune cells (B, CD4 T and CD8 T cells), we constructed a spatial neighbor graph connecting all cells within 30 μm of each other. We identified lymph nodes as connected components in the spatial neighbor graph with > 250 cells, a parameter that we manually tuned to capture visually identified lymph nodes. Cells in lymph nodes were computationally ignored.

After *in silico* dissection, the premalignant pancreas epithelium corresponded to 84% of cells in the data. We used this subset of cells to analyze differential abundance of tumor microenvironment cells in response to perturbation.

#### Shifts in microenvironmental states upon oncogenic KRAS inhibition

Acute inhibition of oncogenic KRAS led to a dramatic reduction in progenitor-like cell abundance within 48 h of the first dose. To characterize the effect of this reduction on microenvironmental states and communication potential, we (1) quantified progenitor-like cell enrichment in the vicinity of different cell states, and (2) computed differential abundance with MiloR and interpreted results in light of the spatial association of enriched or depleted states with progenitor-like cells. We quantified and visualized changes in parenchyma cell-state abundance, excluding lymph nodes and interlobular spaces (see <u>In silico dissection of</u> <u>pancreatic parenchyma</u>).

We derived a quantitative scoring scheme to identify cellular states enriched near progenitor-like cells. For each niche anchored at a non-epithelial cell, we computed the fraction of progenitor-like or gastric-like cells relative to all epithelial cells within the niche (**Figure S5G**). Next, we defined ‘progenitor enrichment’ as the log-ratio of the progenitor-like cell fraction, relative to the fraction of progenitor-like cells in the dataset. We defined ‘gastric enrichment’ following the same logic for gastric-like cells.

To identify a set of microenvironment states associated with progenitor-like cells, we selected cells with a progenitor enrichment score > 2, a threshold beyond which the gastric enrichment score sharply decreased. Furthermore, we required a gastric enrichment score < 0, thereby selecting for niches specifically enriched for progenitor-like cells relative to other premalignant states. This analysis resulted in (1) a continuous feature for each microenvironment cell, describing the extent of enrichment of progenitor-like cells in their vicinity, and (2) a discrete set of cells tightly associated with progenitor-like cells.

Density estimates on tumor microenvironment UMAP embeddings revealed that cell niches enriched with progenitor-like cells were depleted in MRTX1133-treated samples (**Figure 6C**). To determine the significance of depletion, we applied the Milo algorithm^63^. Milo first constructs a set of tight and potentially overlapping cellular neighborhoods defined by transcriptional similarity on a kNN graph (makeNhoods function from miloR package, *prop* = 0.01, *k* = 30, *refined* = TRUE, *reduced_dims* = “PCA”, using pre-computed kNN and PCA), then tests for differences in cell counts between experimental conditions in each neighborhood using a generalized linear model with negative binomial residuals. Using Milo, we identified significantly enriched or depleted cell states upon acute oncogenic KRAS inhibition (SpatialFDR < 0.1).

To interpret differential abundance in light of defined cellular states, we annotated Milo neighborhoods by cell-state label, and found that 28% were composed of a single discrete label, representing pure cell states. For the remaining 72% of neighborhoods, we calculated the enrichment of the most frequent cell state relative to the second most frequent cell state. We conservatively excluded the 5% of Milo neighborhoods with lowest enrichment score (frequency of the most abundant cell state was less than 2.89-fold that of the second most abundant cell state), and labeled remaining neighborhoods by the predominant state. This strategy resulted in a set of transcriptional neighborhoods with a single cell-state label.

Lastly, we integrated spatial information into the interpretation of our differential abundance analysis. For each cell in a Milo neighborhood, we identified the cells in its niche, hereby termed Milo niche cells. Next, we calculated a progenitor enrichment score as the log ratio of the frequency of progenitor-like cells among Milo niche cells, relative to the frequency of progenitor-like cells in the dataset. We used this enrichment score to visualize the relationship between enrichment or depletion of neighborhoods, and their association with progenitor-like cells. Milo neighborhoods depleted upon acute oncogenic KRAS inhibition are strongly associated with progenitor-like cells in their niches (**Figure 6D**).

Tissue-wide cell state proportions in KC^shp53^ and KC^shCtrl^ samples To assess the tissue-wide consequences of *p53* loss in the injured pancreas, we quantified the frequency of cell-state proportions relative to their coarse cell type in each biological replicate. We restricted our analysis to the pancreatic parenchyma, to avoid potential confounding by large histological structures such as lymph nodes (see <u>In silico dissection of pancreatic</u> <u>parenchyma</u>). Since tissues collected in KCshp53 showed substantial heterogeneity, we leveraged the continuity and heterogeneity in abundance of progenitor-like cells to identify microenvironmental states that increase in abundance in concert with this epithelial subpopulation. **Figure S7A** shows the three of the microenvironmental states that most strongly correlate with the abundance of progenitor-like cells at the tissue level.

#### Molecular programs in *Itgax*+/*Cd274^high^* cells

To better understand the molecular properties of *Itgax*+/*Cd274^high^* macrophages/monocytes that accumulated upon *p53* knockdown in premalignant pancreatic epithelial cells, we leveraged our myeloid-specific embedding of dissociated single cell data from KC^ctrl^ or KC^shp53^ mice three weeks post injury (see <u>Analysis of premalignant tumor microenvironment data</u>). First, we identified *Cd163*, *Maf*, *Itgax* and *Cd274* as four landmark genes that described progressive gene expression changes along the average gastric-progenitor niche axis in myeloid cells.

Plotting and visualization of these four genes in UMAP embeddings of dissociated myeloid cells revealed that they captured gradual shifts in cell myeloid cell states in these data, consistent with spatial transcriptomics analyses (**Figure S7D,E**). Moreover, we observed accumulation of Itgax+/Cd274^high^ cells in KC^shp53^ samples, corroborating our findings in spatial data, and providing the opportunity to study this subpopulation in greater depth. We used the wald test in the diffxpy package (v0.7.4, https://github.com/theislab/diffxpy?tab=readme-ov-file) and library size as numeric covariates. We identified upregulated genes using the following thresholds: *qval* < 0.05, log_2_ fold-change > 1. To highlight the top upregulated and downregulated genes from this analysis, we identified genes with mean_expression > 1, thus reporting changes in genes that are robustly expressed in at least one of the subpopulations (**Figure S7E**).

#### Quantitative relationship between cell state frequencies in cellular niches

Knockdown of *p53* in premalignant epithelial cells led to the accumulation of progenitor-like cells with advanced mesenchymal phenotypes (**Figure 7B–F**). These cells formed large tissue domains that could encompass entire pancreatic lobes (**Figure S7B,C**), and were accompanied by the accumulation of *Itgax*+/*Cd274^high^* (PD-L1^high^) macrophages/monocytes. Remarkably, the same tissue could contain regions devoid of progenitor-like cells and their associated microenvironments. This heterogeneity suggested that the local density of progenitor-like cells could determine properties of their microenvironment. To quantify such relationships, we determined cell-state frequencies relative to their coarse cellular compartment in cellular niches of different sizes. We log-transformed cell state frequencies, setting as pseudount the minimum non-zero value in each frequency vector. Next, we computed the joint distribution of the log frequency of progenitor-like cells (relative to epithelial cells in the niche) and the log frequency of *Itgax*+/*Cd274^high^*cells (relative to myeloid cells in the niche). To visualize the cell-state relationship in a manner that is agnostic to the marginal distribution of the progenitor-like state frequency, we computed the distribution of *Itgax*+/*Cd274^high^*cell frequencies conditioned on the frequency of progenitor-like cells in the niche, following the strategy outlined in DREVI^158^. Our analysis revealed that changes in the local density of progenitor-like cells are statistically related to changes in the frequency of their associated *Itgax*+/*Cd274^high^*cells, suggesting also the presence of non-linearities in the quantitative relationship between the relative abundance of these two cell states (**Figure S7F**).

#### Detection of significant cell–cell interactions in single-cell dissociated data

To identify emergent cell–cell interactions facilitated by tissue remodeling upon *p53* loss in the premalignant pancreas, we leveraged our single-cell dissociated dataset, and our cell–cell communication inference method Calligraphy. This allowed us to gain transcriptome-wide information of cell–cell communication potential, while leveraging the knowledge regarding the subpopulations that become enriched in the progenitor niche upon *p53* knockdown, as revealed by our spatial transcriptomic profiles. Specifically, we used single-cell transcriptomes from dissociated and sorted premalignant epithelial cells derived from KC^shp53^ (*n* = 5) and KC^shRen^ (*n* = 4) mice 3 weeks post injury. Myeloid and fibroblast data comes from single-cell transcriptomes derived from FFPE blocks and sequenced using the 10x Genomics Flex approach (*n* = 2 for control and shp53 samples). For fibroblasts, we subset the data to include only the myofibroblast cluster, as these are the most abundant fibroblasts in the pancreatic parenchyma. Inclusion of other fibroblast populations (e.g., apCAFs or iCAFs) make it harder to identify smaller subpopulations in the myCAF compartment because differences between broad populations dominate the covariance structure of the data.

Our algorithm first identifies communication modules—groups of communication genes with tight correlations, that define specific cell subpopulations in single-cell data. We useimputation to enhance gene–gene correlation estimates, as dropouts dilute the signal from correlated gene pairs. We imputed counts on each cellular compartment independently using MAGIC^129^ (affinity matrix constructed on a kNN with *k* = 30, adaptive kernel width with *k* = 10, *t* = 2). For the epithelial compartment, we imputed shRen and shp53 samples separately to avoid smoothing out potential genotype-specific features, using the same PC space for constructing the kNN graph in either genotype. Following genotype-specific imputation, we concatenated the imputed matrices.

We next used the CellChat^159^ ligand–receptor database to define potential interacting communication gene pairs. Although the database specified more than pairwise interactions (e.g., one complex interacting with one ligand), we transformed this into a binary interaction network (for a receptor complex, listing all possible pairwise interactions with a ligand or complex of ligands).

In undertaking feature selection, we aimed to identify communication genes that are robustly expressed in each cellular compartment. We aggregated cells by PhenoGraph cluster, and computed the mean imputed expression of each gene in each cluster. Next, we identified the maximum cluster-averaged expression for each gene and computed the distribution. Using triangle thresholding, we identified a cutoff for expressed vs non-expressed genes in each compartment. In addition, we noticed that despite ambient correction in our datasets, there was still low expression of specific communication genes that was suggestive of residual contamination of gene counts between cell types (e.g., *Cdh1* in myeloid cells, or *Pecam1* in epithelial cells). We censored these “cell-type” marker communication genes from cell types that are not expected to express them. Lastly, we restricted analysis to paracrine interactions due to the promiscuity of ECM–integrin interactions. Because our shuffling strategy in Calligraphy does not control for the connectivity of communication genes, inclusion of these genes obscure the statistical significance of other interactions driven by communication genes with lower connectivity.

To identify communication modules, we followed the original approach from Burdziak, Alonso-Curbelo et al.^7^. First, we constructed a gene–gene graph based on the communication gene correlation matrix. We tuned the correlation threshold for each cellular compartment to make sure that we captured the subpopulations that we identified in spatial data. Next, we used Jaccard similarity to remove edges that are not well connected to the rest of the nodes in a clique (Jaccard similarity index < 0.05), and to include edges that are below the correlation threshold for any given gene pair, but that are well connected within a clique (Jaccard similarity index > 0.95). Lastly, we used OSLOM for graph-based community detection, allowing genes to participate in multiple cliques. In the case of myeloid cells, we identified a communication module that expressed high levels of fibroblast markers. We removed this module from further inference, as we reasoned that this small group of cells most likely represented doublets or rare fibroblasts misclassified as myeloid cells during data preprocessing (**Data S1.3S–U**). **Data S5** specifies the communication modules that we identified in different compartments.

Next, we identified statistically significant interactions. For each pair of cliques, we computed the number of ligand–receptor interactions (directional approach). We then used the shuffling strategy defined by Burdziak*, Alonso-Curbelo* et al.^7^ to construct a null distribution of the expected number of ligand–receptor interactions for each pair of communication modules. At each iteration, we shuffled the gene-module identity, preserving the number of genes assigned to each module. By shuffling module identity 5000 times, we identified pairs of modules with an observed number of predicted ligand-receptor interactions that was greater than the expectation by chance. **Data S5** specifies significant inter-module interactions.

We visualized the results using a circular layout with the networkx python package (**Figure S7G**). Our graph shows only modules with at least one significant intermodule interaction.

